# Known unknowns and model selection in ecological evidence synthesis

**DOI:** 10.1101/2024.12.18.629303

**Authors:** Shane A. Blowes

## Abstract

Quantitative evidence synthesis is a prominent path towards generality in ecology. Generality is typically discussed in terms of central tendencies, such as an average effect across a compilation of studies, and the role of heterogeneity for assessing generality is not as well developed. Heterogeneity examines the transferability of ecological effects across contexts, though between-study variance is typically assumed as constant (i.e., homoscedastic). Here, I use two case studies to show how relaxing the assumption of homoscedasticity and cross validation can combine to further the goals of evidence syntheses. First, I examine scale-dependent heterogeneity for a meta-analysis of plant native-exotic species richness relationships, quantifying the relationships among unexplained effect size variation, spatial grain and extent. Second, I examine relationships among patch size, study-level covariates and unexplained variation in species richness using a database of fragmentation studies. Heteroscedastic models quantify where effects can be transferred with more or less certainty, and provide new descriptions of transferability for both case studies. Cross validation can be applied to a single or multiple models, adapted to either the goal of intervention or generalization, and showed that assuming homoscedasticity can limit transferability for both case studies.

## Introduction

Quantitative evidence synthesis aims for general insights into the direction, magnitude and variability of ecological effects (Gurevitch et al. 2018, Fox 2019, Spake et al. 2022). Two common forms of quantitative synthesis in ecology are meta-analysis (i.e., analyses of effect sizes collated or calculated from existing studies), and analyses of primary data compilations (Mengersen et al. 2013, Spake et al. 2022). For both approaches, generalities typically come in the form of central tendencies, such as an average effect size across studies. While both approaches frequently quantify the heterogeneity of effect sizes across studies to examine how consistent ecological effects are (Lau et al. 2013, Senior et al. 2016, Gurevitch et al. 2018, Spake et al. 2022, Nakagawa et al. 2023a), the role of heterogeneity for assessing generality is less well developed.

Identifying sources of heterogeneity can be critical for understanding a given phenomenon (Lau et al. 2013, Gurevitch et al. 2018), and contributes to the goal of generality by determining how transferable effect sizes are across different contexts (Fox 2019). However, effect size variance is typically assumed to be constant (i.e., homoscedastic; Williams et al. 2021, Viechtbauer & Lopez-Lopez 2022, Nakagawa et al. 2025), equating to an assumption that included studies sample a single statistical population (Williams et al. 2021). Moreover, assuming homoscedasticity largely limits questions to those focused on the magnitude of responses, whereas many questions would benefit from explicit (quantitative) examinations of variability (Cleasby & Nakagawa 2011, Nakagawa et al. 2025). Location-scale (mean-variance) models that can model heteroscedasticity have been described for both meta-analytic models (Williams et al. 2021, Viechtbauer & Lopez-Lopez 2022, Nakagawa et al. 2025), and the multi-level or mixed effect models frequently used in syntheses of primary data (Lee & Nelder 1996, 2006, Pinherio & Bates 2000, Zuur et al. 2009), but their use in quantitative evidence synthesis remains relatively rare (Nakagawa et al. 2025).

Specifying models for unexplained variation can provide direct assessments of the limits to transferability. For example, covariates associated with unexplained variation could help delineate contexts where quantitative evidence syntheses generalize with more or less uncertainty. If different groups, such as taxon groups or geographic regions are associated with more or less unexplained variation, researchers could better communicate where effect sizes can be transferred with more or less confidence. Such quantitative assessments of unexplained variation can help identify promising directions for both empirical and theoretical research (Viechtbauer & Lopez-Lopez 2022).

To evaluate and/or compare homoscedastic and heteroscedastic models for evidence synthesis, some form of quantitative model testing is needed. Cross validation is a broadly applicable method used to evaluate model predictive performance (Hastie et al 2009), and has the potential for direct assessments of transferability in an evidence synthesis context (Spake et al. 2022). Out-of-sample model predictions are key component of transferability (Spake et al. 2022), and cross validation out-of-sample tests can be useful for a single model (e.g., does the model make more or less accurate predictions for different geographic regions or taxonomic groups?), as well as for comparing out-of-sample predictions between different models (Vehtari 2025). Moreover, cross validation is highly flexible, and data splitting (Hastie et al 2009) can be designed to compare the ability of models to make different types of predictions (e.g., Merkle et al. 2019, Yates et al. 2022). For example, an evidence synthesis with a goal of intervention assessment (Guretvitch et al. 2018) might be best evaluated by testing within sample predictions, assuming included studies are a probability sample (Boyd et al. 2023) of the target group or population for the intervention. Whereas to meet the goal of generality (Gurevitch et al. 2018, Fox 2019), models can be more usefully evaluated by their ability to predict data for a new study (or studies) outside of the data used to train the model.

To show how location-scale models and cross validation can combine to further the goals of quantitative evidence synthesis in ecology, I present two case studies. Before the details of each case study, I describe how cross validation is used in the case studies, briefly introduce software capable of fitting location-scale models, and describe the model fitting workflow used to validate all models in each case study. The first case study extends a meta-analysis of spatial scale-dependence in plant native-exotic species richness relationships (Peng et al. 2019) to quantify relationships among unexplained variation, grain size and spatial extent. The second case study uses a primary data compilation of habitat fragment diversity studies (Chase et al. 2019). I focus on the relationship between fragment size and local (i.e., patch-scale) species richness, and examine whether residual variation is related to fragment size and other study-level covariates. For both case studies, cross validation shows that the assumption of constant unexplained variation limits model predictive performance, especially when making predictions to new studies (i.e., out-of-sample predictions).

### Cross validation for model comparison in evidence synthesis

Cross validation uses data splitting techniques to test model predictive performance. Models are fit to a ‘training’ data set and assessed on their ability to predict the ‘test’ data set (Hastie et al 2009). Cross-validation also requires a function to quantify predictive performance, and here I used pointwise *expected log predictive density* (*elpd*; see Yates et al. 2022 for further discussion of other common loss functions). Additionally, to minimize an inherent bias to support overfit models, I used the modified one-standard-error rule (Yates et al. 2021) that aims to select the least complex model that is comparable to the best scoring model.

To show how cross validation could be used to further different goals of evidence synthesis (Gurevitch et al. 2018), I use cross validation to evaluate model performance for two different types of predictions: i) within-sample or conditional predictions that examine model performance when making predictions to new data within existing studies, and (ii) out-of-sample or marginal predictions that assess predictive performance to new studies (Merkle et al. 2019, Yates et al. 2022).

To assess conditional (within sample) predictions, approximate or exact leave-one-out (loo) cross validation is considered the gold standard (Yates et al. 2022). However, approximate loo diagnostics showed many influential observations for the models in both case studies presented here, which reduces the reliability of performance estimates (Vehtari et al. 2017); and, because exact loo can be computationally expensive for large data compilations (due to the need to refit the model as many times as there are data points), I approximated loo using *k*-fold cross validation. Importantly for cross validation, most quantitative evidence syntheses, including the case studies presented here, have dependencies in the data structure. For example, studies might contribute multiple effect sizes to a meta-analysis, or the data might have spatial or other dependencies. Block cross validation (i.e., including structure in the sub-setting process when portioning training and test data; Roberts et al 2017, Yates et al. 2022) is typically recommended when the data themselves are structured. Accordingly, I use a stratified *k*-fold approximation of loo for both case studies. Stratified *k*-fold cross validation is appropriate for the hierarchical data typical of evidence syntheses because it balances data splitting among subgroups when creating the *k* blocks of data to which models are refit, ensuring that the relative group frequencies (here, the data coming from included studies) are maintained. Similar to loo, *k*-fold cross validation is focused on making predictions to new data points within existing studies conditional on the model parameters, and I used *k* = 10 folds in both case studies to assess within sample predictive performance.

To assess out-of-sample predictive performance, I used leave-one-group-out cross validation. Individual studies were removed one at a time, and models assessed on their ability to predict the data in the held-out study.

### Model fitting workflow

Location-scale statistical models allowing predictor variables for both the mean (location) and residual variation (scale) are well studied (Lee & Nelder 1996, 2006, Pinheiro & Bates 2000, Zuur et al. 2009), and have been described for meta-analytic models (Williams et al. 2021, Rodriguez et al. 2022, Viechtbauer & Lopez-Lopez 2022, Nakagawa et al. 2025). Many R packages commonly used by ecologists allow for location-scale models to be fit using either frequentist (e.g., *nlme*, Pinheiro & Bates 2024; *glmmTMB*, Brooks et al. 2017, *metafor*, Viechtbauer 2010) or Bayesian methods (e.g., *brms*, Bürkner 2017); note that (as far as I am aware) frequentist packages are currently limited to fitting non-varying (i.e., fixed) parameters for the scale component. Due to the availability of tools for diagnosing model pathologies (Betancourt 2016, Monnahan et al. 2017), cross validation (Vehtari et al. 2017) and model calibration (Modrák et al. 2023), as well as the ability to fit varying (random) parameters for the scale component, I fit models using the Hamiltonian Monte Carlo (HMC) sampler Stan (Carpenter et al. 2017). Models were coded using the ‘brms’ package (Bürkner 2017). Code (and data) for all analyses is available: https://github.com/sablowes/heterogeneity-evidence-synthesis (and will be archived following acceptance).

To robustly build increasingly complex heteroscedastic models I followed a workflow that used simulations to check the calibration of all models (Gelman et al. 2020, Talts et al. 2020, Modrák et al. 2023, Säilynoja et al. 2025). To examine whether inference with a particular model is feasible conditional on the observed data, I focus on calibration in the parameter space of the empirical posterior (i.e., posterior simulation-based calibration, Säilynoja1 et al. 2025). Briefly, the model is first fit to the empirical (i.e., observed) data. Then, the same model with priors informed by the fit to the empirical data is used to simulate many new data sets (with the same size, shape and structure as the empirical data), and the model is fit to each of the simulated data sets. Finally, plots examining model calibration (e.g., are known parameters of interest recovered with reasonable coverage?) are inspected. Appendix S1 presents the simulation-based model calibrations for case study one; the calibration for models in case study two are in Appendix S2.

#### Case study one: Scale-dependent uncertainty in plant native-exotic richness relationships

Communities with more species are often thought to be more resistant to invasion by exotic (or non-native) species than communities with fewer species (Elton 1958). However, the spatial scale-dependence of biodiversity can complicate overly simple interpretations of this idea. Negative relationships between the numbers of native and non-native species are thought more likely at small scales, with a switch to positive relationships expected at larger spatial scales (Levine 2000). Such scale-dependence has been linked to niche opportunities (Shea & Chesson 2002). At relatively small scales, more species rich native communities leave fewer niche opportunities for exotics to invade. As spatial scale increases, so too does heterogeneity in resources, natural enemies and the physical environment, which creates greater niche availability for exotic species to become established (Shea & Chesson 2002).

Peng et al. (2019) synthesized evidence for relationships between spatial scale (grain and extent) and the correlation of native and exotic species richness using multilevel meta-regressions across 101 observational studies, encompassing 204 effect sizes. On average, Peng et al. (2019) found native and exotic richness positively correlated, and that positive correlations between native and exotic species richness got stronger with increasing (log) grain size (i.e., the size of the sampling unit for the observations, e.g., 1m2 quadrat). All models fit by Peng et al. (2019) assumed constant between-study variance, and adjusted for the non-independence of multiple effect sizes coming from some studies with (nested) multilevel varying (random) intercepts. Here I relax the assumption that between study heterogeneity is constant, and examine whether unexplained variation in the effect size is related to either grain size or extent. I compare how all models perform for within and out-of-sample predictions using cross validation.

Peng et al. (2019) transformed all correlations to Pearson product moment correlation coefficients (*r*), and calculated effect sizes using the Fisher *z*-transformation: 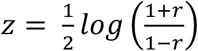, with sampling (within-case) variance estimated as 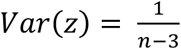 (denoted 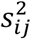 below); *log* refers to the natural logarithm (as it does hereafter), and *n* is the sample size (i.e., number of native-exotic species pairs; effect size data sourced from Peng et al. 2019). I start by reproducing the main result from Peng et al. (2019) using a multilevel linear meta-regression model for grain size that assumes the effect sizes, *z*_*ij*_, are normally distributed with known within-case variance 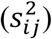, and constant between-study variance (*τ*^2^), which can be expressed as:

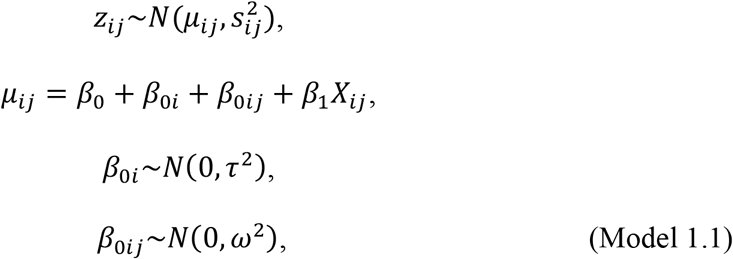

where cases (*j*) are nested within studies (*i*), and have among-case (within study) variance, *ω*^2^; between-study heterogeneity, *β*_0*i*_, has constant variance *τ*^2^, and varies around the overall linear relationship for the location (*μ*_*ij*_) with intercept, *β*_0_, slope *β*_1_, and predictor *X*_*i*_ (here the natural logarithm of grain size in study *i*, case *j*). Given the relationship between effect size and grain size (Peng et al. 2019), I retain (log) grain size as a predictor in all subsequent models.

Peng et al. (2019) observed that variation around the average (positive) relationship between grain size and effect sizes decreased with increasing grain size. That is, residual heteroscedasticity decreased as a function of grain size, though they did not explicitly model this relationship. Recall 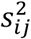 is the known (so called sampling) variance of the effect size estimate (and is not estimated from the data in any of the homoscedastic or heteroscedastic models presented here); the first heteroscedastic model introduces a new parameter 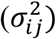 for the scale component of the model to be estimate from the data, and I model the (log) standard deviation of this parameter (σ) as function of grain size:

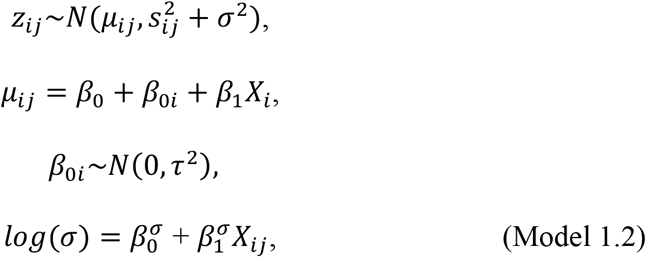

where *X*_*ij*_ is grain size on a log-scale for the *j*th case in study *i*. This model did not converge when the case-level random intercept (*ω*, see Model 1.1) was included, likely due to identification problems associated with multiple parameters (i.e., σ and *ω*) describing the same level of variation in the data.

To model extent, Peng et al. (2019) used discrete bins with a range of one order of magnitude in square kilometers, i.e., (0, 10), [10, 100), [10^2^, 10^3^), …, [10^6^, >10^6^). Multiple meta-regression models were fit to examine for an effect of spatial extent on effect sizes. Peng et al. (2019) did not report a strong influence of spatial extent on effect sizes, though they did describe an interaction between extent and grain. Here, I examine whether unexplained variation (i.e., after adjusting for the relationship between effect size and grain size) was related to extent using a model where residual variation was a function of spatial extent. I fit extent as a categorical predictor of the residual variation 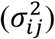, i.e., the same model for the scale as 1.2, but without the intercept 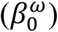, and extent categories coded as indicator variables for the predictor, *X*_ij_ [Model 1.3], instead of the continuous predictor fitted for grain in model 1.2.

Finally, I fit a model that allowed between study variation in the scale component. Models with varying (or random) parameters specified for both the location and the scale components were first introduced as double hierarchical generalized linear models (Lee & Nelder 1996, 2006), and they can be estimated with or without correlations among the varying parameters (see e.g., case study two below for an example with co-varying parameters for the location and scale). The model was again fit with (log) grain size as a predictor (*X*_*i*_) for the location (mean, *μ*_*ij*_) effect size, and varying parameters for the scale were estimated independently of the location:

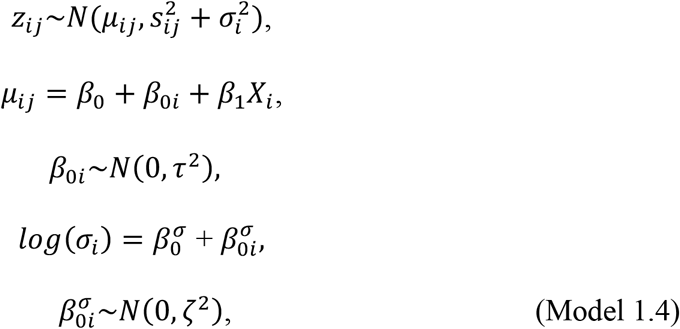

where 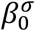 is the average residual variation (on a log-scale), and 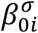 is a normally distributed study-level departure from the average intercept for the scale parameter with zero mean and *ζ* standard deviation.

#### Case study one: results

All models reproduced the observed data well, and simulation-based calibration showed all models had reasonable coverage for the parameters of interest (Appendix S2: Figs. S1-S16).

Model selection identified the same model (1.4) as best for making predictions within existing studies (Fig. 1a) and to new studies (Fig. 1b), though subsequent ordering among the other models depended on predictive task (Fig. 1). All models produced similar estimates for the parameters they shared (i.e., β_0_, β_1_, τ; Appendix S1: Fig. S17). The constant heterogeneity (i.e., homoscedastic) meta-regression (model 1.1) performed worst for making both within (Fig. 1a), and out-of-sample predictions (Fig. 1b). Model 1.2 quantified the average decrease in residual variation with increasing grain size (as was observed qualitatively by Peng et al. 2019; 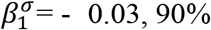 credible interval: -0.09 *–* 0.02), and showed uncertainty surrounding the relationship was greatest for the smallest and largest grains (Fig. 1c). After adjusting for the positive relationship between average effect sizes (i.e., the location) and grain size, model 1.3 showed unexplained variation was on average greatest at the smallest extents (i.e., the total geographic area covered by the samples; Fig. 1d).

**Figure 1:**
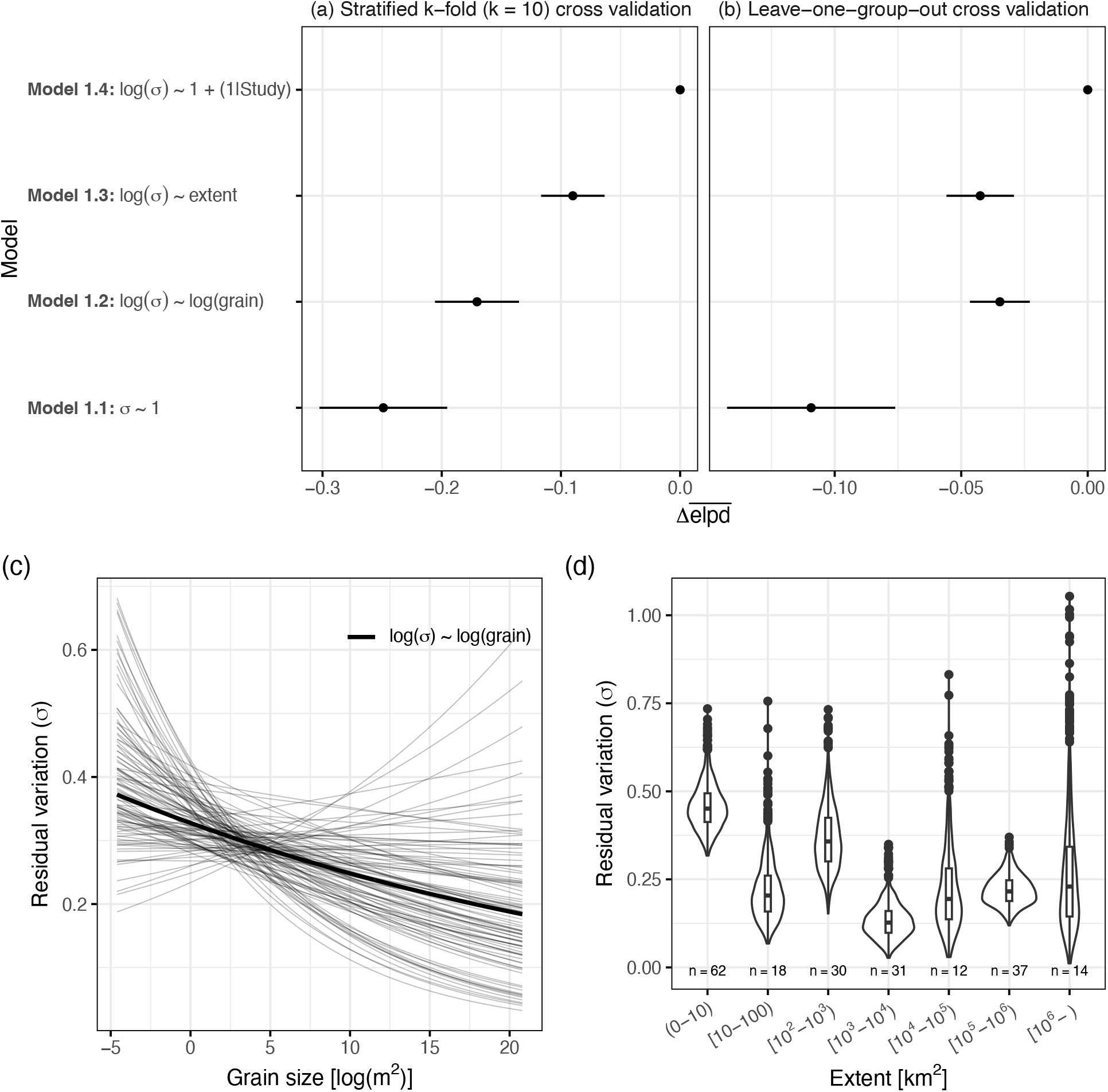
Heterogeneous heterogeneity outperforms constant heterogeneity for predicting plant native-exotic species richness relationships. Model selection for predictions (a) within existing studies using stratified k-fold cross-validation, and (b) for new studies using leave-one-group-out cross validation. Scale-dependent unexplained variation for (c) grain, and (d) extent. Thick lines on (c) show median expectation for grain-size dependent unexplained variation from model 1.2 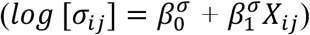; thin lines show 100 draws from the posterior distribution to visualise uncertainty. Violin plots (d) show the distribution of 1000 draws of the posterior distribution of unexplained variation (σ) for each of the extent categories; boxplots show the median (bar), 25% and 75% quantiles (box), whiskers show 1.5 times the interquartile range, points show observations beyond 1.5 times the interquartile range; n = number of effect sizes for each extent category (cases).

### Case study two: Variation in patch-scale effects of ecosystem decay

Habitat loss is a major driver of biodiversity loss (Diaz et al. 2019). As habitat is removed from landscapes, biodiversity reductions in remaining habitat fragments can be thought of as arising from one of two processes (Chase et al. 2020). First, because fewer individuals and species can live in smaller habitat fragments, the species found small habitat fragments might simply be a random (or passive) sample of those found in large habitat fragments. Alternatively, if smaller habitat fragments negatively impact demographic rates relative to larger habitat fragments, then ecosystem decay might result in greater diversity declines compared to those expected from a solely passive sampling process (Chase et al. 2020).

To test these competing hypotheses for (patch-scale) diversity fragment size relationships, Chase et al. (2019) compiled assemblage data (counts of individuals of each species) in effort-standardized (or standardizable) samples across habitat fragments of different sizes within landscapes. Initial analyses of these data showed that across 123 studies and 1509 habitat fragments, neither the number of individuals nor the species found in small fragments were simply a random sample of those in large habitat fragments (Chase et al. 2020). Instead, altered demographic rates in smaller habitat fragments, due, e.g., to edge effects, reduced dispersal or demographic stochasticity (collectively referred to as ecosystem decay), resulted on average in fewer individuals, fewer species, and less even communities than expected from a passive sample of diversity in larger fragments (Chase et al. 2020). Here, I revisit this analysis to quantify heteroscedasticity, and examine how modelling heteroscedasticity impacts within and out-of-sample model predictions. For simplicity, I focus on species richness, and start by refitting the same multilevel model used by Chase et al. (2020) that assumed residual variation was constant across all of the studies. I then fit a series of models of increasing complexity that relax the assumption of homoscedasticity.

The homoscedastic model fit by Chase et al. (2020) to effort-standardized species richness (*S*_*ij*_) in fragment *j* from study *i* took the form:

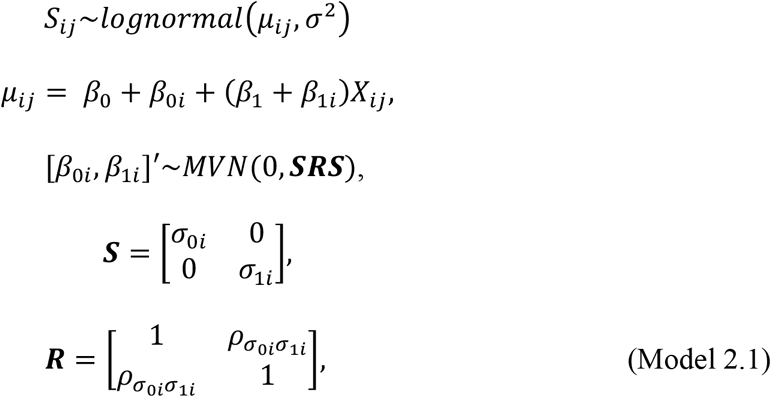

where *X*_*ij*_ is the fragment size on a (natural) log-scale, which was centered by subtracting the overall mean from each observation before modelling; *β*_0i_ and *β*_1i_ are study-level departures from the overall intercept (*β*_0_) and slope (*β*_1_), respectively, drawn from a multivariate normal (*MVN*) distribution with standard deviations, *σ*_0*i*_ and *σ*_1*i*_ that estimated correlations (*ρ* in the ***R*** matrix) between the varying intercepts and slopes for the location (i.e., mean richness, *μ*_*ij*_).

Next, I consider heteroscedastic extensions of increasing complexity; all extensions take the form of double hierarchical generalized linear models (i.e., varying parameters for both the location and scale; Lee & Nelder 1996, 2006). The first heteroscedastic model is motivated similarly to the varying study-level intercepts and slopes for the location (*μ*_*ij*_). Assuming average richness and its relationship with fragment size varies among studies gets us the benefit of adaptive regularization (i.e., shrinkage, McElreath 2020) when estimating the location. And we can get the same types of benefits when estimating parameters for the scale (i.e., residual variation). Model 2.2 specified varying study-level residuals estimated independently of other varying study-level parameters for the location (mean):

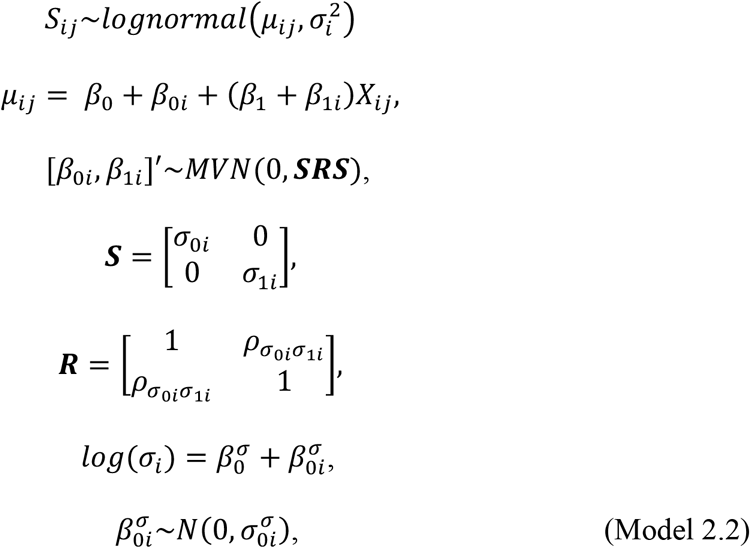

where 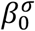 is the overall average of residual variation (on a log-scale), and 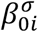 are study-level departures (for the scale or residual variation) drawn from a normal distribution with zero mean and standard deviation 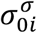.

Next, I extend this model to include (log) fragment size as a predictor of residual variation (i.e., scale). Correlations between varying (study-level) intercepts and slopes of both the location and the scale are estimated, but varying parameters for the location (*μ*) are estimated independently of varying parameters for the scale (σ):

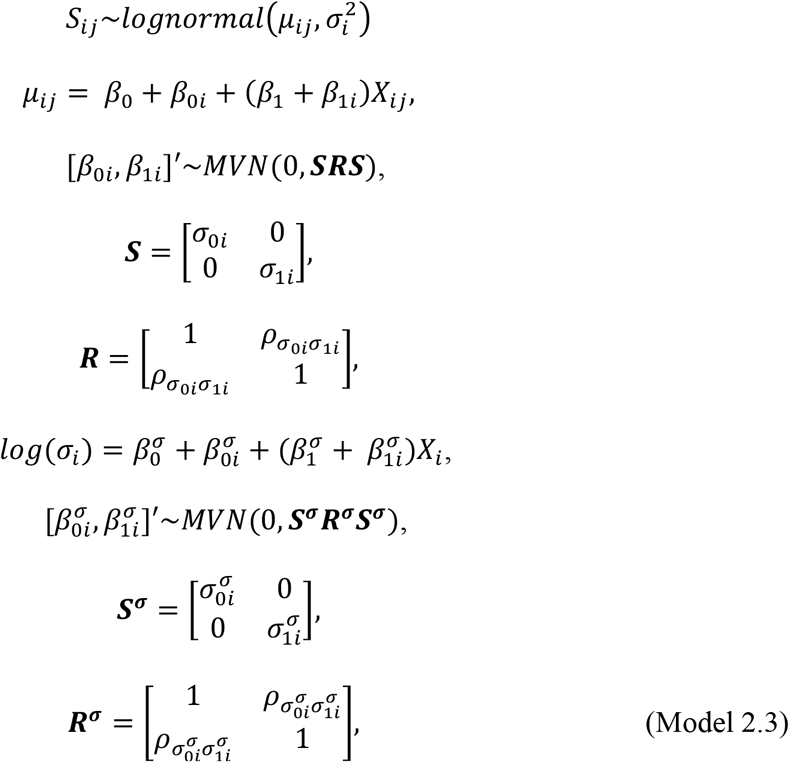

where 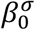 is the overall average residual variation, and 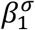 is the overall average slope of residual variation with fragment size; 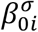 and 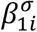 are the varying study-level departures from the intercept and slope, respectively, and were drawn from a multivariate normal distribution with zero mean and standard deviation 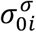 and 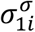, with correlations estimated in matrix ***R***^***σ***^.

Finally, I estimate models with study-level residual variation both with and without fragment size as a predictor that allow for correlations between varying (study-level) parameters for the location (μ) and scale (σ). The model without fragment size is:

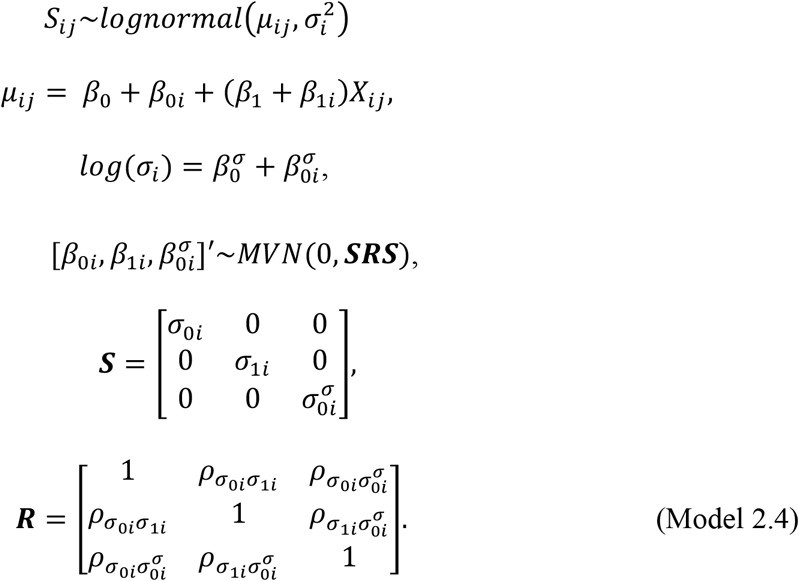

The model with fragment size as a predictor of both the mean (location) and residuals (scale) that allows for correlations among the varying study-level parameters is:

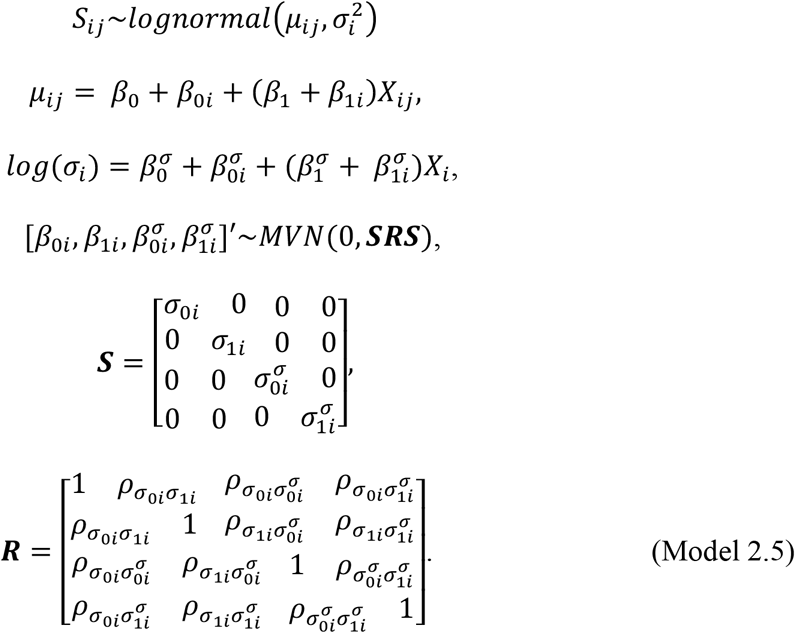

#### Case study two: results

All models were able to reproduce the observed data well, and simulation-based calibration showed that all models had reasonable coverage for parameters of interest (Appendix S2: Fig. S1-S20).

Modelling heteroscedasticity did not qualitatively impact the support for the ecosystem decay hypothesis for species richness (Appendix S2: Fig. S21; Chase et al. 2020). However, heteroscedasticity models outperformed the model with constant residual variation for making predictions of species richness for new fragments in existing studies (Fig. 2a), and to entire new studies (Fig. 2b), though different models were favored for within versus out-of-sample predictions. For making predictions to new data in existing studies, cross validation supported the most complex model (model 2.5; Fig. 2a). This model shows residual variation was a decreasing function of fragment size (Fig. 2c), and estimated correlations between varying study-level parameters for the location and scale (i.e., mean and residual variation; Appendix S2: Fig. S22). For example, study-level fragment size slopes for the mean effect and residual variation were negatively correlated (Appendix S3: Fig. S22a), meaning that the strongest ecosystem decay effects were associated with the strongest decline in residual variation with increasing fragment size (Appendix S3: Fig. S23).

**Figure 2:**
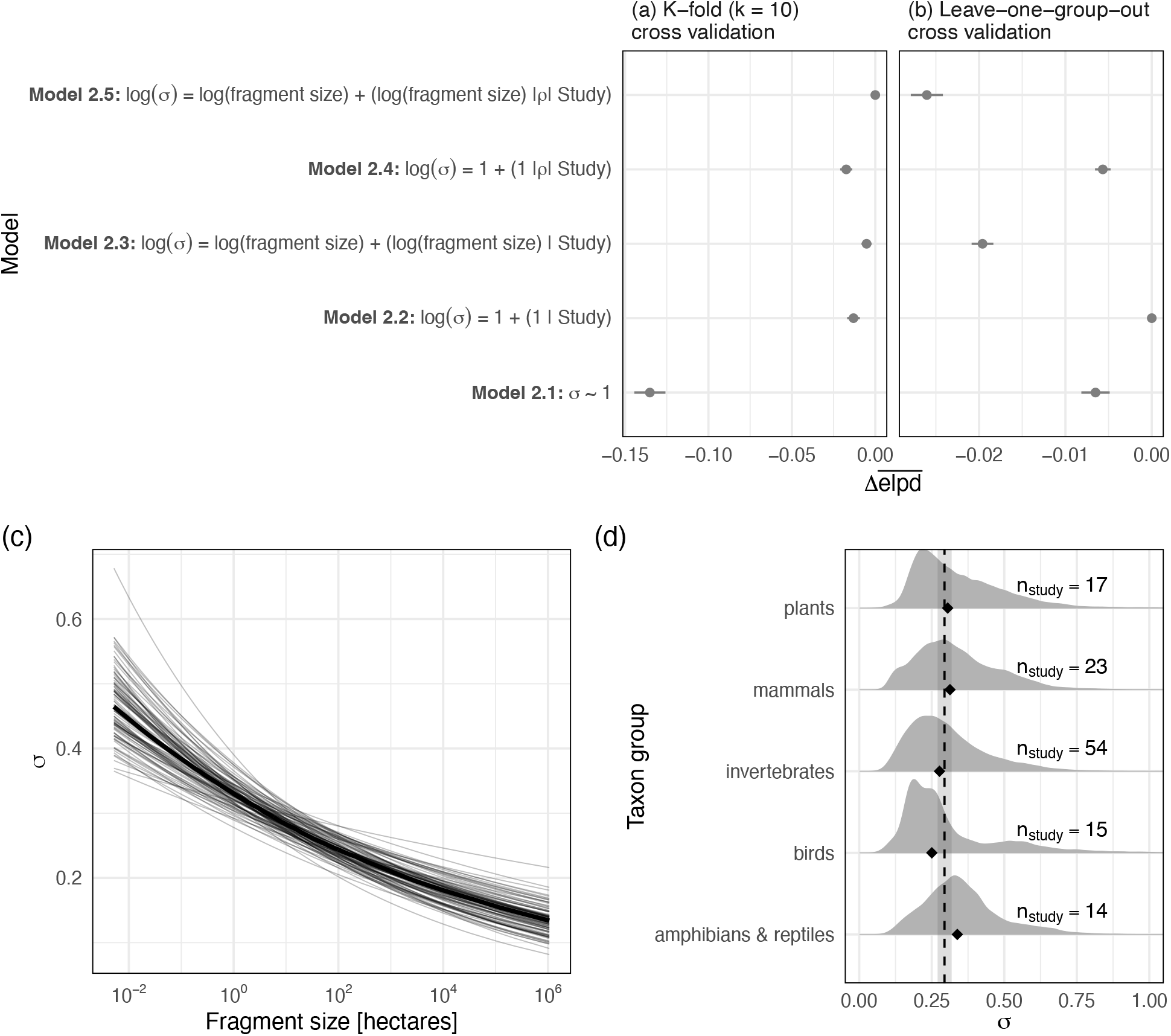
Patch-scale species richness in habitat fragments is predicted best by models with heteroscedastic residual variation. Model selection for predictions (a) within existing studies using stratified k-fold cross-validation, and (b) for predictions to new studies using leave-one-group-out cross validation; (c) fragment size-dependent residual variation, and (d) study-level residual variation grouped in taxon groups. Bold line on (c) shows median predicted average relationship between fragment size and residual variation from model 2.5, thin lines show 100 draws from the posterior distribution. Density plots (d) for taxon groups show study-level variation (1000 draws from the posterior) of residual variation from model 2.2 (i.e., 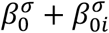 grouped into taxon groups); black triangle shows median σ for each taxon group, black dashed line and surrounding shading are the overall mean 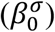 and 95% credible interval.

For predictions to new studies leave-one-group-out cross validation supported a simpler, multilevel model for residual variation (Fig. 2b), where study-level residuals varied around an overall mean (model 2.2); each study gets its own (regularized) estimate of residual variation. Study encodes unique values for many other covariates (e.g., taxon group, matrix quality, time since fragmentation), so this model can be used to further examine for systematic variation in the unexplained variation by plotting posterior samples of study-level residual variation (*σ*_*i*_) against study-level covariates (see Chase et al. 2020 for a similar approach to examining predictors of average [location] ecosystem decay effect size estimates). For example, studies of amphibians and reptiles had more unexplained variation than average, while studies of birds had less unexplained variation than average (Fig. 2d). The relatively small sample sizes (n = 14 and n = 15 studies, respectively) precludes strong inference, and I highlight this result to show: 1) explicit models of unexplained variation can help communicate where evidence for patch-scale fragmentation effects are more or less uncertain; and, (2) as an example that the more nuanced description of the data provided by the heteroscedastic model can yield new insights for researchers to build on. For example, do differences in connectivity between patches for birds compared to amphibians and reptiles lead to greater patch-scale diversity variation?

## Discussion

Quantitative evidence synthesis has become an increasingly prominent (Anderson et al. 2021) path towards generality for ecology (Gurevitch et al. 2018, Fox 2019, Spake et al. 2022). Frequently, the heterogeneity of effects are quantified across different contexts (Lau et al. 2013, Senior et al. 2016, Gurevitch et al. 2018), providing insights into how transferable effects are (Fox 2019). However, heterogeneity has typically been assumed to have constant variance, equivalent to an assumption of homoscedasticity (Williams et al. 2021, Viechtbauer & Lopez Lopez 2022, Nakagawa et al. 2025). Here, I show how location-scale models that relax the assumption of homoscedasticity can provide quantitative descriptions of where effects sizes can be transferred with more or less uncertainty, furthering the generalization goal of evidence synthesis. I also showed how cross validation can be used to advance different goals of evidence synthesis.

Relaxing the assumption of homoscedasticity in meta-analytic statistical models is relatively new (Williams et al. 2021, Rodriguez et al. 2022, Viechtbauer & Lopez-Lopez 2022, Nakagawa et al. 2025). This advent of location-scale meta-analytic models means that evidence syntheses of effects on variation (Cleasby & Nakagawa 2012) are now possible using meta-analytical models (Viechtbauer & Lopez-Lopez 2022, Nakagawa et al. 2025). For the meta-analysis case study, all heteroscedastic models produced qualitatively similar estimates of parameters shared with the homoscedastic model (Appendix S1: Fig. S3), though this won’t always be the case (Williams et al. 2021). Here I used a heteroscedastic model to model the observation that variation in native-exotic plant species richness relationships decreased with increased grain size (Peng et al. 2019), which revealed considerable uncertainty for the smallest and largest grain sizes (Fig. 1c). Heteroscedastic models also showed that the smallest extents had the most unexplained variation (Fig. 1d), suggesting that there is considerable context-dependency in plant native-exotic richness relationships at the smallest spatial scales (i.e., small grains and small extents).

Location-scale models have a longer history for statistical models fit to primary data (Lee & Nelder 1996, 2006). Heteroscedastic models for the relationship between patch-scale (standardized) species richness and fragment size quantified new patterns of variation: smaller habitat fragments exhibit greater richness variation than larger habitat fragments. Some of this relationship might be due to the typically few large fragments sampled within landscapes (i.e., less scope for residual variation among large fragments). Yet ecological variation is also possible. Small habitat fragments could experience greater variation in processes associated with ecosystem decay (e.g., demographic stochasticity, edge effects, or connectedness on a given landscape), which could promote variation in patch-scale diversity. Indeed, the most complex heteroscedastic model showed that the strongest ecosystem decay effects on average patch-scale richness accompanied the fastest decline in (residual) variation with increasing fragment size (Appendix 2: Fig. S22a, S23), suggesting that ecosystem decay processes could be amplifying patch-scale richness variation in small habitat fragments.

To date, model selection in evidence synthesis has typically used either variance explained (*R*^2^, Nakagawa et al. 2023a) or information criterion methods (Cinar et al. 2021). Here, I introduced cross validation as a flexible alternative. Cross validation can be used to evaluate a single model, or to compare multiple models (Vehtari 2025). Cross validation can also be used to assess different types of predictions, such as the within versus out-of-sample predictive performance compared in the case studies here. This adaptability means that cross validation can be tailored for the different goals an evidence synthesis might have (Gurevitch et al. 2018), and the case studies presented here showed that the ranking of models can depend on the predictive task. Within sample predictions are likely most suited to evidence syntheses where the goal is predicting the success of an intervention (Gurevitch et al. 2018), assuming that the compiled data are a representative sample of the population targeted for intervention (Boyd et al. 2023), though it is important to note that neither of the case studies had the goal of assessing intervention efficacy.

Evidence syntheses in ecology are more typically seeking broad generalizations (Gurevitch et al. 2018). For this goal, cross validation tests of out-of-sample predictions can provide direct evidence for how transferable model predictions are to different contexts (Spake et al. 2022). Here, I used leave-one-group-out cross-validation to assess out-of-sample predictions with the simplest (and most general) grouping structure typical of an evidence synthesis, i.e., predictions to a single new study. However, studies could be further grouped, e.g., by taxonomic group, (bio)geographically, or phylogenetically, and combined with a different loss function for easier interpretation (e.g., mean squared or absolute error; Yates et al. 2022) to provide a more constrained test of transferability for a single model. When comparing models, both case studies found that a model with study-level variation for the scale (residual) component was best for out-of-sample predictions, suggesting this so-called double hierarchical model (Lee & Nelder 1996, 2006) might be a good starting point for most evidence syntheses (Nakagawa et al. 2025), particularly where the goal is generalization.

Heteroscedastic models add complexity to analyses. Here, I used simulation-based calibration (Gelman et al. 2020, Talts et al. 2020, Modrák et al. 2023) to validate all of the models fit in both case studies (Appendix S1, Appendix S2). I focused on model calibration for parameter space conditioned on the observed data (Säilynoja et al. 2025). This process is particularly suited to empirical ecology as it improves confidence in model-based inferences by validating and visualizing uncertainty for known parameter values in the posterior region of the observed data (Säilynoja et al. 2025).

Location-scale statistical models and cross validation promise to strengthen quantitative evidence synthesis in ecology. Location-scale models will help communicate where effects can be transferred with more or less uncertainty, and broaden the scope of questions to include variability of effects. Cross validation can be used flexibly to meet different goals of evidence synthesis. For the case studies here, heteroscedastic models were favored for making out-of-sample predictions (i.e., to a new study), suggesting that these tools can be particularly additions to ecological evidence syntheses seeking generalities.

## Author contributions

SAB conceived and did the study.

## Acknowlegements

I thank the Biodiversity Synthesis lab for constructive discussions, and for providing much fodder for thinking about models for evidence synthesis. SAB wa here the natural logarithm of grain sizessupported by the German Centre for Integrative Biodiversity Research (iDiv) Halle-Jena-Leipzig, funded by the German Research Foundation (FZT 118), and by the European Union. Views and opinions expressed are however those of the author only, and do not necessarily reflect those of the European Union or the European Research Council. Neither the European Union nor the granting authority can be held responsible for them.

## Appendix S1

### Case-study one: Simulation-based calibration and supplemental figures

I used Simulation-Based Calibration (Talts et al. 2020, Modrák et al. 2023, Säilynoja et al. 2025) to check whether models could recover known parameter values with reasonable coverage (i.e., the probability that a constructed [e.g., credible] interval contains the true value), accuracy and precision. I focus calibration on the region of the parameter space around the empirical posterior (i.e., posterior simulation-based calibration; Säilynoja et al. 2025). Briefly, this involves: (1) fitting a model to the empirical data; (2) using the same model with priors informed by the fit to the empirical data to simulate many new (fake) datasets with the same size, shape and structure as the empirical data (i.e., the same number of observations, and the same number of groups for each level in the hierarchical structure); (3) refitting the model to each simulated data set; and, (4) calculating SBC diagnostics and plotting results of the model fits to simulated data to check for reasonable coverage (Talts et al. 2020, Modrák et al. 2023, Säilynoja et al. 2025).

To examine model calibration, I focus on three plots from the SBC diagnostics: (1) a histogram of the posterior ranks of the prior draws, which (if the algorithm and model are working correctly) should be approximately normally distributed; (2) the empirical coverage of parameters of interest (coverage is the proportion of known variable values that fall within the interval: a well calibrated model would have coverage exactly matching the interval width, e.g., 50% credible interval contains the known value 50% of the time); and, (3) a plot of estimated parameter values as a function of known (simulated) parameter values for parameters of interest, which shows how accurately and precisely the model estimates focal parameters.

#### Model 1.1

I begin case study one by reproducing the main result from Peng et al. (2019) using a multilevel linear meta-regression model for grain size that assumes the effect sizes, *z*_*ij*_, are normally distributed with known within-case variance 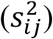 and constant between-study variance (*τ*^2^), which can be expressed as:

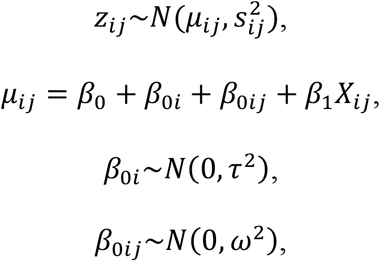

where cases (*j*) are nested within studies (*i*), and have among-case (within study) variance, *ω*^2^; study-level heterogeneity, *β*_0*i*_, has variance *τ*^2^, and varies around the overall linear relationship with intercept, *β*_0_, slope *β*_1_, and predictor *X*_*i*_ (here the natural logarithm of grain size in study *i*). The model was fit to empirical data with weakly regularizing priors:

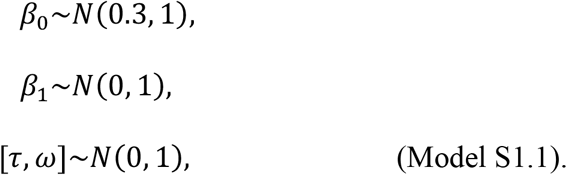

This model had good convergence (all Rhat < 1.01), and showed a reasonable fit to the empirical data (Appendix S2: Fig. S1).

**Figure S1:**
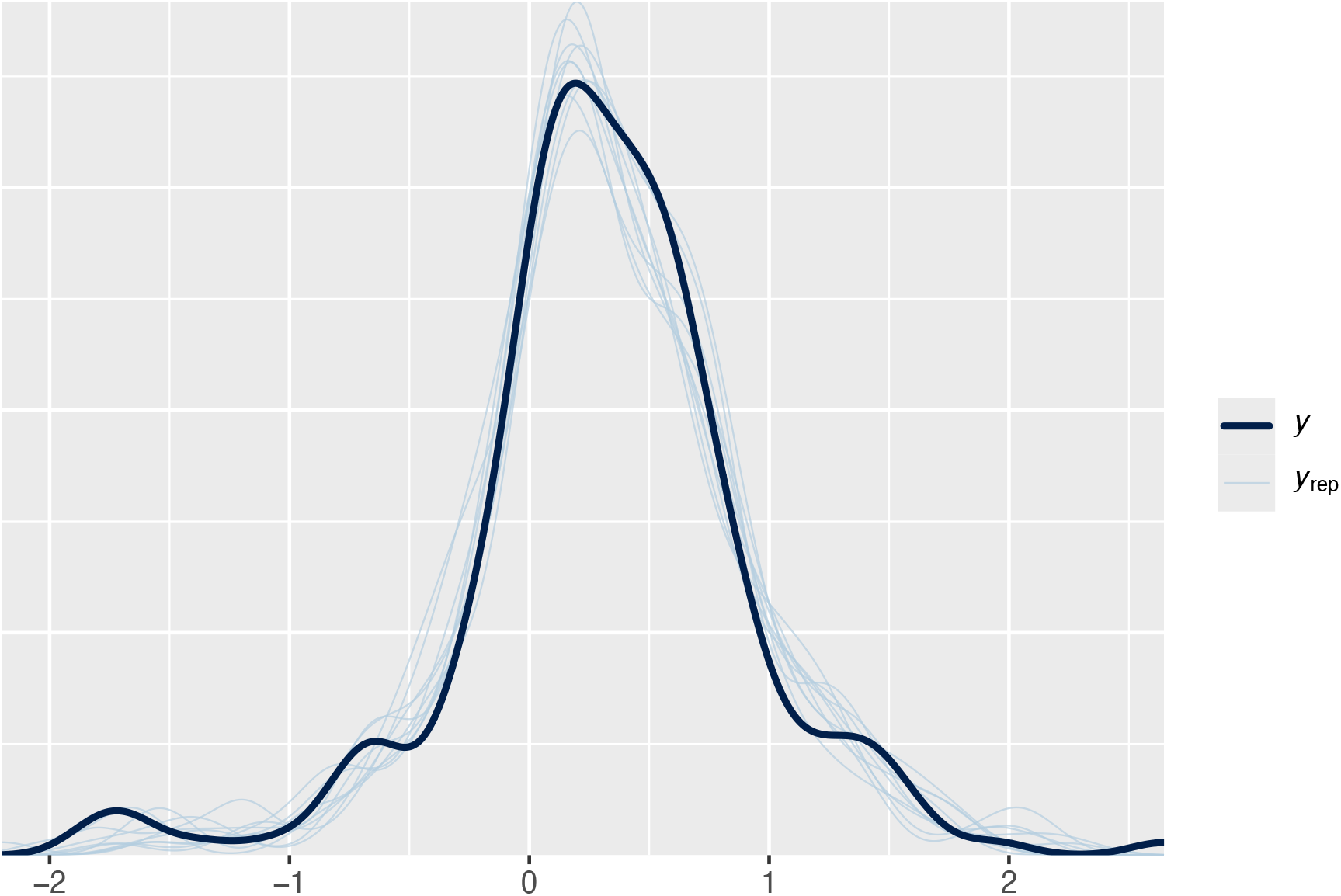
Posterior predictive check for model 1.1 fit to the empirical data (*z*-scores).

The parameter estimates from the fit of model S1.1 to empirical data (β_0_: 0.13 [95% credible interval: 0.01 – 0.24]; β_1_: 0.04 [95% credible interval: 0.02 – 0.05]; τ: 0.34 [95% credible interval: 0.25 – 0.43]; *ω*: 0.3 [95% credible interval: 0.24 – 0.35]) were used to inform the following priors:

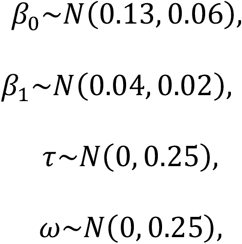

which were combined with model S1.1 to simulate (fake) data sets. To ensure as much realism as possible in the simulated data, each simulated data set retained the characteristics of the empirical data: 204 observations distributed across 101 studies (with the same balance, i.e., effect sizes per study, and the known standard errors [*s*_*ij*_] from the empirical data). I refit model S1.1 to each simulated data set, and identified models fit to fake data with poor diagnostics (e.g., divergent transitions and Rhats > 1.05), likely due to the simulated data have extreme values for the simulated data; I report the number of simulated data sets used to calculate simulation diagnostics in figure captions.

Simulation-based calibration for model 1.1 showed that the rank statistics were approximately uniformly distributed (Appendix S2: Fig. S2), and that the coverage of parameters was reasonable (Appendix S2: Fig. S3); known parameters were approximately recovered (Appendix S2: Fig. S4).

**Figure S2:**
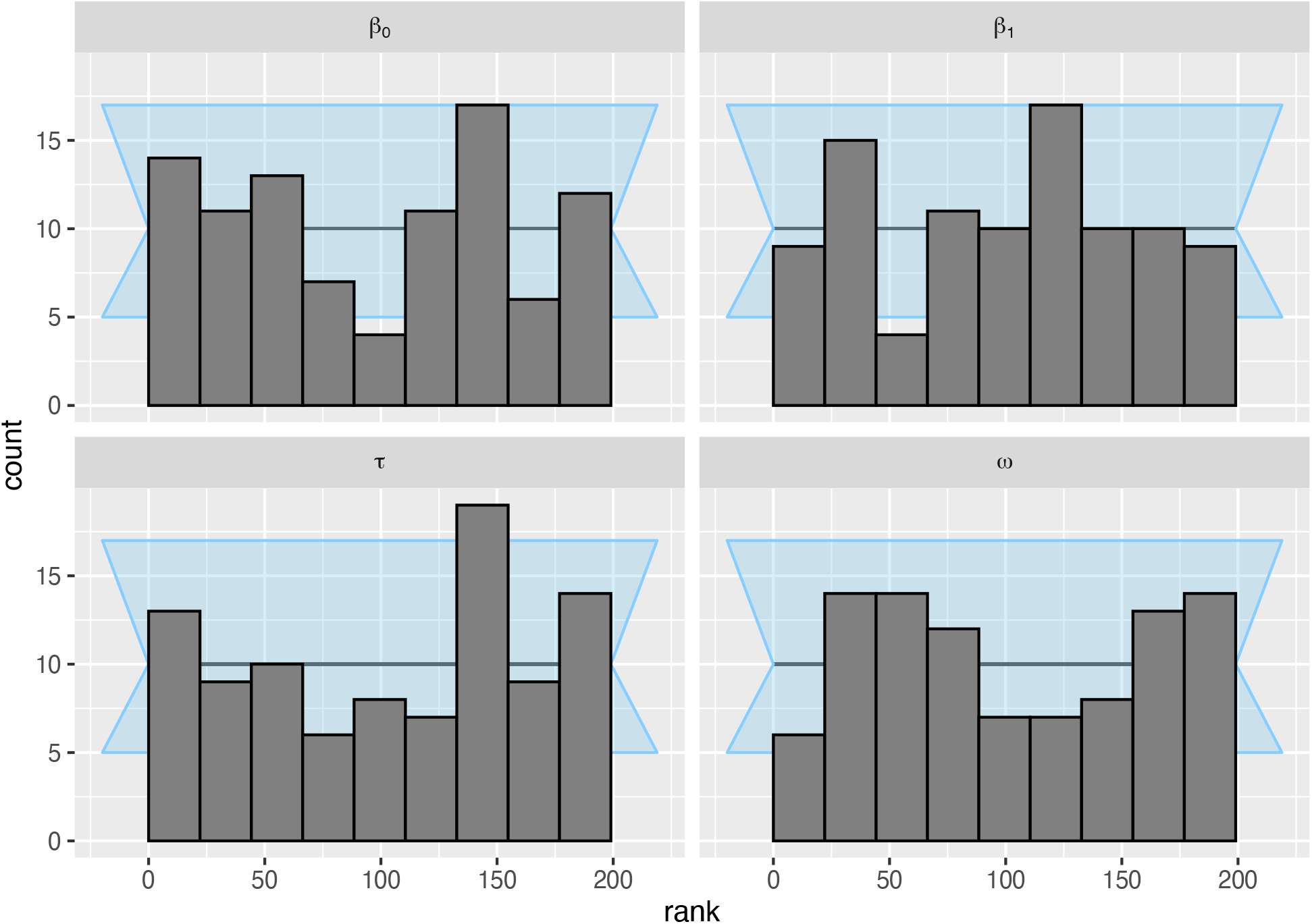
The posterior ranks of the prior draws were approximately normally distributed for the parameters of interest in model S1.1. Results are shown for n = 95 simulated data sets. Background (light blue shading) shows an approximate 95% interval for expected deviations.

**Figure S3:**
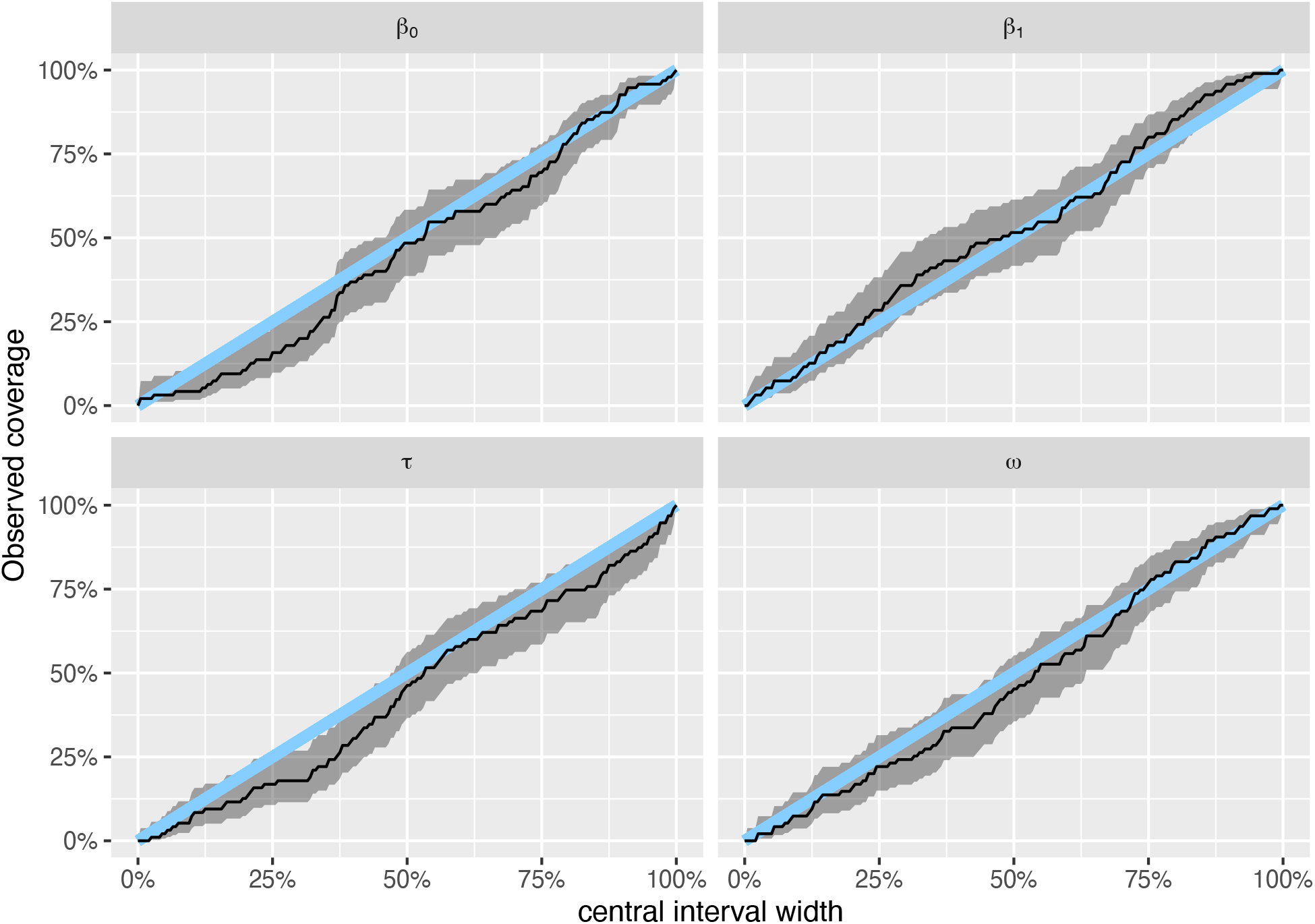
Model S1.1 had reasonable coverage for the parameters of interest. Results are shown for n = 95 simulated data sets. Blue line is 1:1 line, and shading shows 95% uncertainty interval for the coverage.

**Figure S4:**
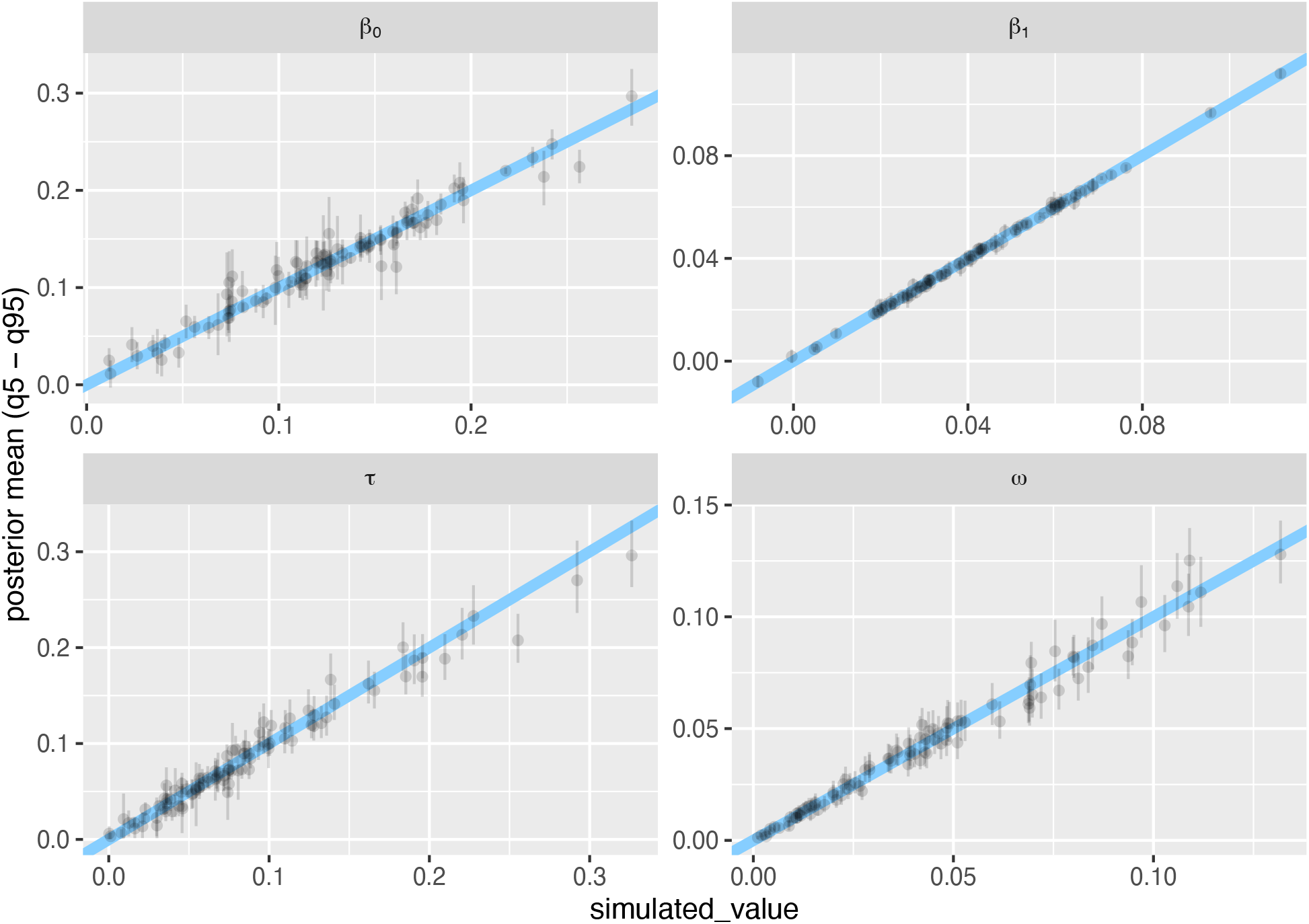
Model S1.1 was largely able to recover known parameter values. Results are shown for n = 95 simulated data sets; “simulated_value” (x-axis) is the known value of the parameter for a given simulation. Each point shows a parameter estimate, whiskers show 95% credible interval; diagonal line is the 1:1 line.

#### Model 1.2

The second model in case study one extended model S1.1 to include an additional parameter (*σ*_*ij*_) for residual variation as a function of grain size:

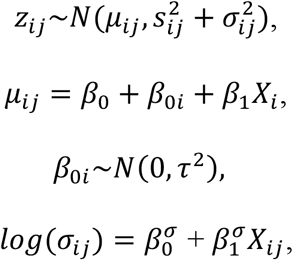

where *X*_*ij*_ is grain size on a log-scale for the *jt*h case in study *i*. I fit this model to the empirical data with weakly regularizing priors:

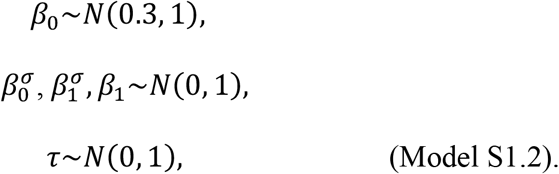

Model 1.2 had good convergence (all Rhats < 1.01), and showed a good fit to the empirical data (Appendix S2: Fig. S5).

The parameter estimates from the fit of model S1.2 to empirical data (β_0_: 0.13 [95% credible interval: 0.01 – 0.24]; β_1_: 0.04 [95% credible interval: 0.02 – 0.05]; τ: 0.33 [95% credible interval: 0.24 – 0.42]; 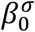: -1.12 [95% credible interval: -1.4 *–* -0.86]; 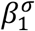: -0.03 [95% credible interval: -0.1 – 0.03]) were used to inform the following priors:

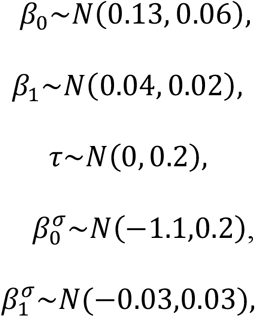

which were combined with model S1.2 to simulate (fake) data sets.

**Figure S5:**
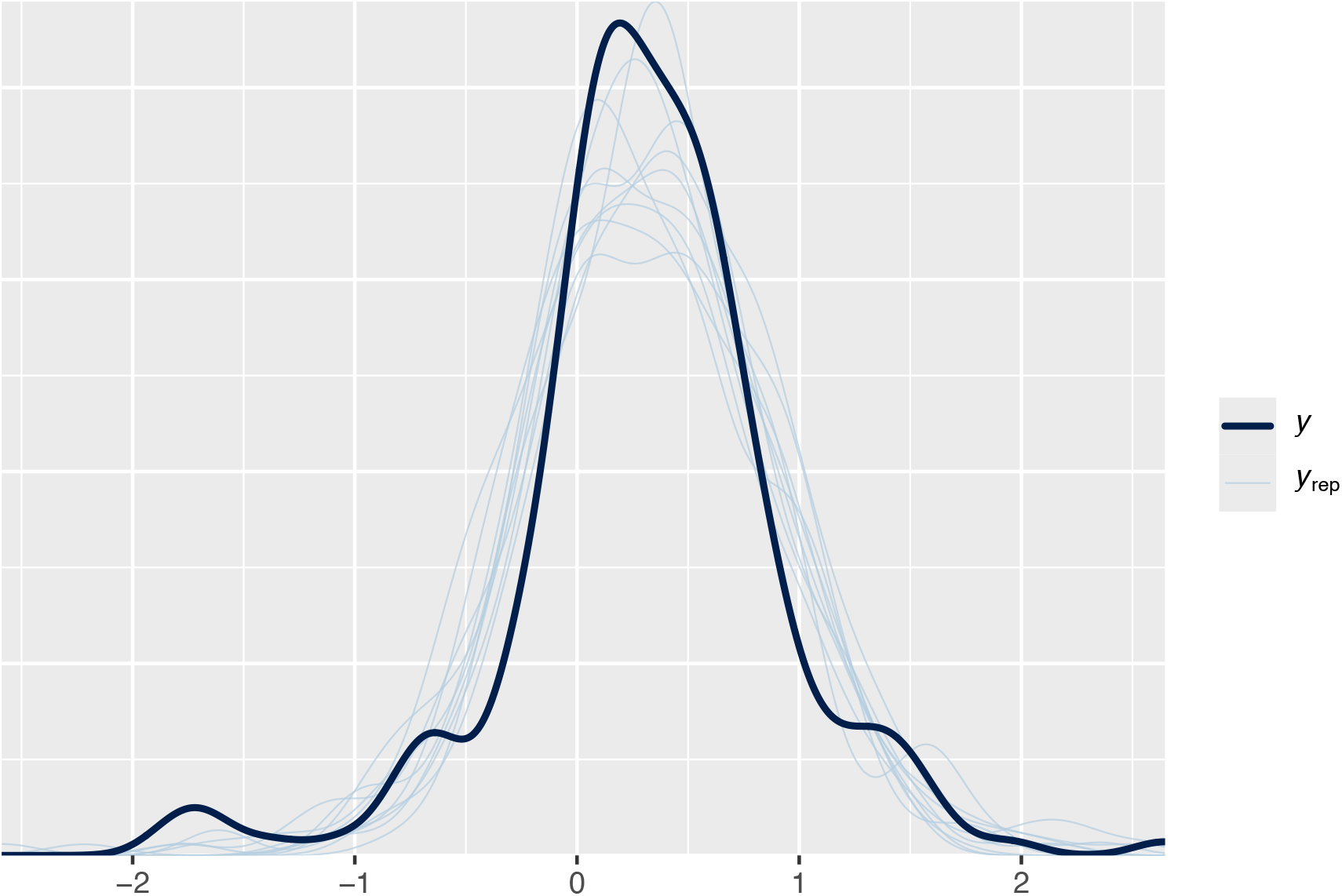
Posterior predictions from model 1.2 were largely consistent with the observed data.

Simulation-based calibration for model 1.2 showed that the rank statistics were approximately uniformly distributed (Appendix S2: Fig. S6), and that the coverage of parameters was reasonable (Appendix S2: Fig. S7); known parameters were approximately recovered, though with a fair amount of uncertainty for the intercept (β_0_), the varying intercept for studies (τ), and the slope of relationship between unexplained variation and grain size (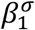 ; Appendix S2: Fig. S8).

**Figure S6:**
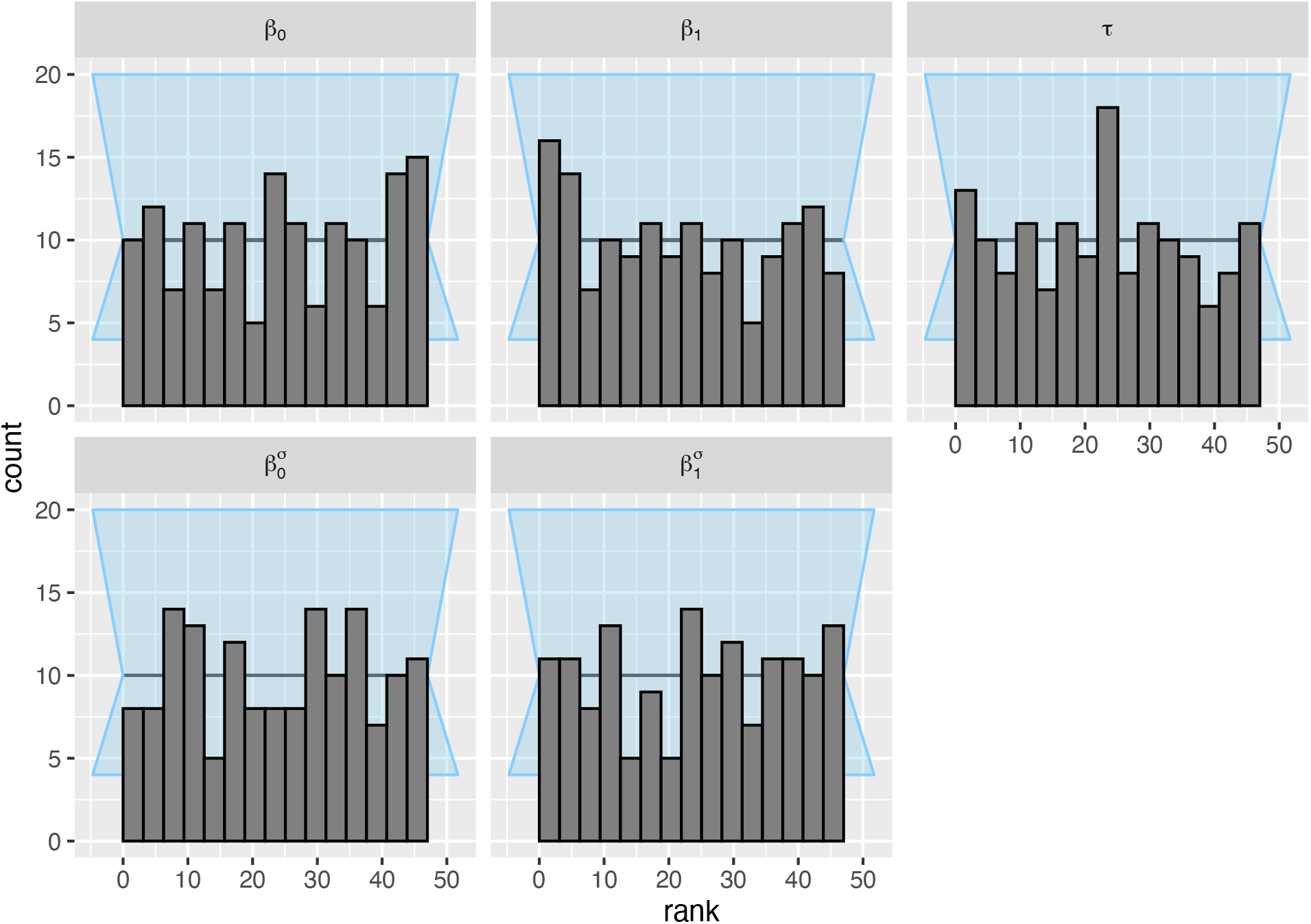
The posterior ranks of the prior draws were approximately normally distributed for the parameters of interest in model S1.2. Results are shown for n = 150 simulated data sets. Background (light blue shading) shows an approximate 95% interval for expected deviations.

**Figure S7:**
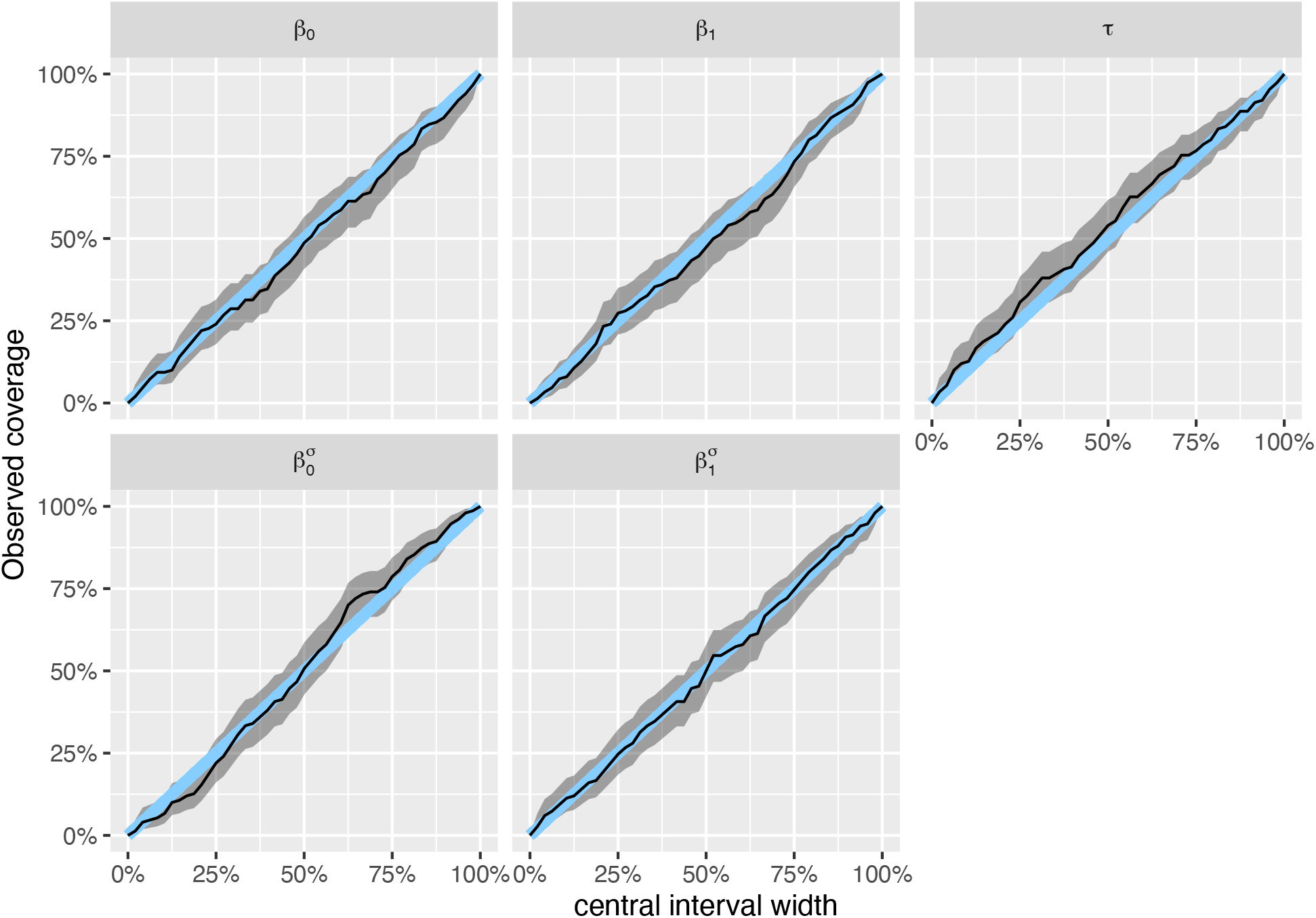
Model S1.2 had reasonable coverage for the parameters of interest. Results are shown for n = 150 simulated data sets. Blue line is 1:1 line, and shading shows 95% uncertainty interval for the coverage.

**Figure S8:**
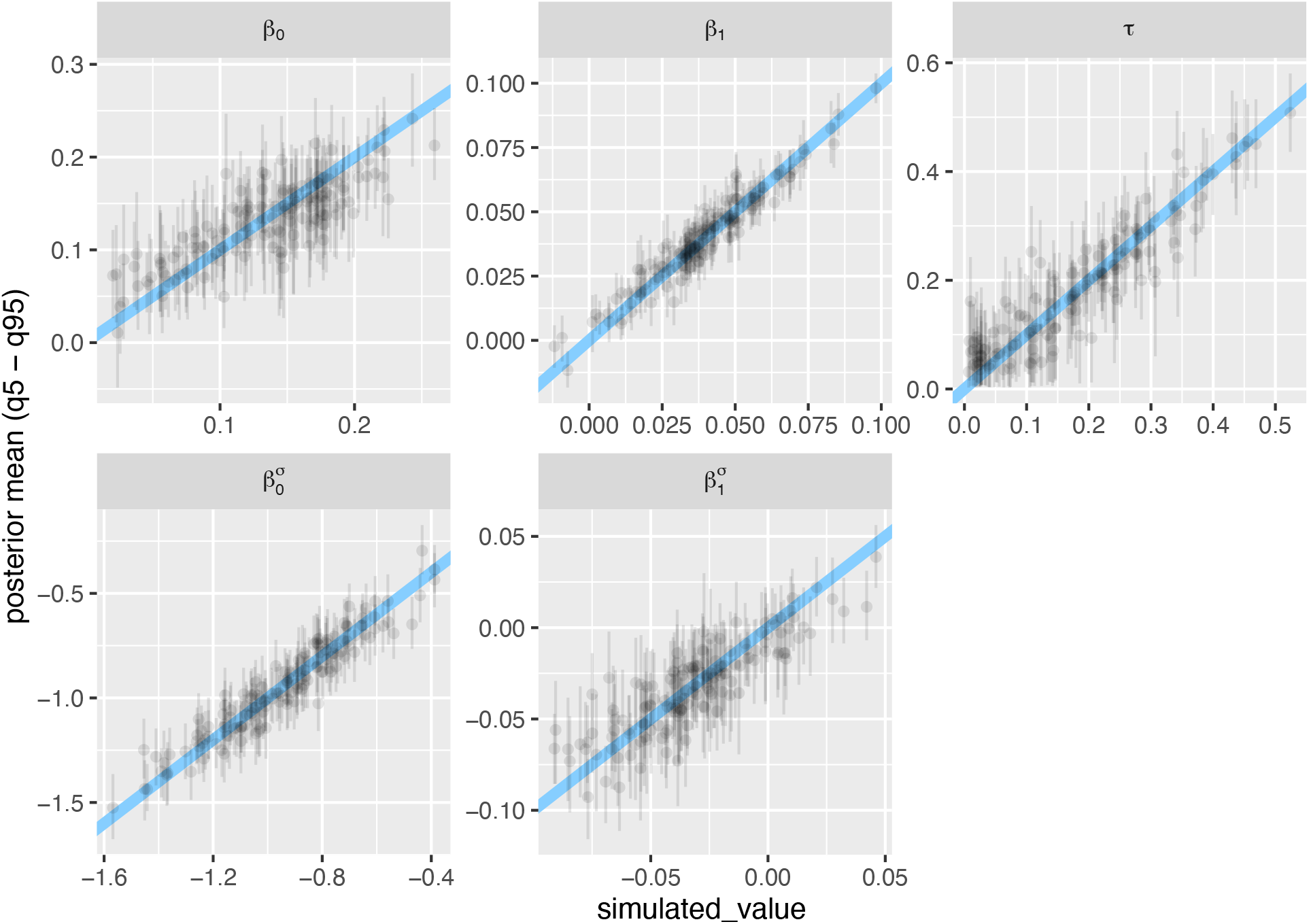
Model S1.2 was able to recover known parameter values, though with some uncertainty (especially for β_0_, τ, and 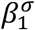). Results are shown for n = 150 simulated data sets; “simulated_value” (x-axis) is the known value of the parameter for a given simulation. Each point shows a parameter estimate, whiskers show 95% credible interval; diagonal line is the 1:1 line.

#### Model 1.3

The third model replaced grain size as a predictor of unexplained variation with spatial extent, which was estimated with indicator variables for each category of spatial extent [i.e., no intercept, see main text]). I fit this model to the empirical data with the same weakly regularizing priors as S1.2 (with the exception of 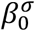, which was dropped from the model).

Model 1.3 had good convergence (all Rhats < 1.01), and showed a reasonable fit to the empirical data (Appendix S2: Fig. S9).

**Figure S9:**
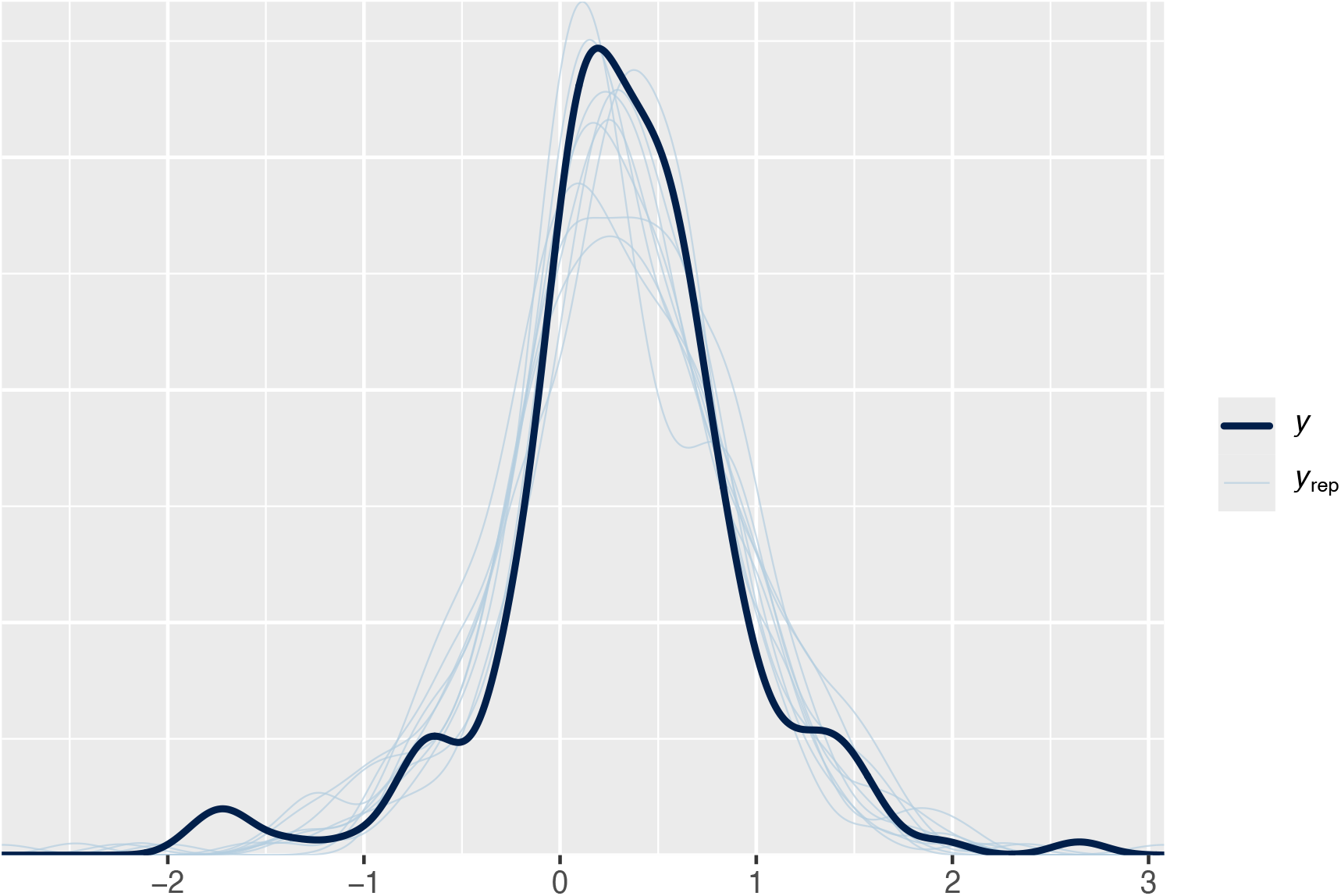
Posterior predictions for model 1.3 showed reasonable fidelity to the empirical data.

The parameter estimates from the fit of model S1.3 to empirical data (β_0_: 0.17 [95% credible interval: 0.06 – 0.28]; β_1_: 0.03 [95% credible interval: 0.02 – 0.04]; τ: 0.34 [95% credible interval: 0.25 – 0.43]; 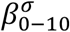: -0.79 [95% credible interval: -1.07 - -0.52]; 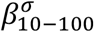: -1.58 [95% credible interval: -2.3 - -0.83]; 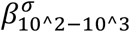: -1.03 [95% credible interval: -1.5 - -0.57]; 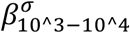: -2.09 [95% credible interval: -2.95- -1.39]; 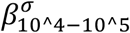: -1.66 [95% credible interval: -2.93 - -0.69]; 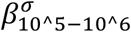: -1.53 [95% credible interval: -1.91 - -1.13]; 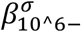: -1.51 [95% credible interval: -2.87- -0.38]) were used to inform the following priors:

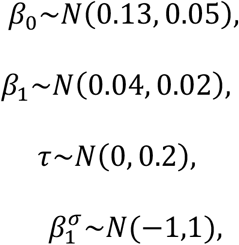

which were combined with model S1.3 to simulate (fake) data sets.

Simulation-based calibration for model 1.3 showed that the rank statistics were approximately uniformly distributed (Appendix S2: Fig. S10), and that the coverage of parameters was reasonable (Appendix S2: Fig. S11); known parameters were approximately recovered, though with a fair amount of uncertainty for the intercept (β_0_), and the varying intercept for studies (τ, Appendix S2: Fig. S12).

**Figure S10:**
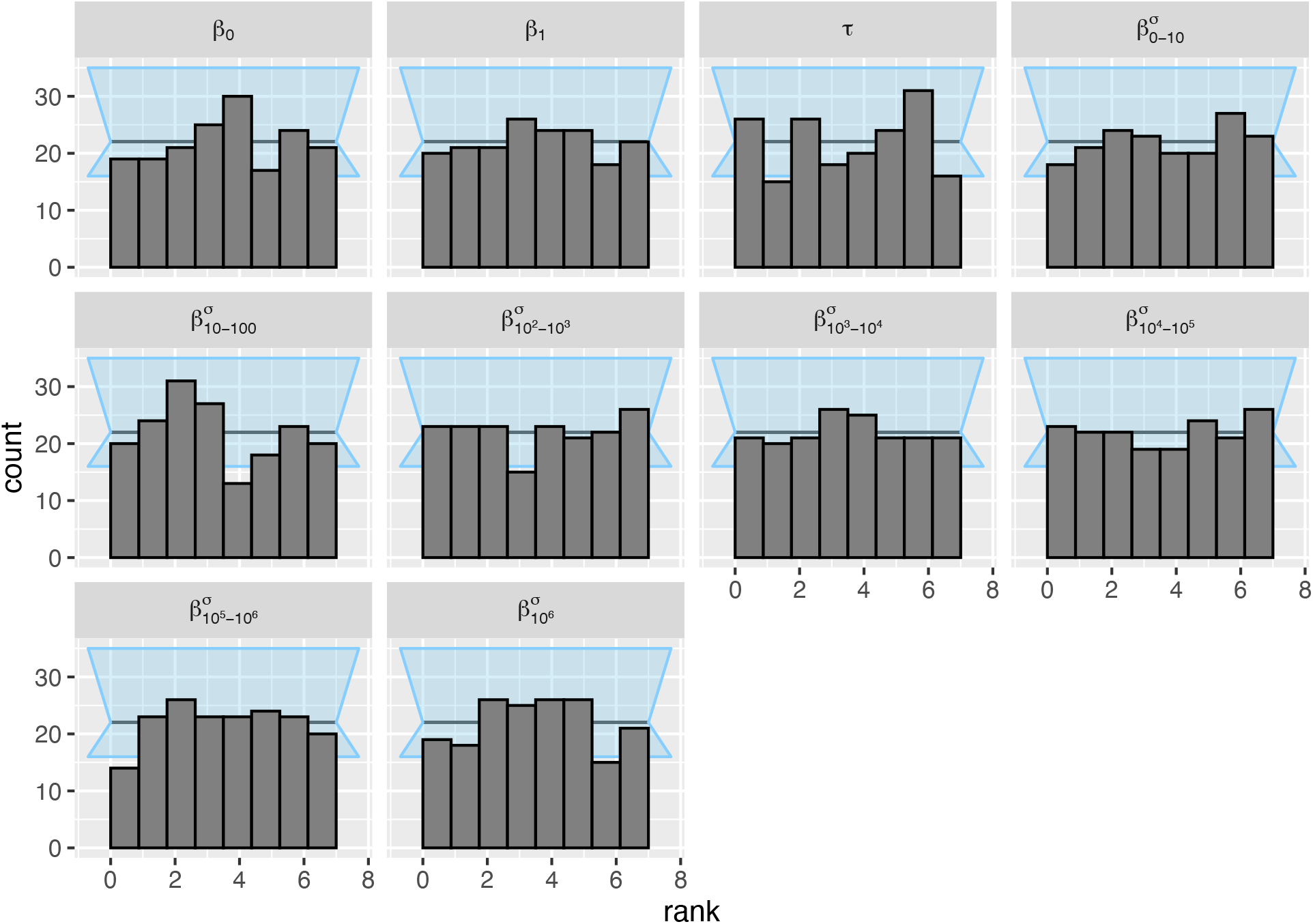
The posterior ranks of the prior draws were approximately normally distributed for the parameters of interest in model S1.3. Results are shown for n = 176 simulated data sets. Background (light blue shading) shows an approximate 95% interval for expected deviations.

**Figure S11:**
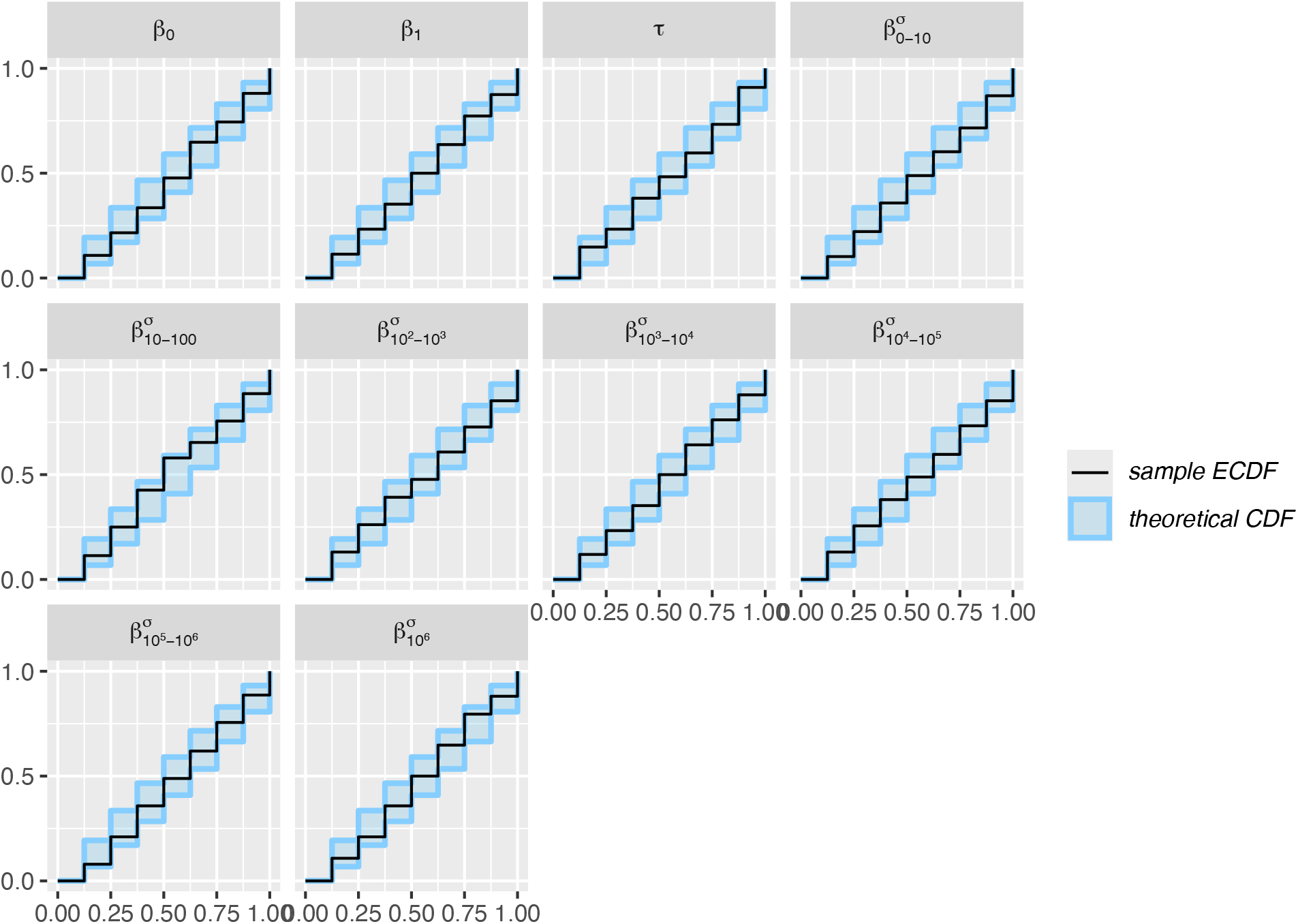
Model S1.3 had reasonable coverage for the parameters of interest. Results are shown for n = 176 simulated data sets. Blue line is 1:1 line, and shading shows 95% uncertainty interval for the coverage.

**Figure S12:**
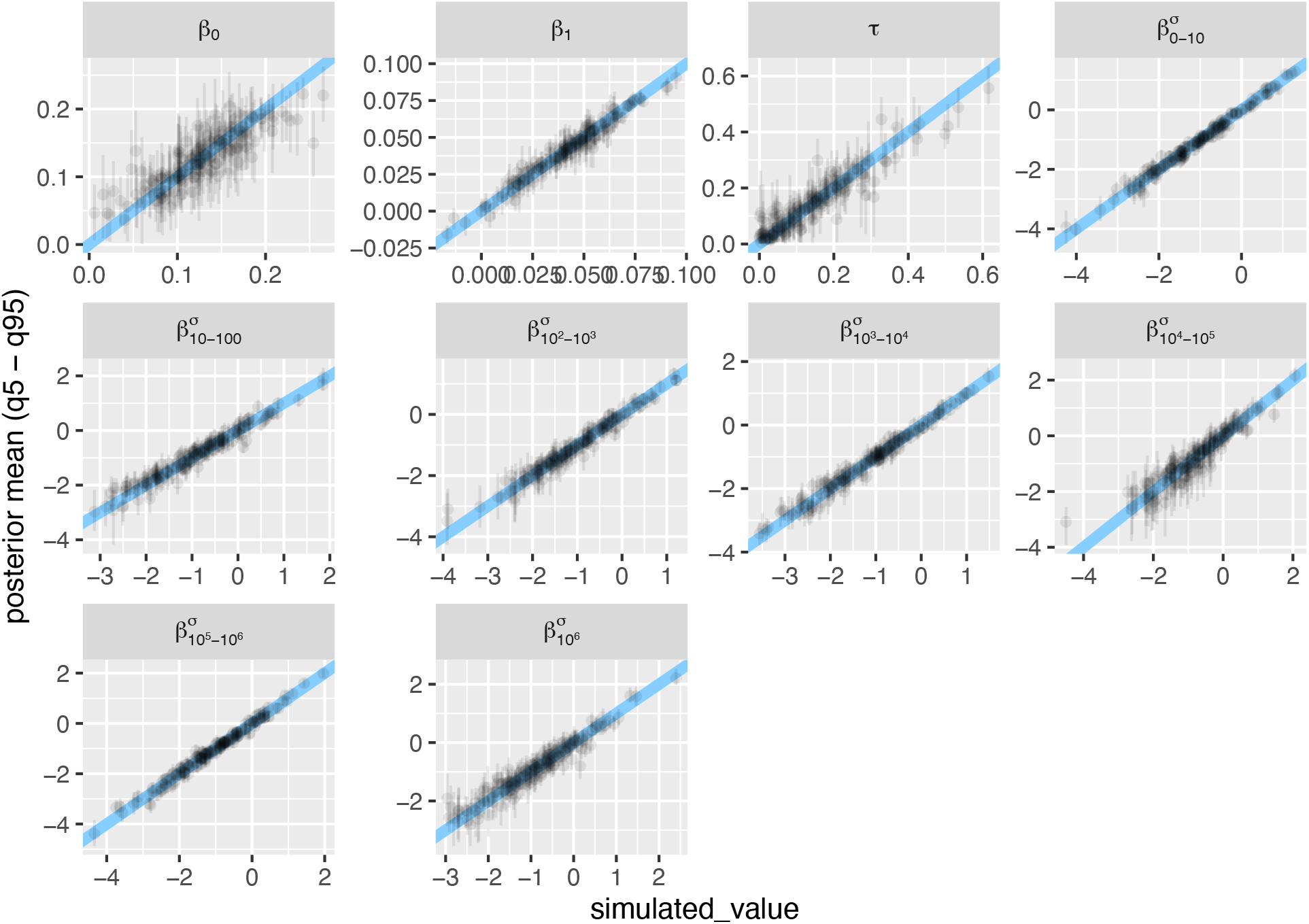
Model S1.3 was able to recover known parameter values, though with some uncertainty (most notably for β_0_, and τ). Results are shown for n = 176 simulated data sets; “simulated_value” (x-axis) is the known value of the parameter for a given simulation. Each point shows a parameter estimate, whiskers show 95% credible interval; diagonal line is the 1:1 line.

#### Model 1.4

The next model examined in the main text included varying study-level residual variation:

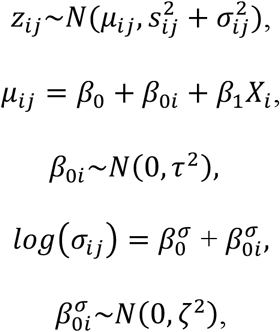

where 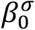 is the average residual variation (on a log-scale), and 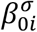 is a normally distributed study-level varying intercept with zero mean and *ζ* standard deviation. The model was fit to the empirical data with weakly regularizing parameters:

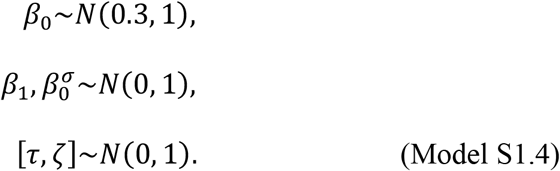

Model 1.4 had good convergence (all Rhats < 1.01), and showed a reasonable fit to the empirical data (Appendix S2: Fig. S13).

**Figure S13:**
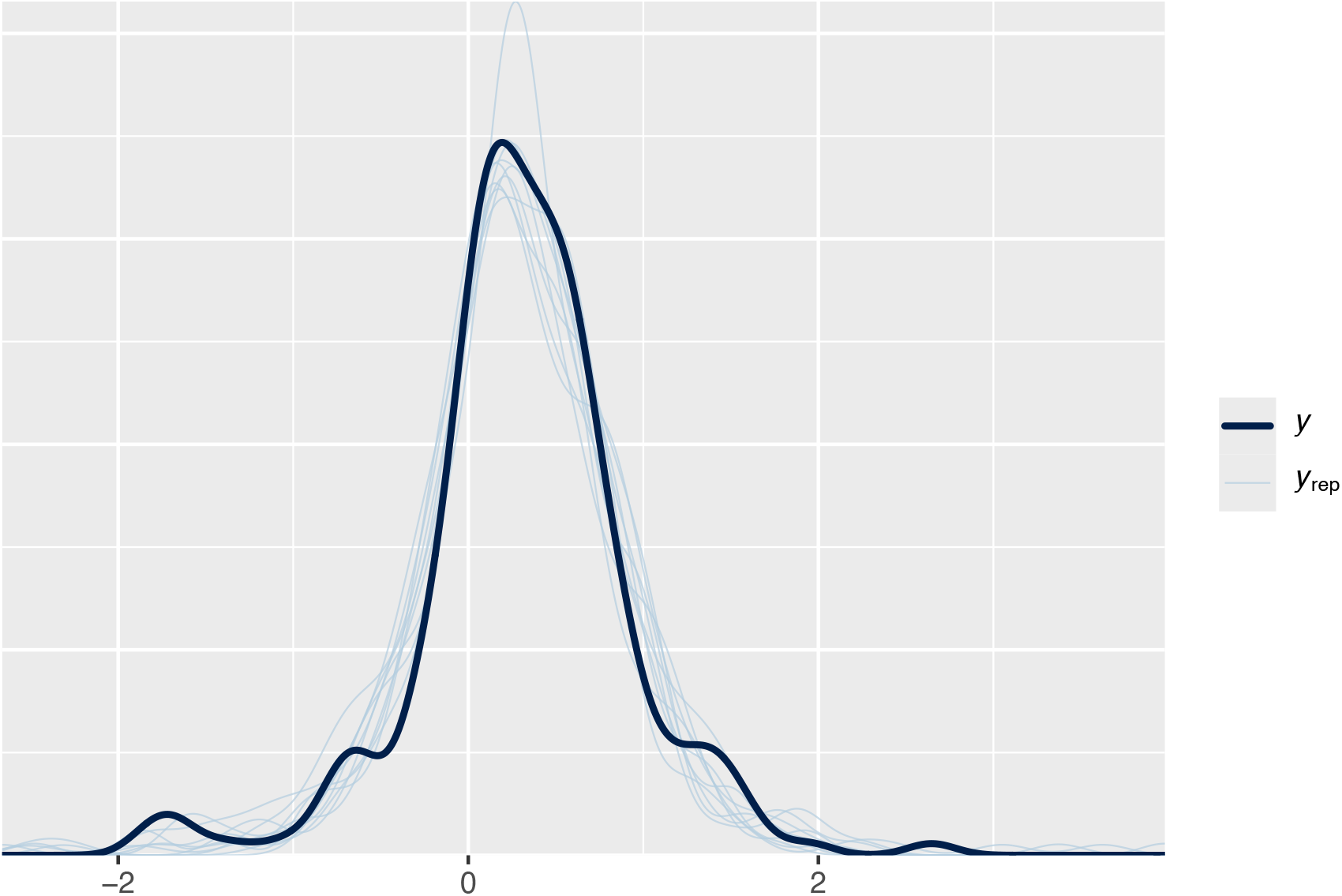
Posterior predictions from model 1.4 showed a good fit to the empirical data.

The parameter estimates from the fit of model S1.4 to empirical data (β_0_: 0.17 [95% credible interval: 0.06 – 0.27]; β_1_: 0.03 [95% credible interval: 0.02 – 0.05]; τ: 0.31 [95% credible interval: 0.22 – 0.40]; 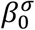 : -1.65 [95% credible interval: -2.15 *–* -1.26]; *ζ*: 0.89 [95% credible interval: 0.56 *–* 1.31]) were used to inform the following priors:

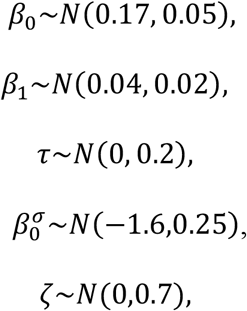

which were combined with model S1.4 to simulate (fake) data sets.

Simulation-based calibration for model 1.4 showed that the rank statistics were approximately uniformly distributed (Appendix S2: Fig. S14), and that the coverage of parameters was reasonable (Appendix S2: Fig. S15). Finally, most of the known parameters were approximately recovered, though with a fair amount of uncertainty for the intercept (β_0_), and particularly the varying intercept for study-level unexplained variation (*ζ*; Appendix S2: Fig. S16). The relatively poor (and highly uncertain) estimate of study-level unexplained variation (*ζ*) is likely due to the limited replication within studies (the median number of effect sizes per study was one; 74 of 101 studies had a single effect size).

**Figure S14:**
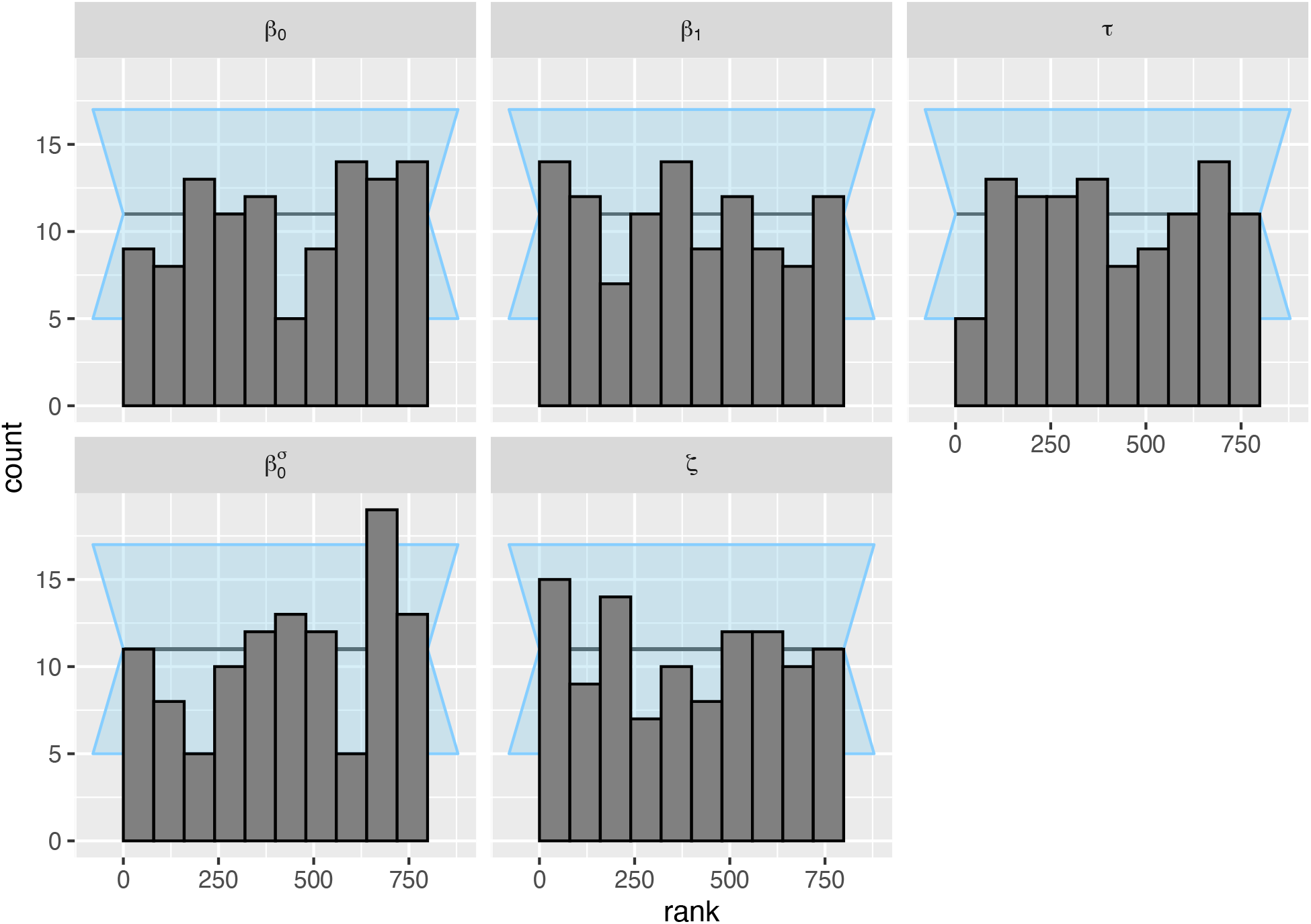
The posterior ranks of the prior draws were approximately normally distributed for the parameters of interest in model S1.4. Results are shown for n = 108 simulated data sets. Background (light blue shading) shows an approximate 95% interval for expected deviations.

**Figure S15:**
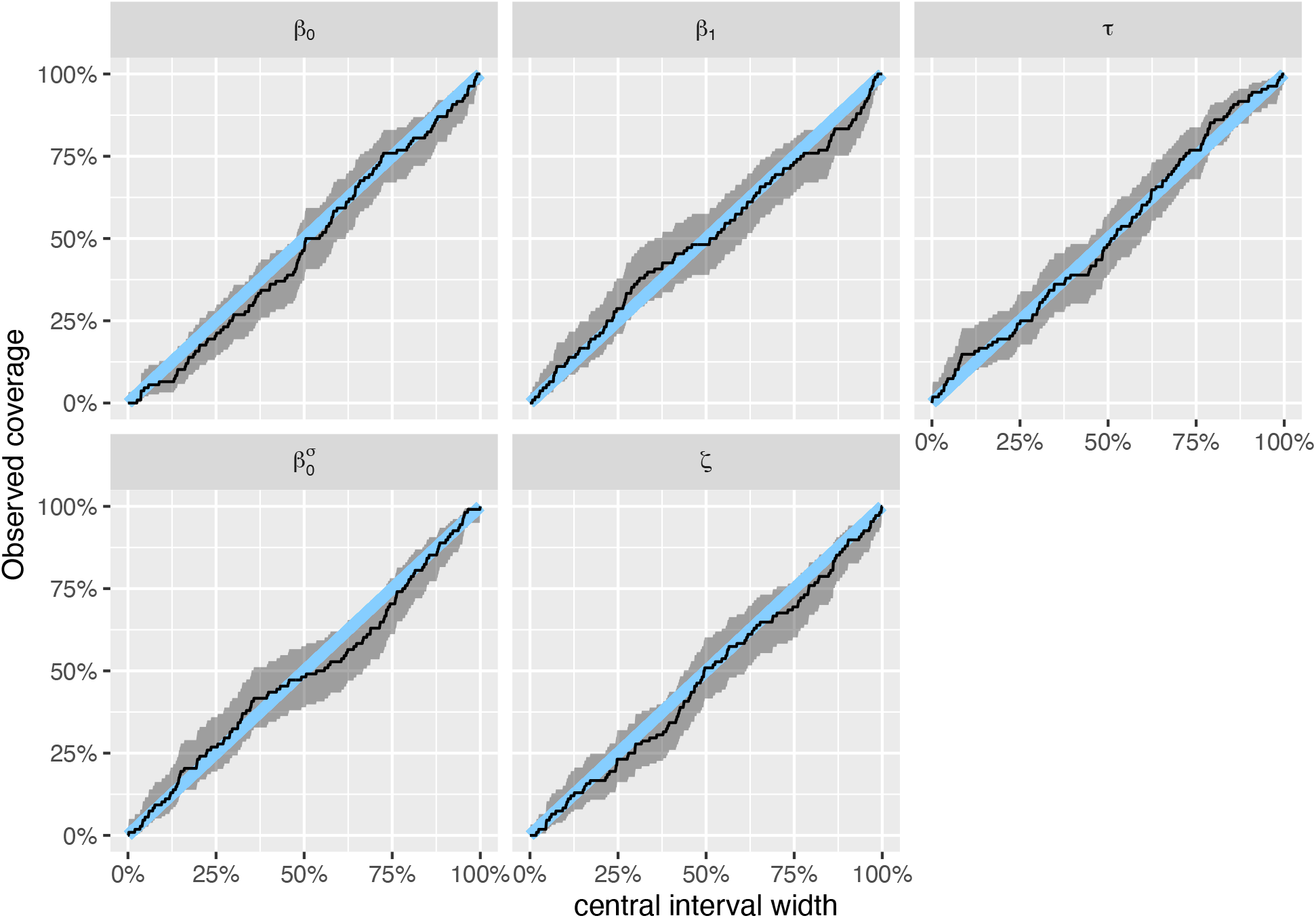
Model S1.4 had reasonable coverage for the parameters of interest, though the intercept (β_0_) showed some discrepancies. Results are shown for n = 108 simulated data sets. Blue line is 1:1 line, and shading shows 95% uncertainty interval for the coverage.

**Figure S16:**
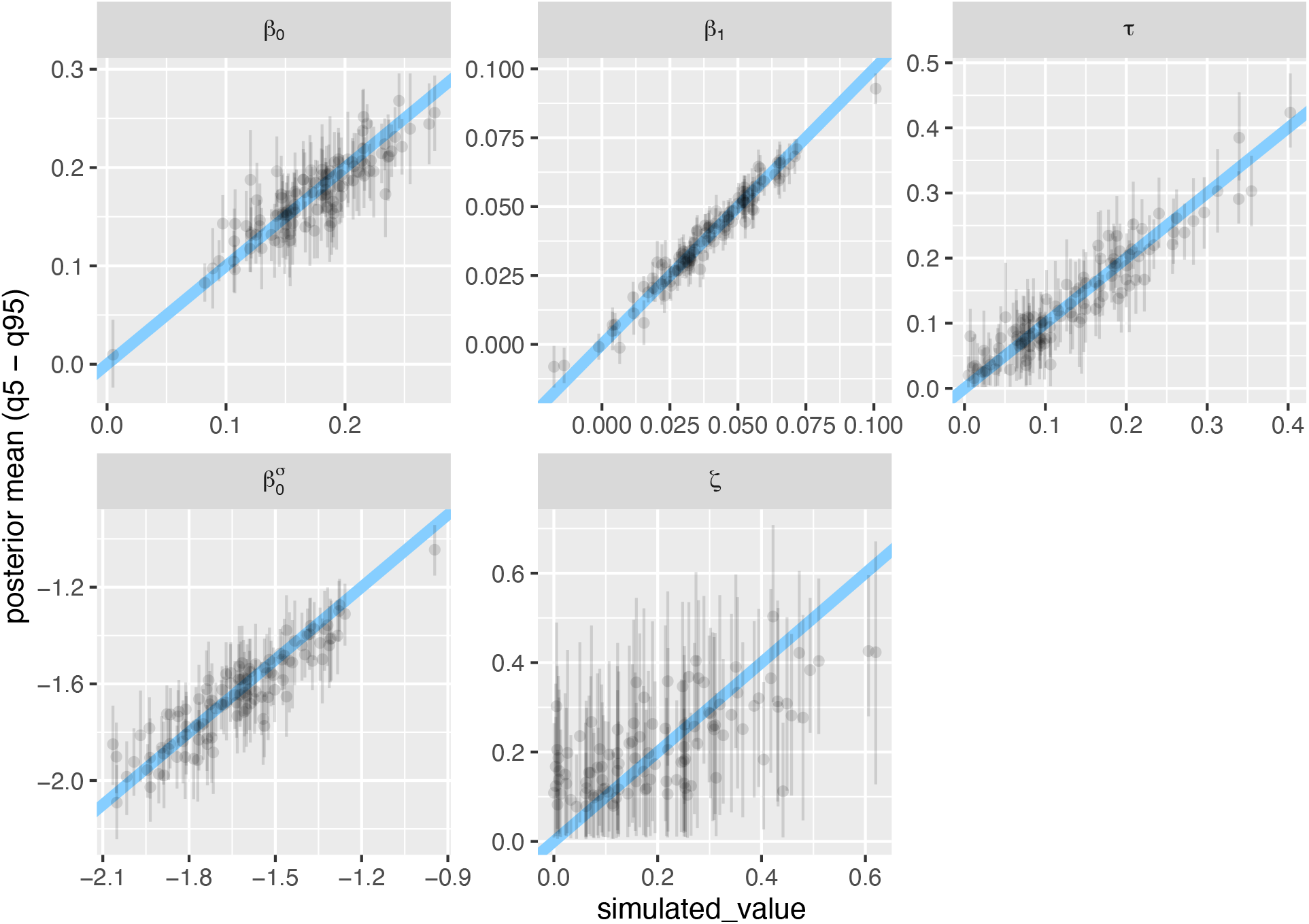
Model S1.4 was able to recover known parameter values, though with little precision for *ζ*. Results are shown for n = 108 simulated data sets; “simulated_value” (x-axis) is the known value of the parameter for a given simulation. Each point shows a parameter estimate, whiskers show 95% credible interval; diagonal line is the 1:1 line.

**Figure S17:**
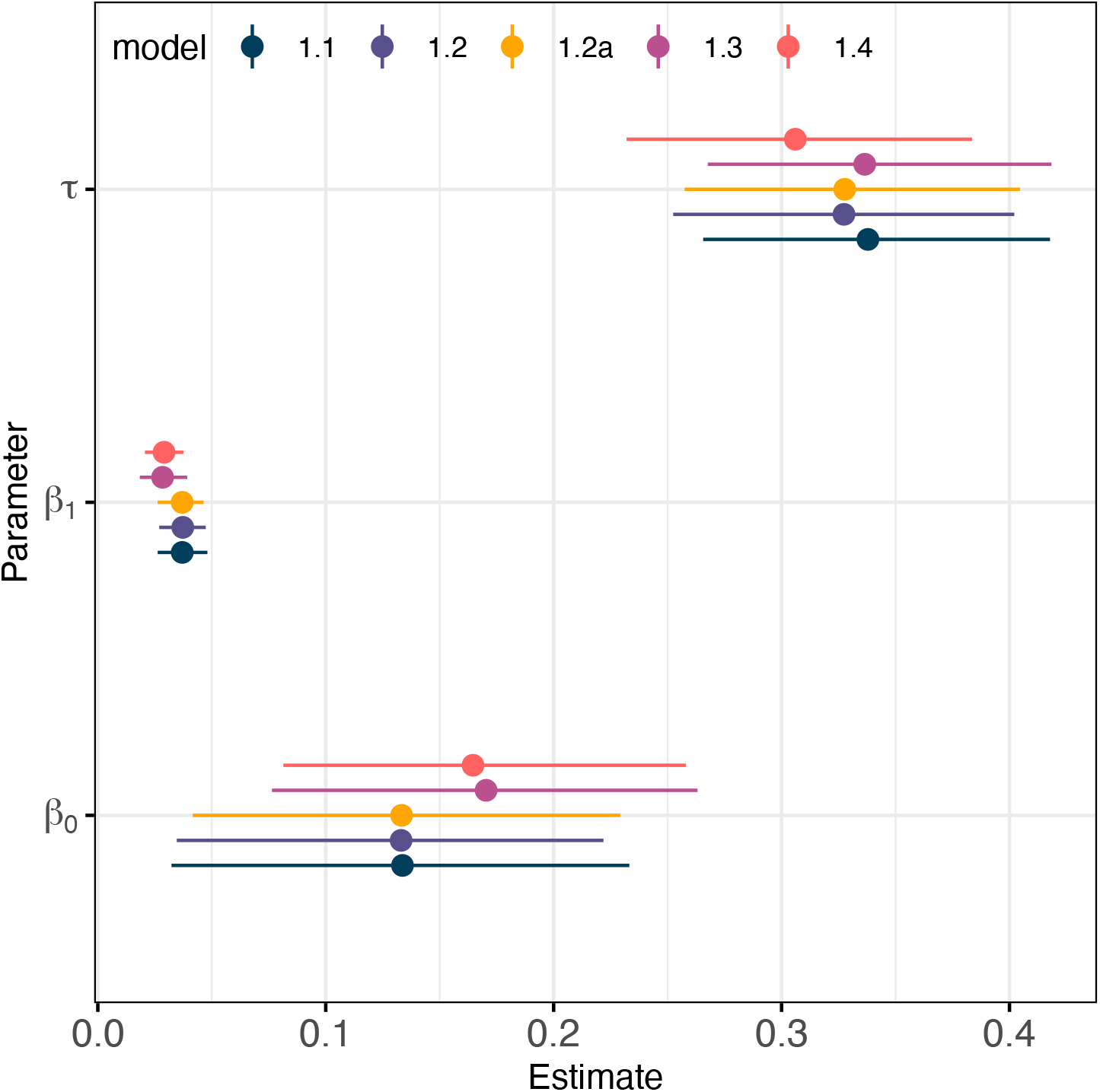
All models produced qualitatively similar estimates for shared parameters (*β*_0_, *β*_1_, *τ*).

## Appendix S2

### Case-study two: Simulation-based calibration and supplemental figures

I used Simulation-Based Calibration (Talts et al. 2020, Modrák et al. 2023) to check whether models could recover known parameter values with reasonable coverage (i.e., the probability that a constructed [e.g., credible] interval contains the true value), accuracy and precision. I focus calibration on the region of the parameter space around the empirical posterior (i.e., posterior simulation-based calibration; Säilynoja et al. 2025). Briefly, this involves: (1) fitting a model to the empirical data; (2) using the same model with priors informed by the fit to the empirical data to simulate many new (fake) datasets with the same size, shape and structure as the empirical data (i.e., the same number of observations, and the same number of groups for each level in the hierarchical structure); (3) refitting the model to each simulated data set; and, (4) calculating and plotting SBC diagnostics of model fits to simulated data to check for reasonable coverage (Talts et al. 2020, Modrák et al. 2023).

To examine model calibration, I focus on three plots from the SBC diagnostics: (1) a histogram of the posterior ranks of the prior draws, which (if the algorithm and model are working correctly) should be approximately normally distributed; (2) the empirical coverage of parameters of interest (coverage is the proportion of known variable values that fall within the interval: a well calibrated model would have coverage exactly matching the interval width, e.g., the 50% credible interval contains the known value 50% of the time); and, (3) a plot of estimated parameter values as a function of known (simulated) parameter values for parameters of interest, which shows how accurately and precisely the model estimates focal parameters.

#### Model 2.1

To start case-study two I refit the model fit by Chase et al. (2020) to effort-standardized for species richness (*S*_*ij*_) in fragment *j* from study *i*:

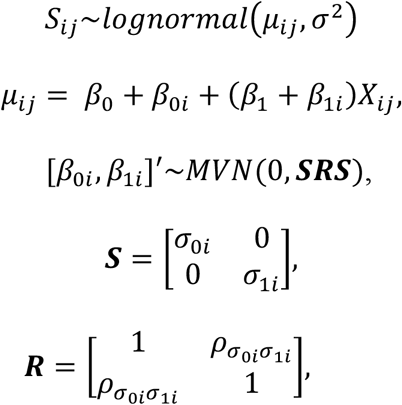

where *X*_*ij*_ is the fragment size on a (natural) log-scale, which was centered by subtracting the overall mean from each observation before modelling; *β*_0i_ and *β*_1i_ are study-level departures from the overall intercept and slope, respectively, drawn from a multivariate normal (*MVN*) distribution with standard deviations, *σ*_0*i*_ and *σ*_1*i*_, that allowed for correlations (***R*** matrix) between the varying intercepts and slopes. The model was fit to empirical data using weakly regularizing priors:

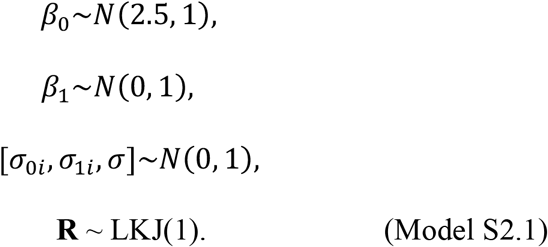

This model had good convergence (all Rhat <= 1.01), and showed a reasonable fit to the empirical data (Appendix S3: Fig. S1).

**Figure S1:**
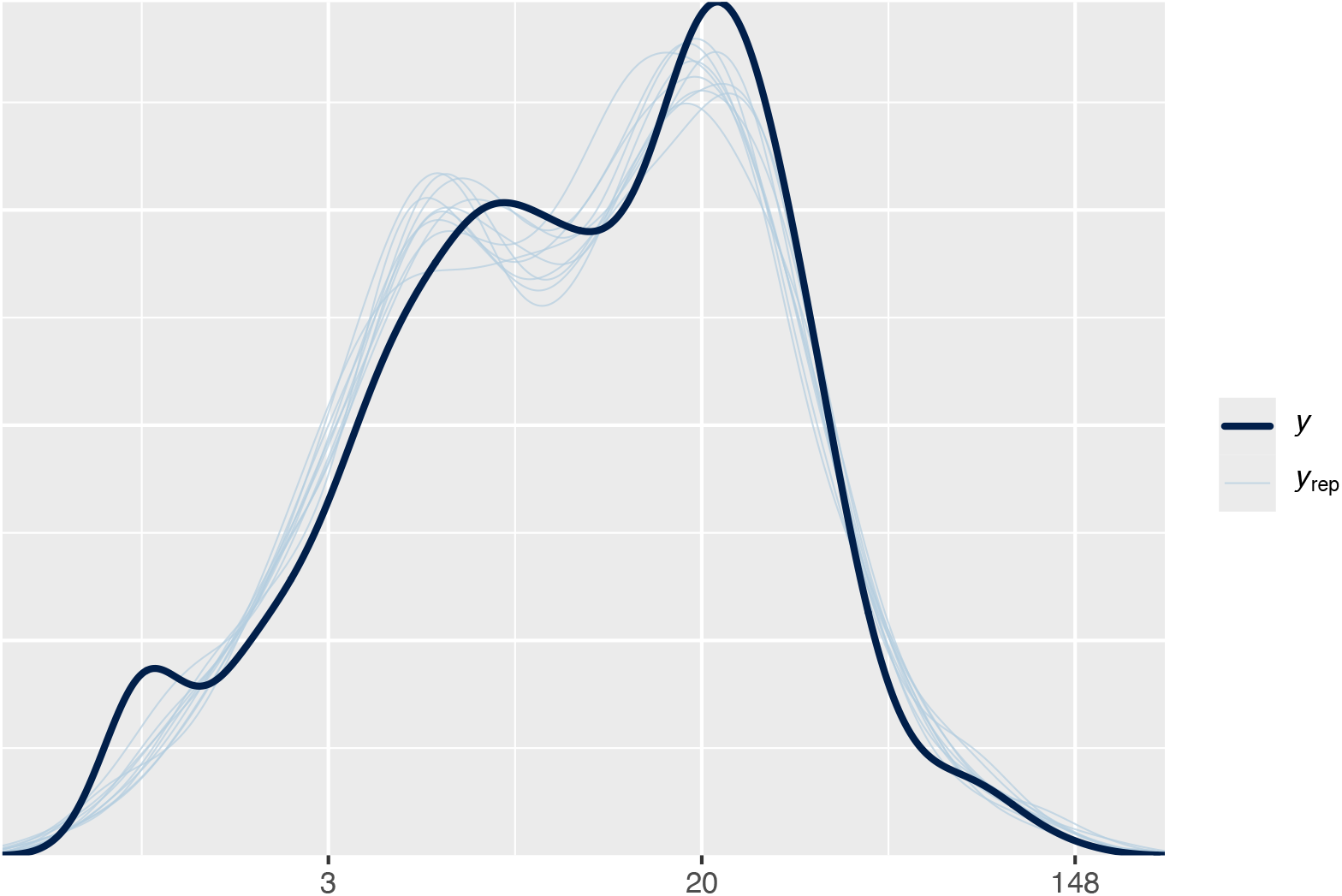
Posterior predictions from model 2.1 showed the model had a good ability to predict the observed data.

The parameter estimates from the fit of model S2.1 to empirical data (β_0_: 2.46 [95% credible interval: 2.28 – 2.65]; β_1_: 0.06 [95% credible interval: 0.04 – 0.07]; *σ*_0i_: 1 [95% credible interval: 0.88 – 1.14]; *σ*_1i_: 0.06 [95% credible interval: 0.04 – 0.08], and σ: 0.36 [95% credible interval: 0.34 – 0.37]) were used to inform the following priors:

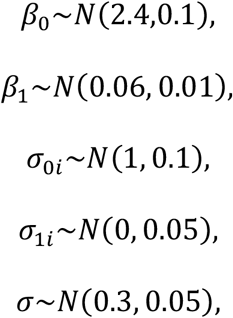

which were combined with model S2.1 to simulate (fake) data sets. To ensure as much realism as possible in the simulated data, each simulated data set retained the characteristics of the empirical data: 1509 observations distributed across 123 studies (with the same balance, i.e., fragment size distribution and fragments per study). I refit model S2.1 to each simulated data set, and identified models fit to fake data with poor diagnostics (e.g., divergent transitions and Rhats > 1.05), likely due to the simulated data have extreme values for the simulated data; I report the number of simulated data sets used to calculate simulation diagnostics in figure captions.

Simulation-based calibration for model 2.1 showed that the rank statistics were approximately uniformly distributed (Appendix S3: Fig. S2), and that the coverage of parameters was reasonable (Appendix S3: Fig. S3); known parameters were recovered with reasonable accuracy (Appendix S3: Fig. S4). Note, (here and in subsequent calibrations for model extensions) I do not show parameters associated with the intercept (i.e., β_0_, *σ*_0i_) for the location component of the models, as these parameters are not readily interpretable. They correspond to the average richness (across all studies), and study-level variation around the overall average. Richness (and other diversity measures) are not readily comparable across this heterogeneous data compilation: richness in any given data set will depend on the sample grain and size of the species pool (among other things).

**Figure S2:**
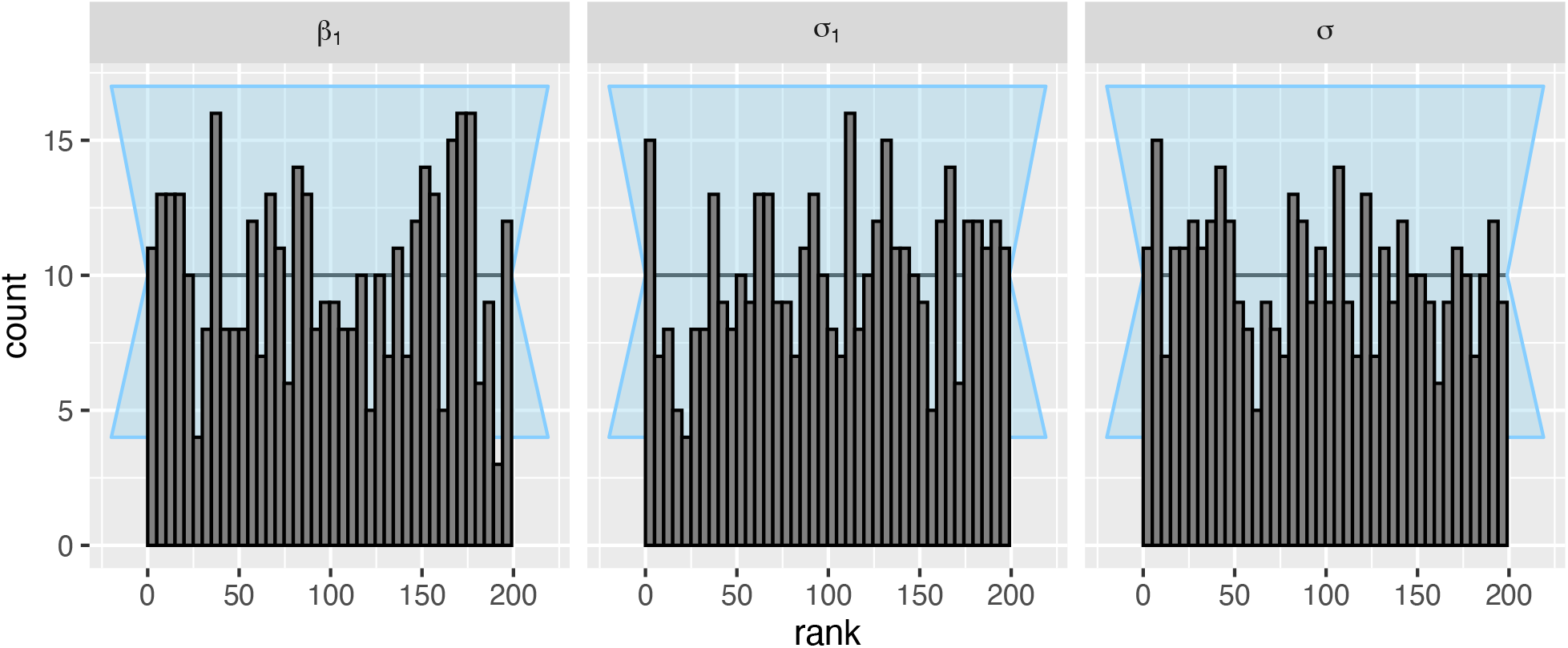
The posterior ranks of the prior draws were approximately normally distributed for the parameters of interest in model S2.1. Results are shown for n = 401 simulated data sets. Background (light blue shading) shows an approximate 95% interval for expected deviations.

**Figure S3:**
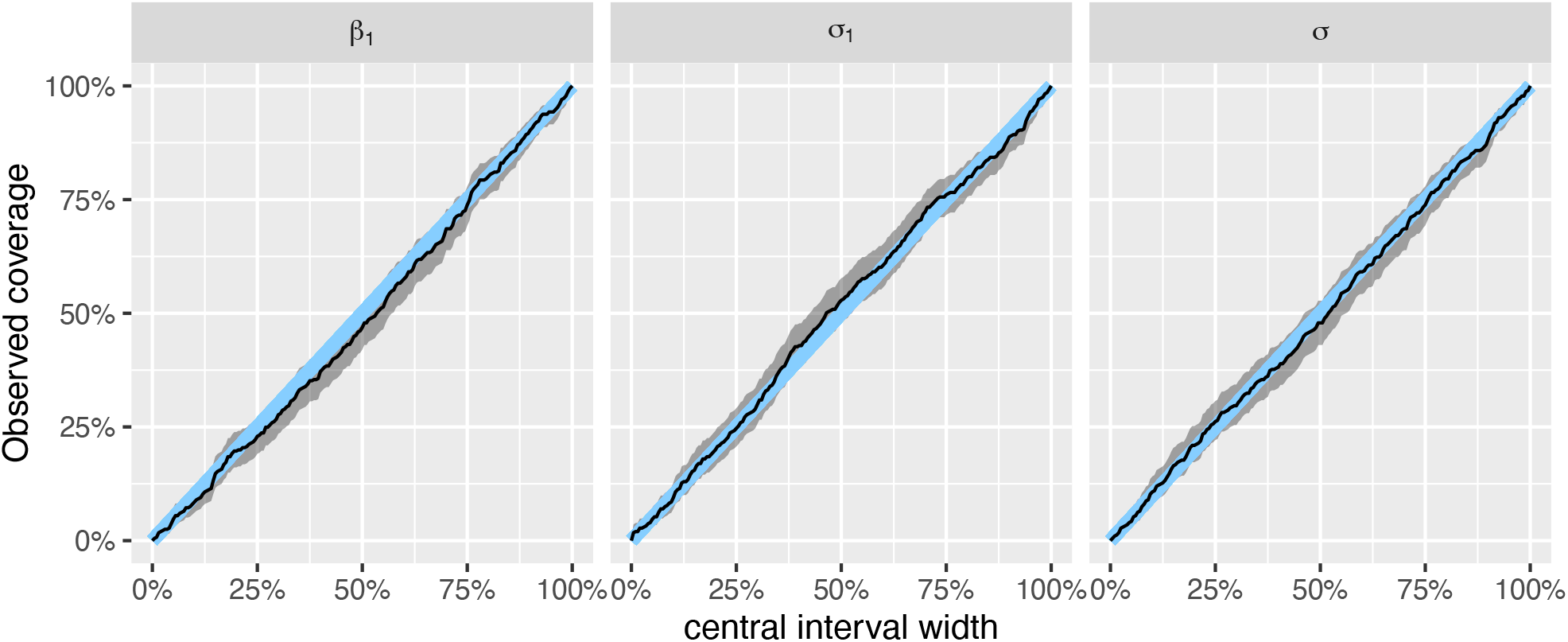
Model S2.1 had good coverage for the parameters of interest. Results are shown for n = 401 simulated data sets. Blue line is 1:1 line, and shading shows 95% uncertainty interval for the coverage.

**Figure S4:**
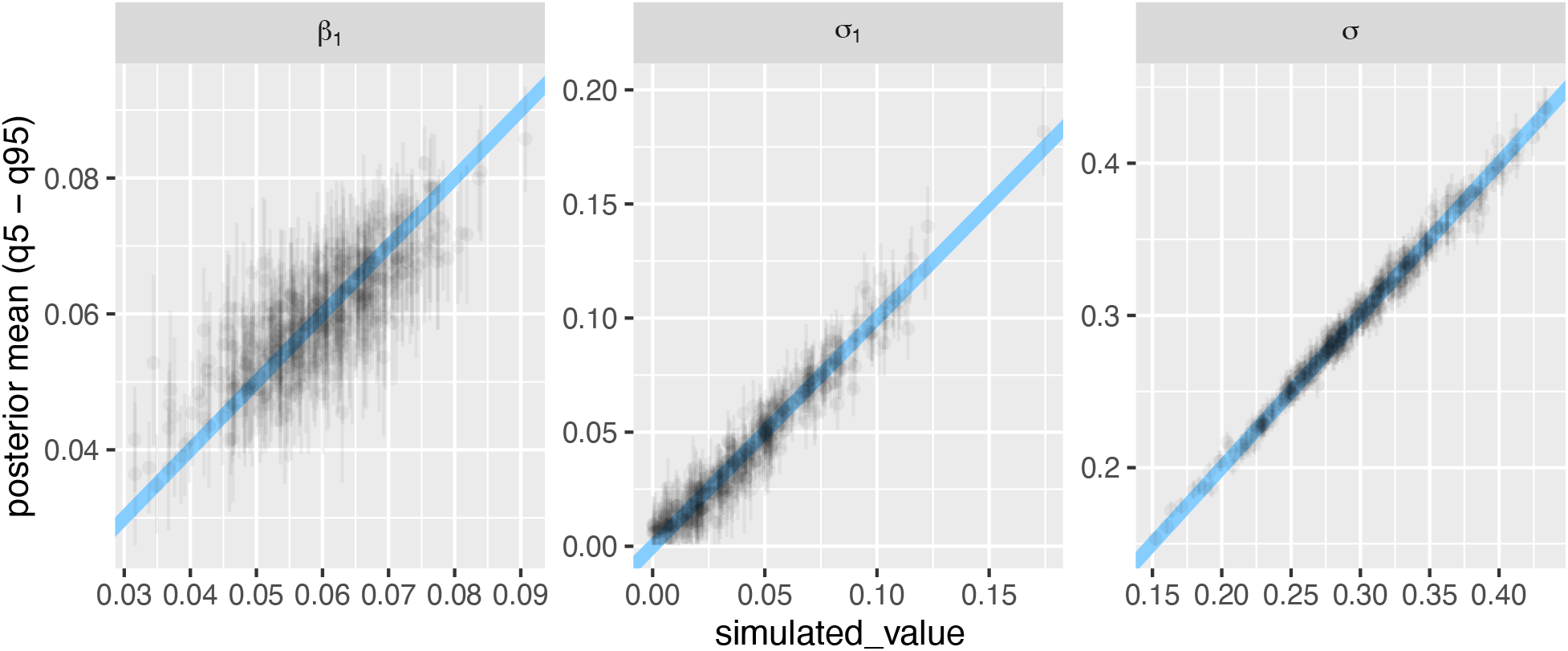
Model S2.1 was able to recover known parameter values, though with considerable uncertainty (with the exception of *σ*_1_ and σ). Results are shown for n = 401 simulated data sets; “simulated_value” (x-axis) is the known value of the parameter for a given simulation. Each point shows a parameter estimate, whiskers show 95% credible interval; diagonal line is the 1:1 line.

#### Model 2.2

The first heteroscedastic model is motivated similarly to the varying intercepts and slopes per study, and allowed varying study-level residuals around an overall average residual standard deviation. I estimated study-level residual variation i parameters for the mean:

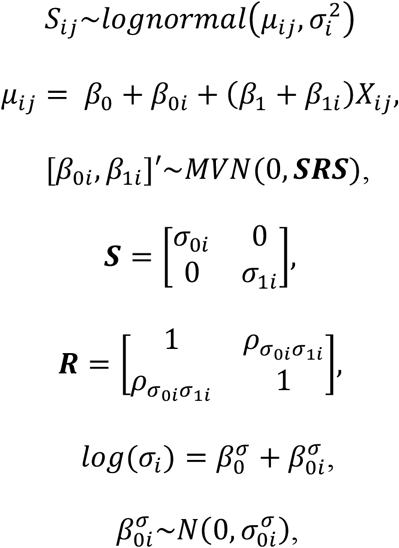

where 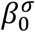 is the overall average of residual standard variation (on a log-scale), and 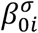 are the varying study-level departures (for the residual variation) drawn from a normal distribution with zero mean and standard deviation 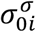 I fit the model with weakly regularizing priors:

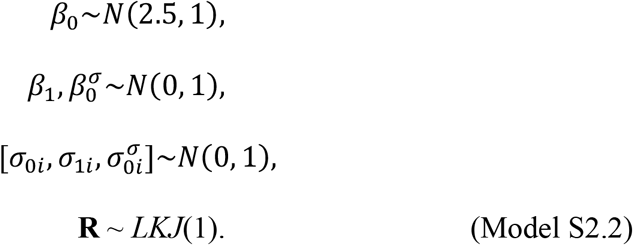

This model had good convergence (all Rhat <= 1.01), and showed a reasonable fit to the empirical data (Appendix S3: Fig. S5).

**Figure S5:**
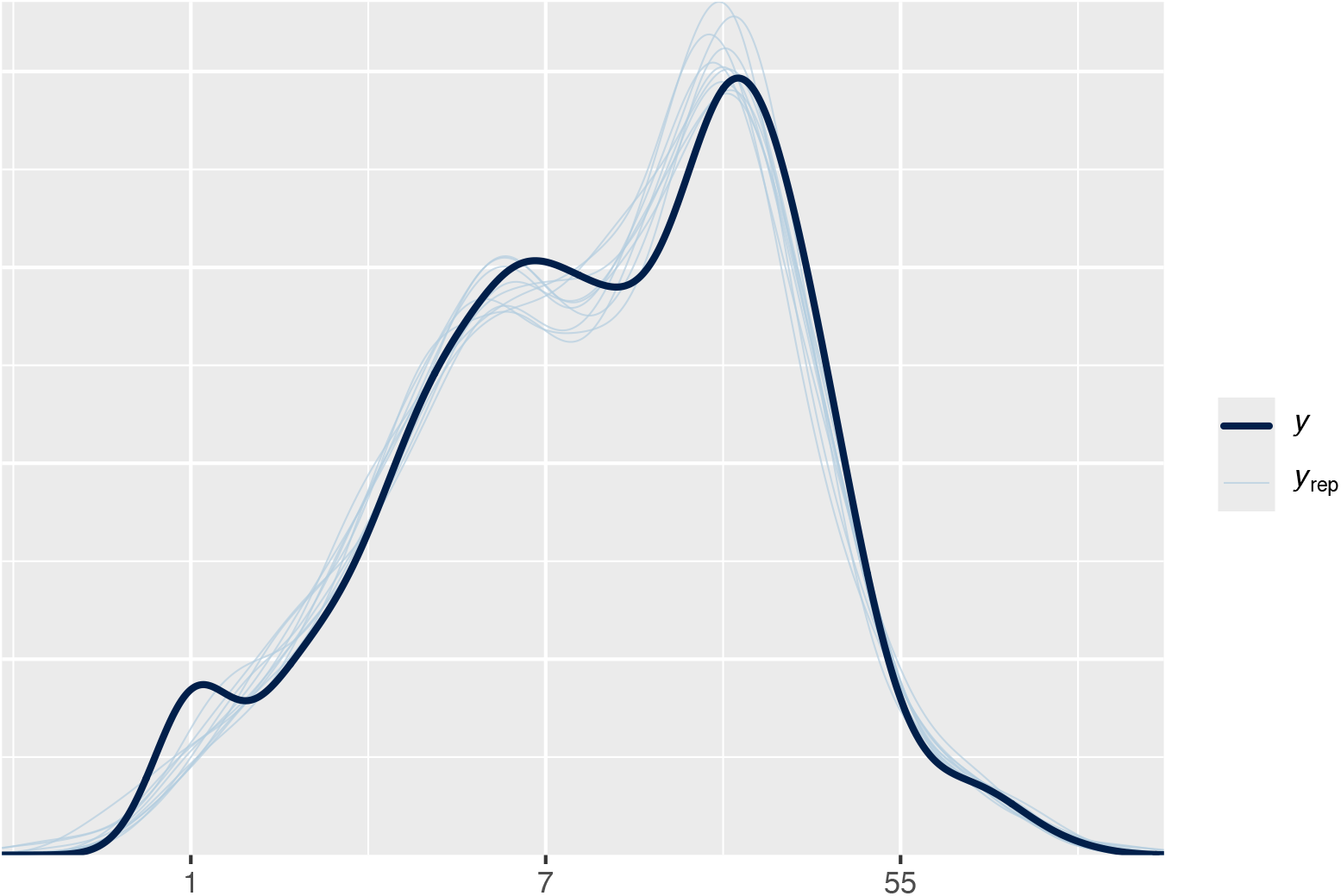
Posterior predictions from model 2.2 showed the model had a good ability to predict the observed data.

The parameter estimates from the fit of model S2.2 to empirical data (β_0_: 2.48 [95% credible interval: 2.31 – 2.66]; β_1_: 0.05 [95% credible interval: 0.04 – 0.07]; *σ*_0i_: 1 [95% credible interval: 0.88 – 1.15]; *σ*_1i_: 0.06 [95% credible interval: 0.04 – 0.07] 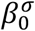 : -1.23 [95% credible interval: -1.32 – -1.13], and 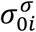 : 0.44 [95% credible interval: 0.36 *–* 0.53]) were used to inform the following priors:

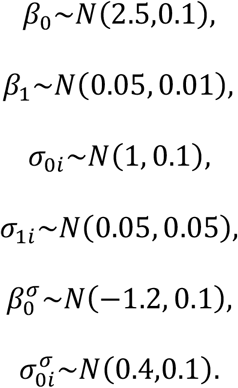

Simulation-based calibration for model 2.2 showed that the rank statistics were approximately uniformly distributed (Appendix S3: Fig. S6), and that the coverage of parameters was reasonable (Appendix S3: Fig. S7); known parameters were approximately recovered (Appendix S3: Fig. S8).

**Figure S6:**
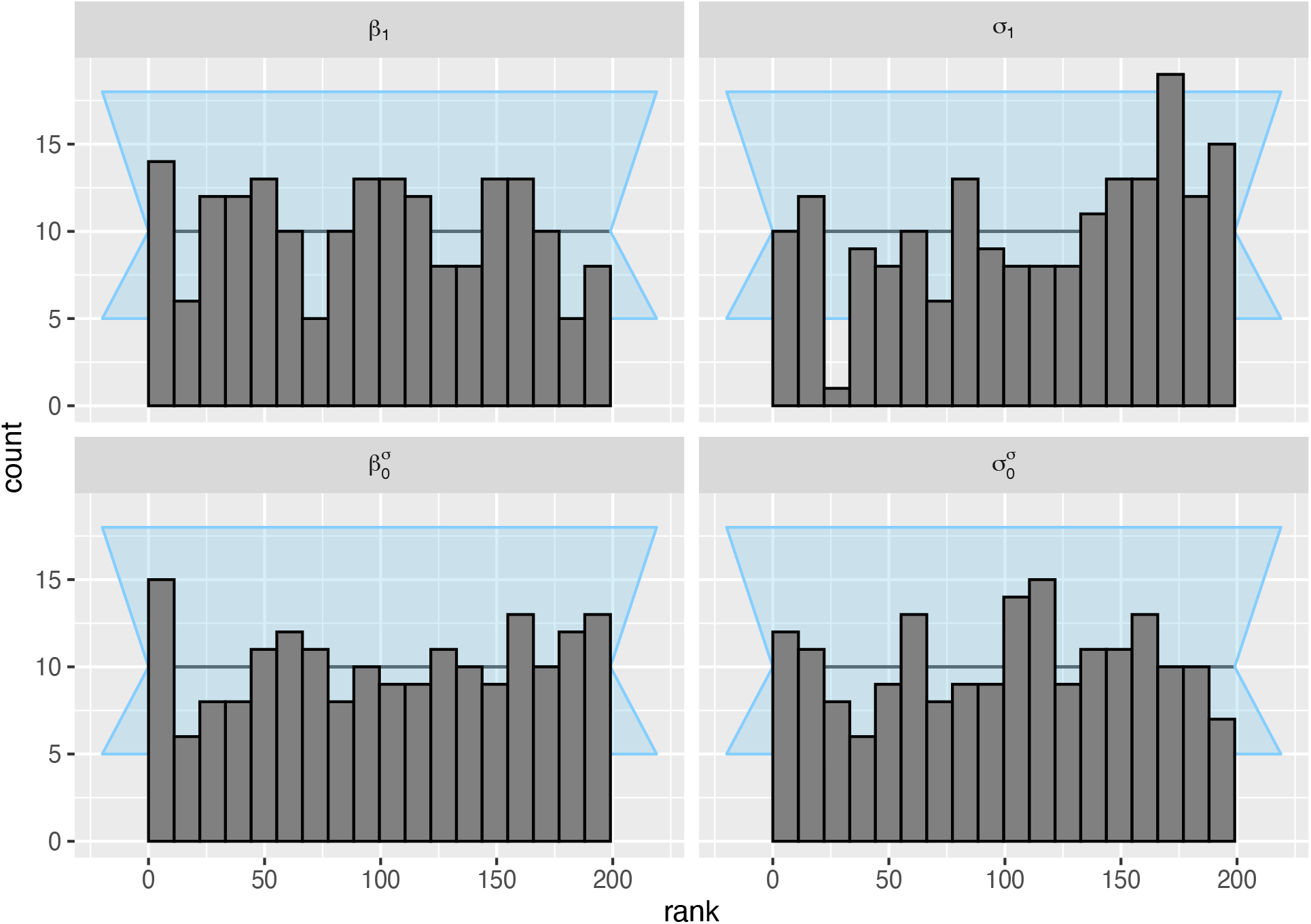
The posterior ranks of the prior draws were approximately normally distributed for the parameters of interest in model S2.2. Results are shown for n = 185 simulated data sets.Background (light blue shading) shows an approximate 95% interval for expected deviations.

**Figure S7:**
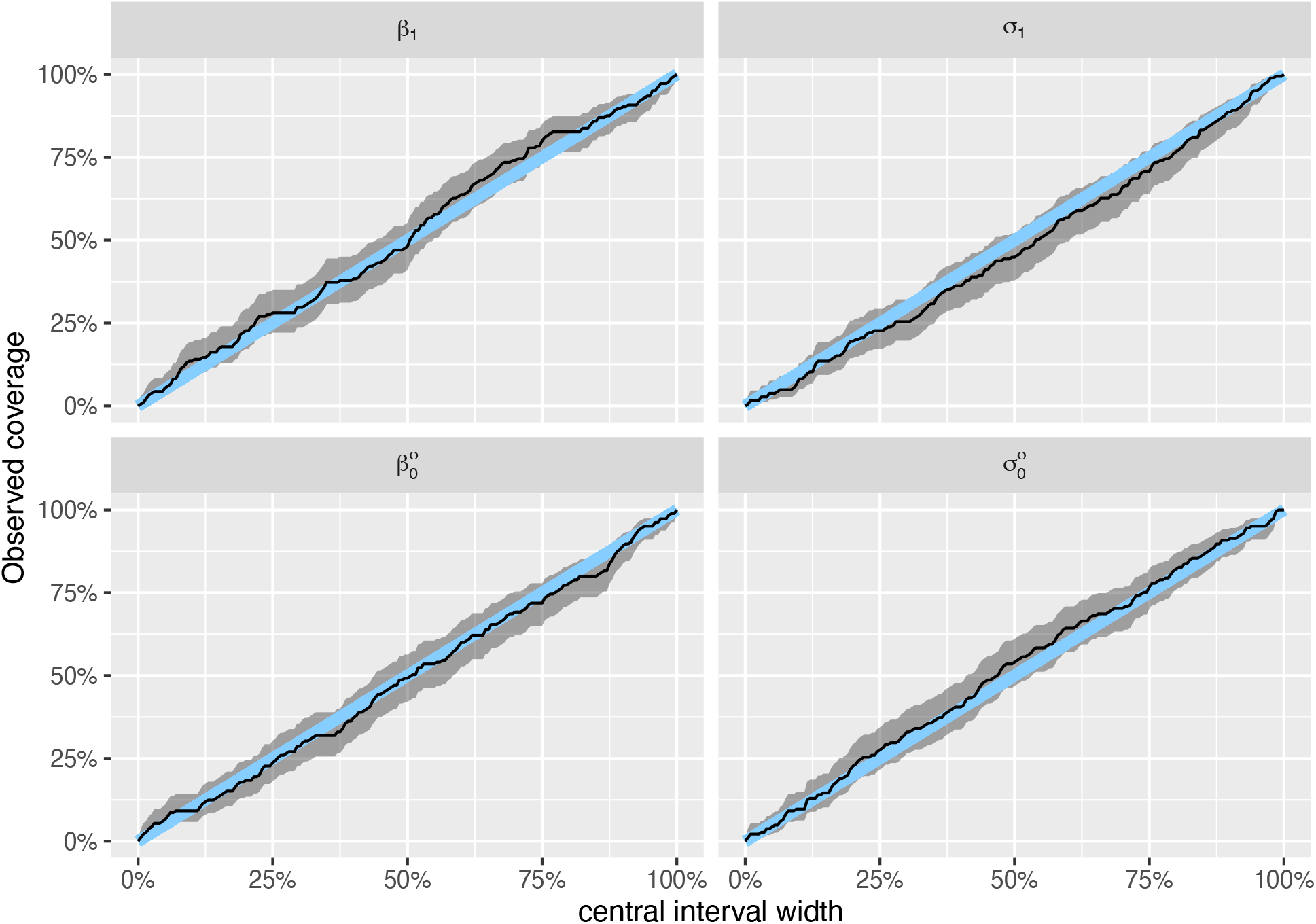
Model S2.2 had good coverage for the parameters of interest. Results are shown for n = 185 simulated data sets. Blue line is 1:1 line, and shading shows 95% uncertainty interval for the coverage.

**Figure S8:**
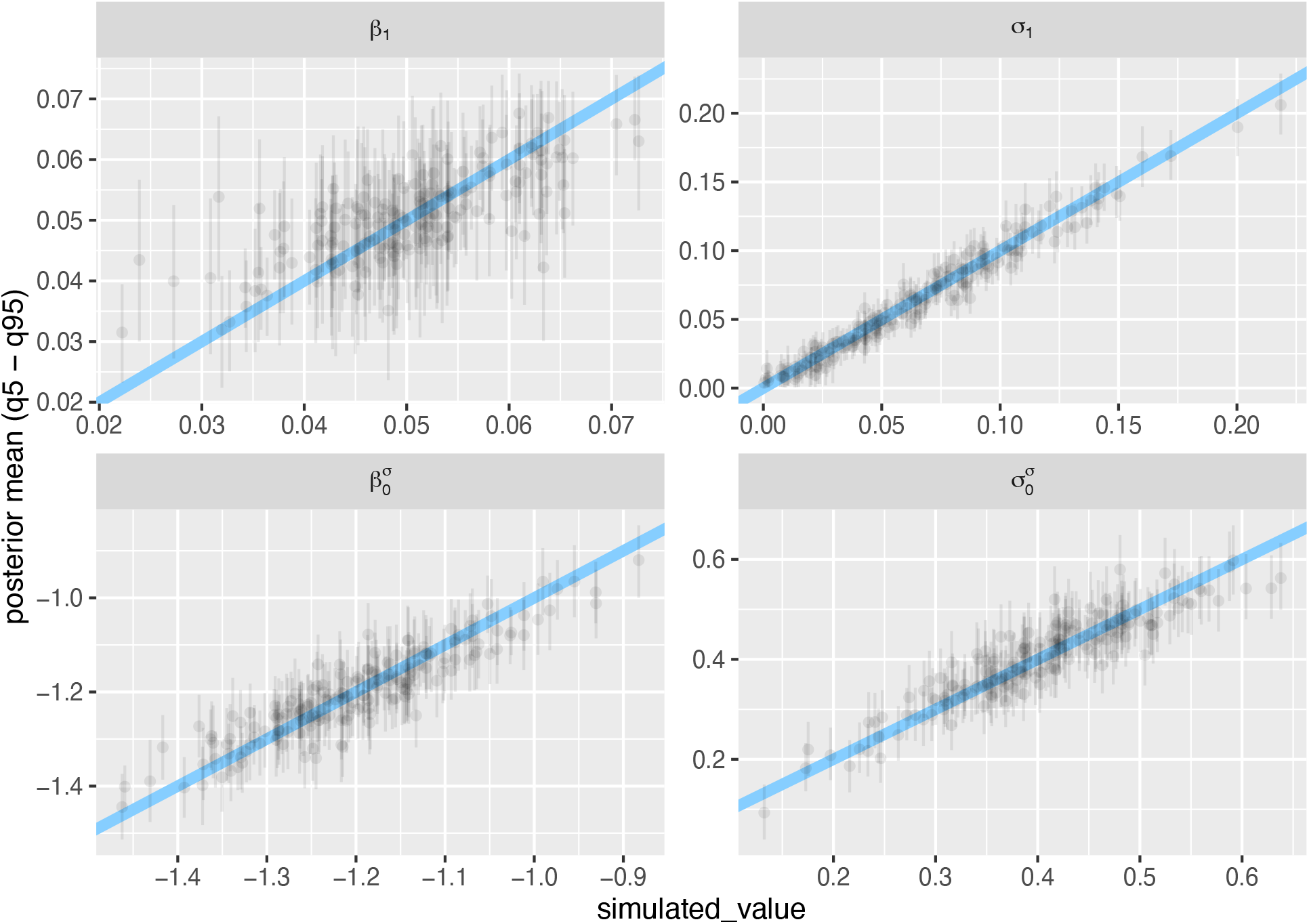
Model S2.2 was able to recover known parameter values fairly accurately, though with relatively low precision (with the exception of *σ*_1_). Results are shown for n = 185 simulated data sets; “simulated_value” (x-axis) is the known value of the parameter for a given simulation. Each point shows a parameter estimate, whiskers show 95% credible interval; diagonal line is the 1:1 line.

#### Model 2.3

Next, I extend this model to include (log) fragment size as a predictor of residual variation, and again allow it to varying between studies independently of varying study-level parameters for the mean:

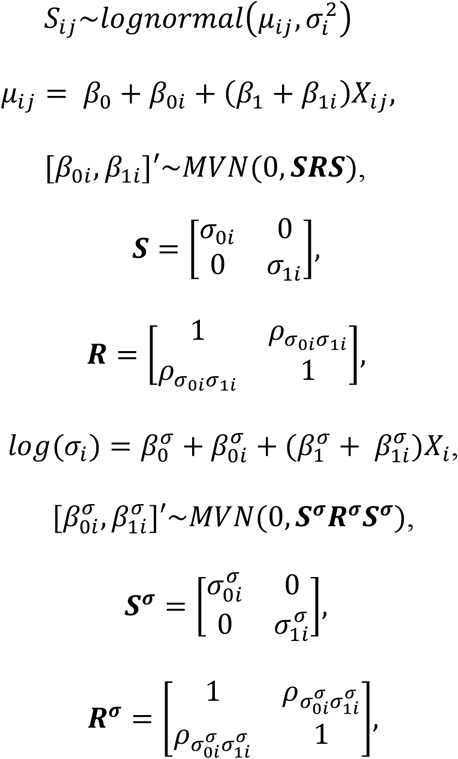

where 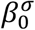 is the overall average residual variation, and 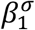 is the overall average slope of residual variation with fragment size; 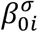 and 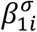 are the varying study-level departures from the intercept and slope, respectively, and were drawn from a multivariate normal distribution with zero mean and standard deviation 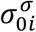 and 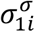, with correlations estimated in matrix ***R***^***σ***^. I fit the model with weakly regularizing priors:

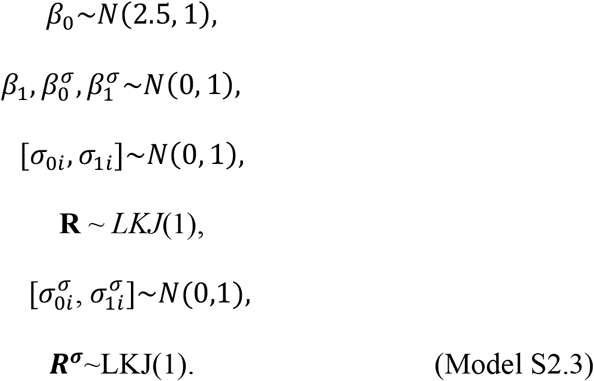

This model had good convergence (all Rhat <= 1.02), and showed a reasonable fit to the empirical data (Appendix S3: Fig. S9).

**Figure S9:**
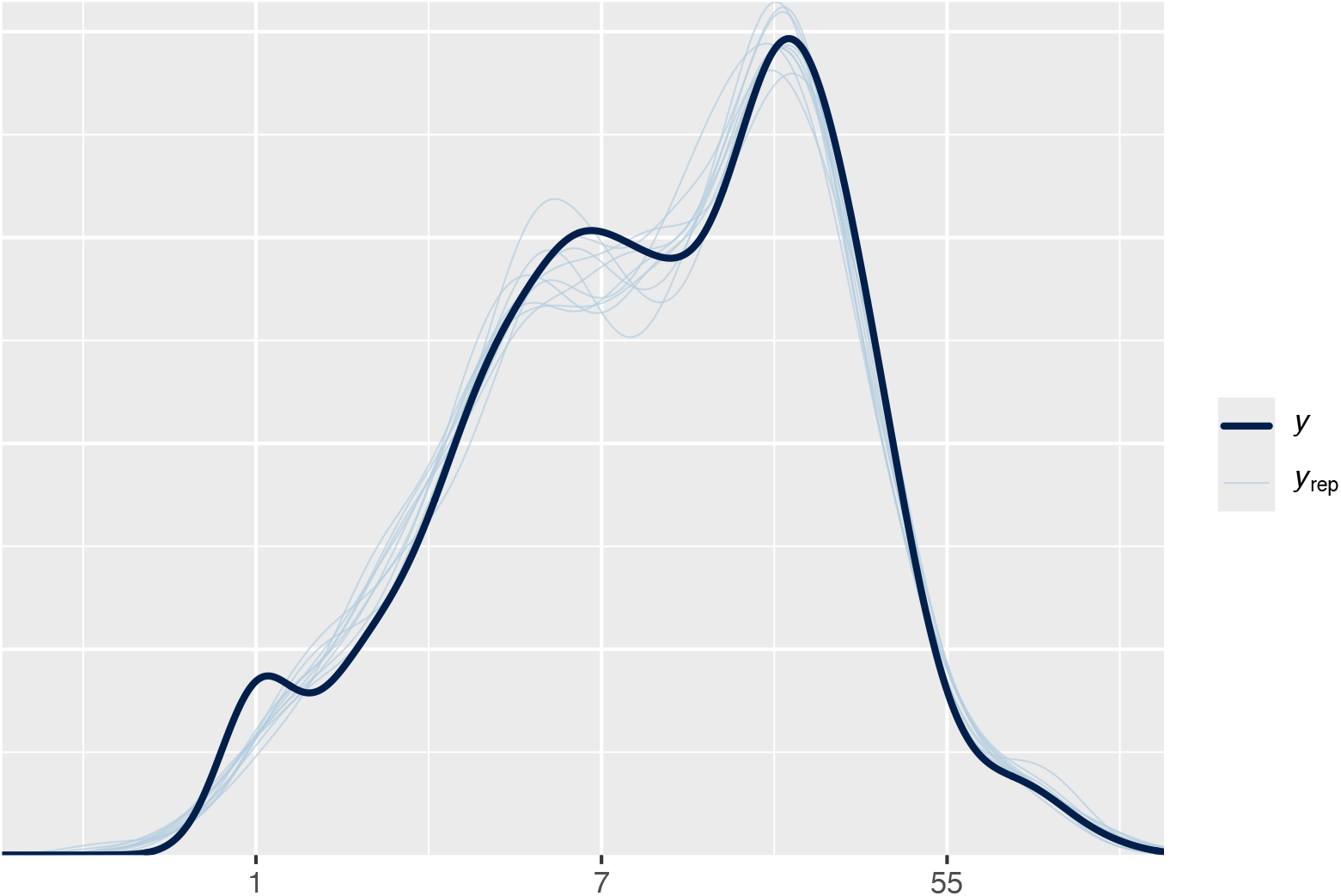
Posterior predictions from model 2.3 showed good fidelity to the observed data.

The parameter estimates from the fit of model S2.3 to empirical data (β_0_: 2.48 [95% credible interval: 2.3 – 2.66]; β_1_: 0.05 [95% credible interval: 0.03 – 0.06]; *σ*_0i_: 1.01 [95% credible interval: 0.89 – 1.14]; *σ*_1i_: 0.06 [95% credible interval: 0.04 – 0.07]; 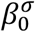 : -1.25 [95% credible interval: -1.35 – -1.15]; 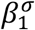 : -0.06 [95% credible interval: -0.09 *–* -0.03]; 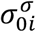 : 0.45 [95% credible interval: 0.37 – 0.55], and 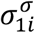 : 0.06 [95% credible interval: 0.01 *–* 0.1]) were used to inform the following priors:

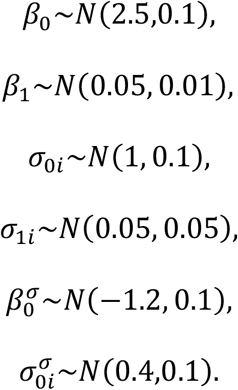

Simulation-based calibration for model 2.3 showed that the rank statistics were approximately uniformly distributed (Appendix S3: Fig. S10), and that the coverage of parameters was reasonable (Appendix S3: Fig. S11); known parameters were approximately recovered (Appendix S3: Fig. S12).

**Figure S10:**
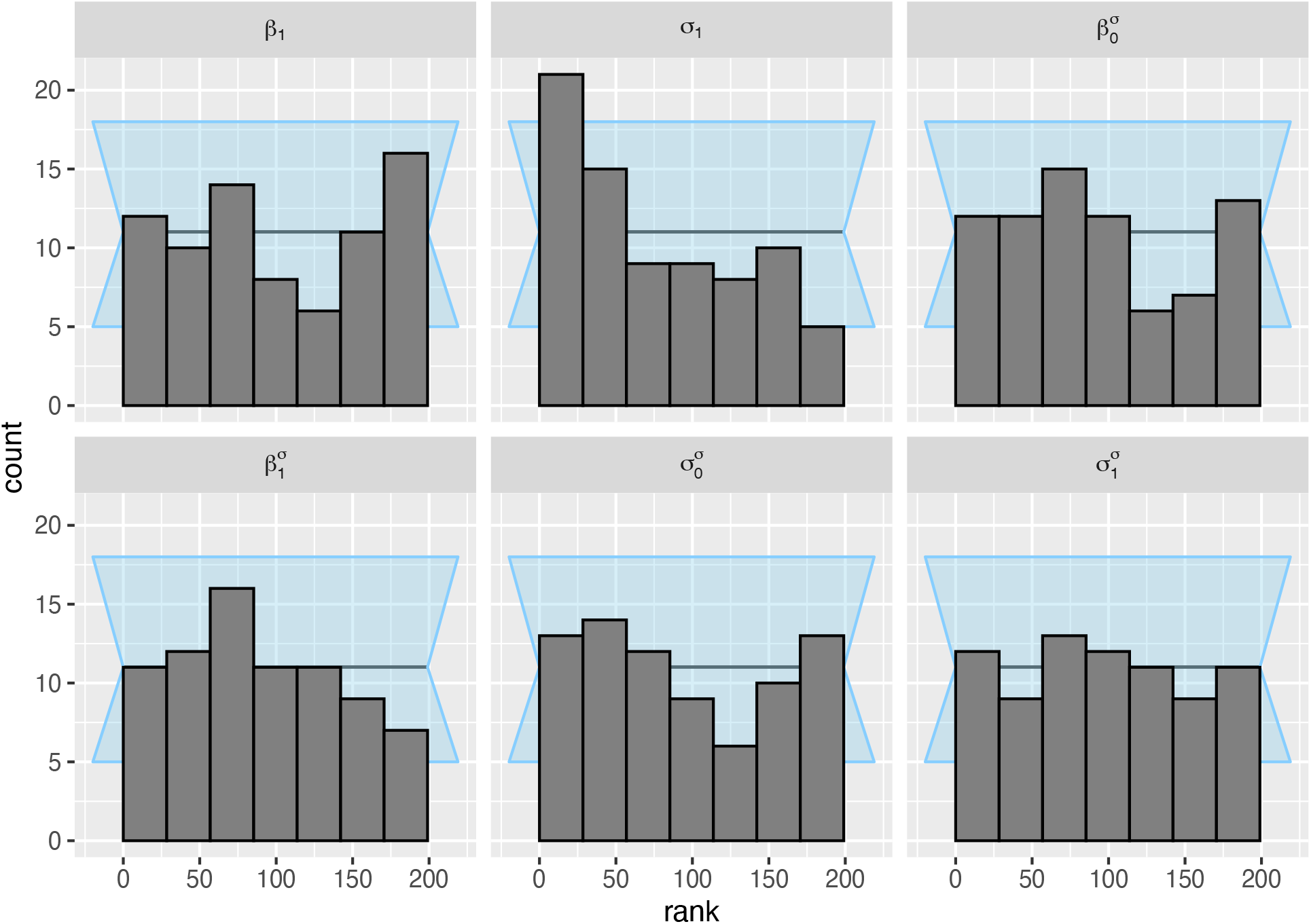
The posterior ranks of the prior draws were approximately normally distributed for the parameters of interest in model S2.3. Results are shown for n = 77 simulated data sets. Background (light blue shading) shows an approximate 95% interval for expected deviations.

**Figure S11:**
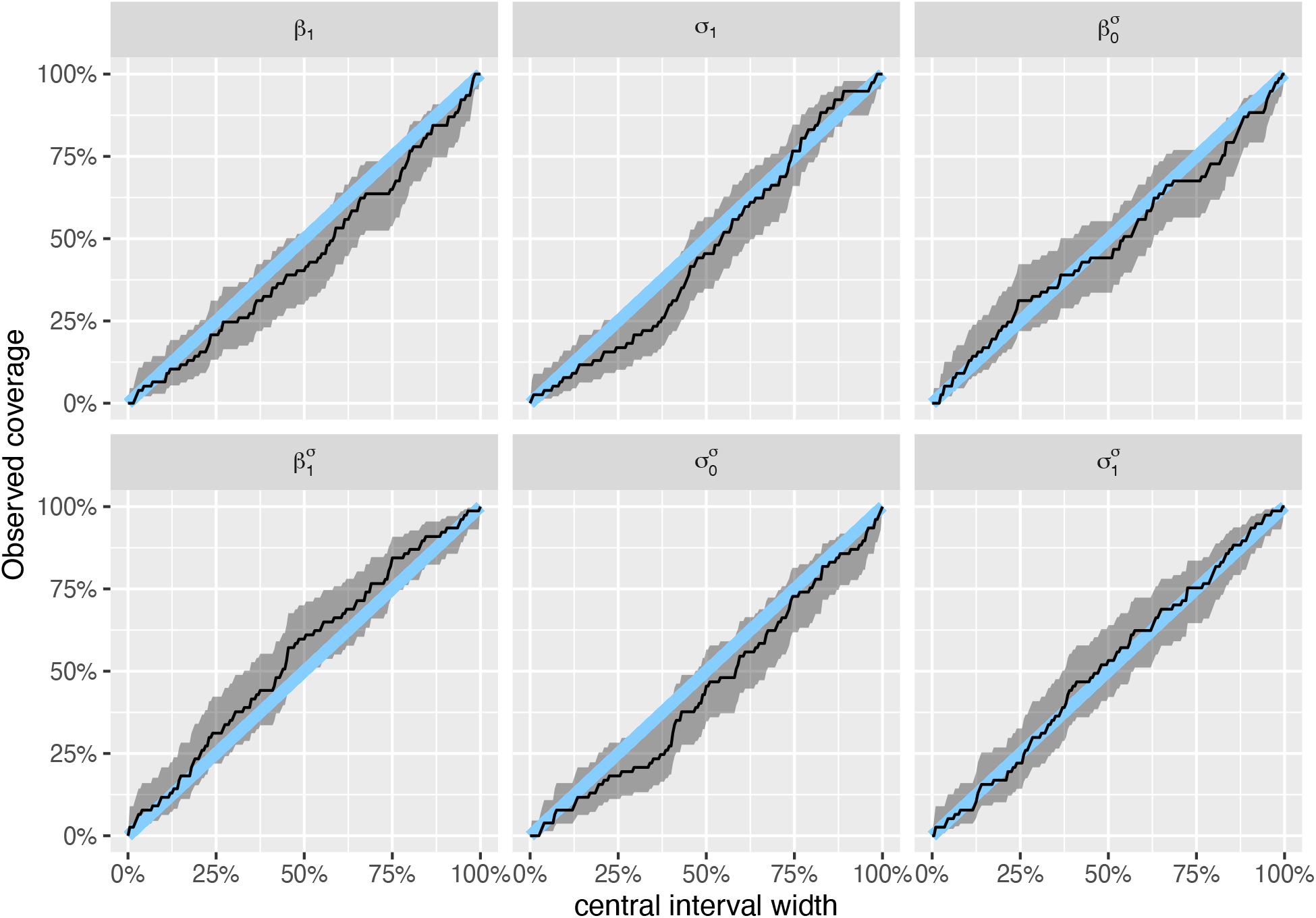
Model S2.3 had good coverage for all parameters of interest. Results are shown for n = 77 simulated data sets. Blue line is 1:1 line, and shading shows 95% uncertainty interval for the coverage.

**Figure S12:**
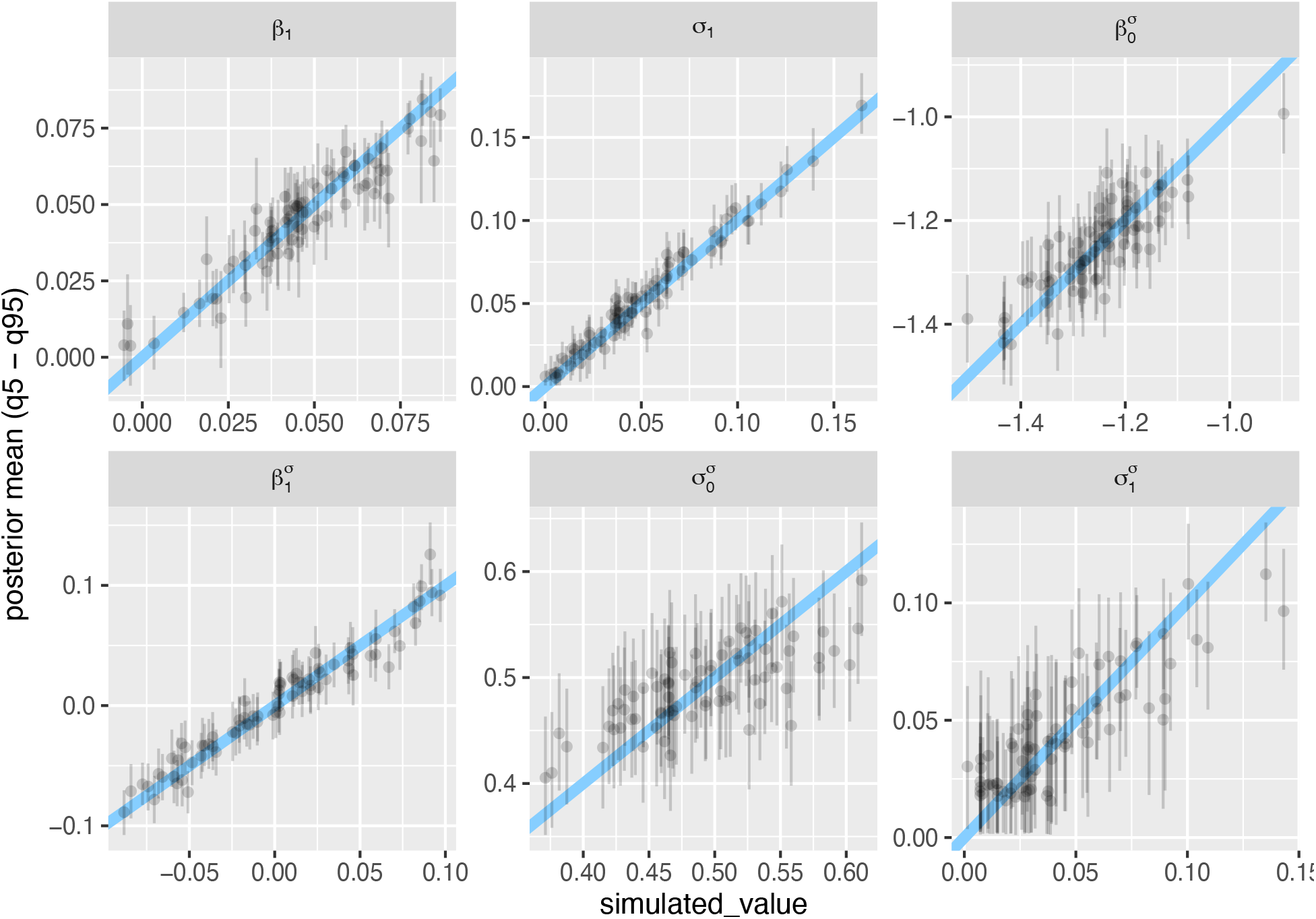
Model S2.3 was able to recover most parameters with reasonable accuracy, albeit with low precision (especially for 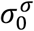 and 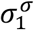). Results are shown for n = 77 simulated data sets; “simulated_value” (x-axis) is the known value of the parameter for a given simulation. Each point shows a parameter estimate, whiskers show 95% credible interval; diagonal line is the 1:1 line.

#### Model 2.4

The next model is the same as model 2.2 (i.e., varying study-level residuals), but allows the study-level residuals to covary with the other varying parameters for the mean:

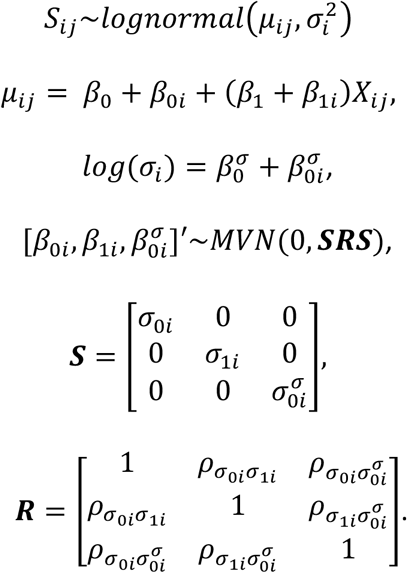

The model was fit to empirical data with weakly regularizing priors:

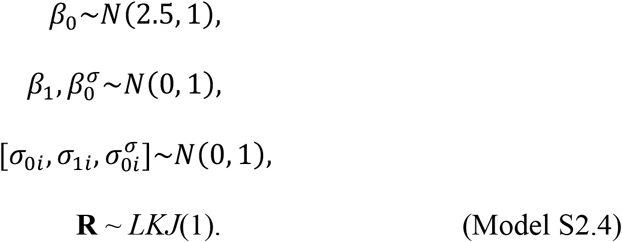

This model had good convergence (all Rhat <= 1.02), and showed a reasonable fit to the empirical data (Appendix S3: Fig. Sx).

**Figure S13:**
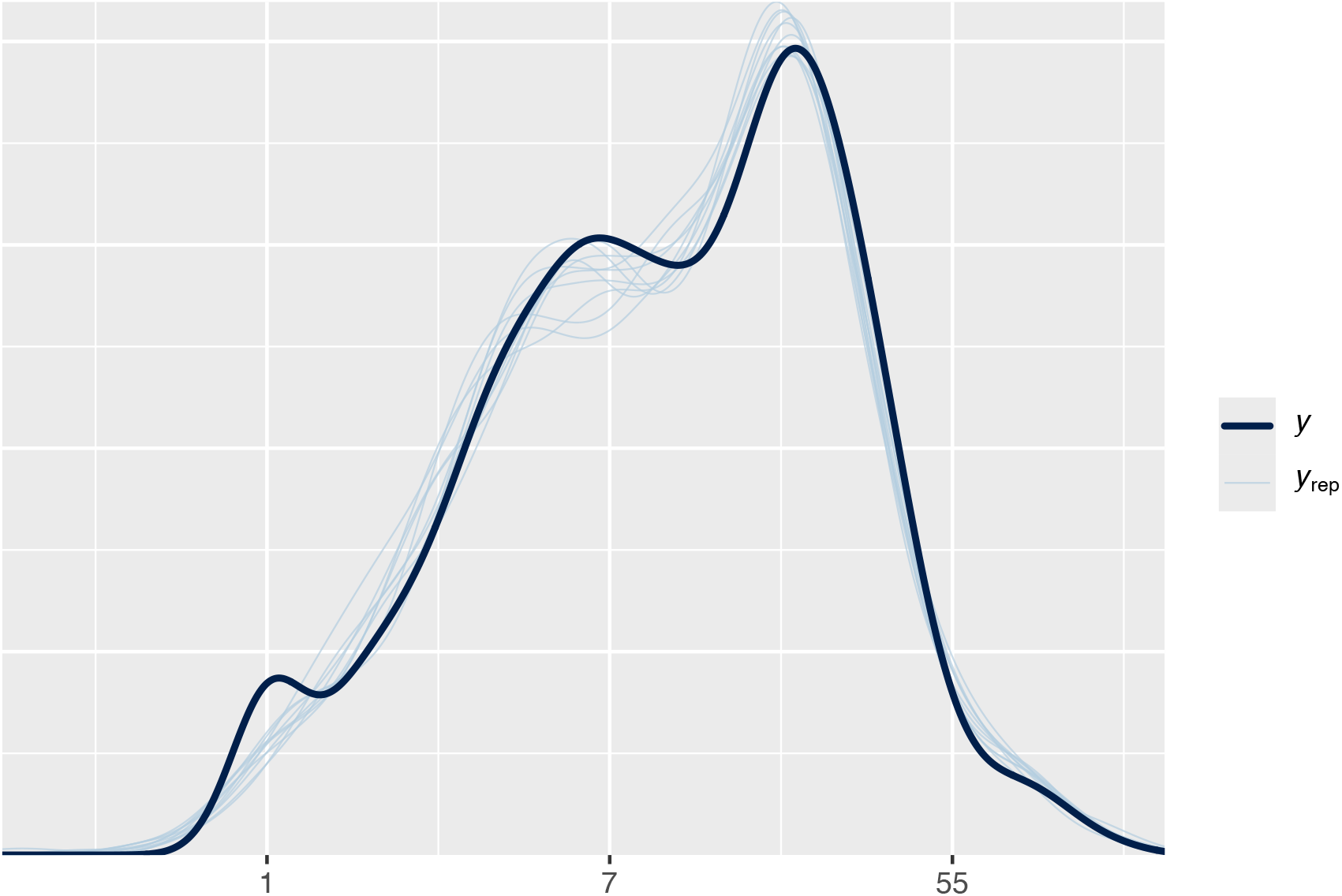
Posterior predictions showed that model 2.4 was able to make predict the observed data well.

The parameter estimates from the fit of model S2.4 to empirical data (β_0_: 2.46 [95% credible interval: 2.29 – 2.66]; β_1_: 0.05 [95% credible interval: 0.04 – 0.07]; *σ*_0i_: 1 [95% credible interval: 0.89 – 1.13]; *σ*_1i_: 0.06 [95% credible interval: 0.04 – 0.07], 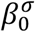 : -1.24 [95% credible interval: -1.35 – -1.15], and 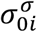 : 0.44 [95% credible interval: 0.37 *–* 0.53]) were used to inform the following priors:

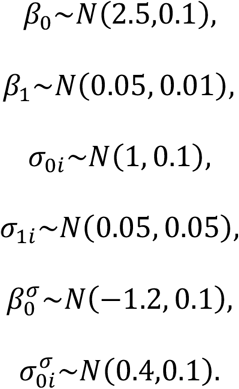

Simulation-based calibration for model 2.4 showed that the rank statistics were approximately uniformly distributed (Appendix S3: Fig. S14), and that the coverage of parameters was reasonable (Appendix S3: Fig. S15); known parameters were approximately recovered (Appendix S3: Fig. S16).

**Figure S14:**
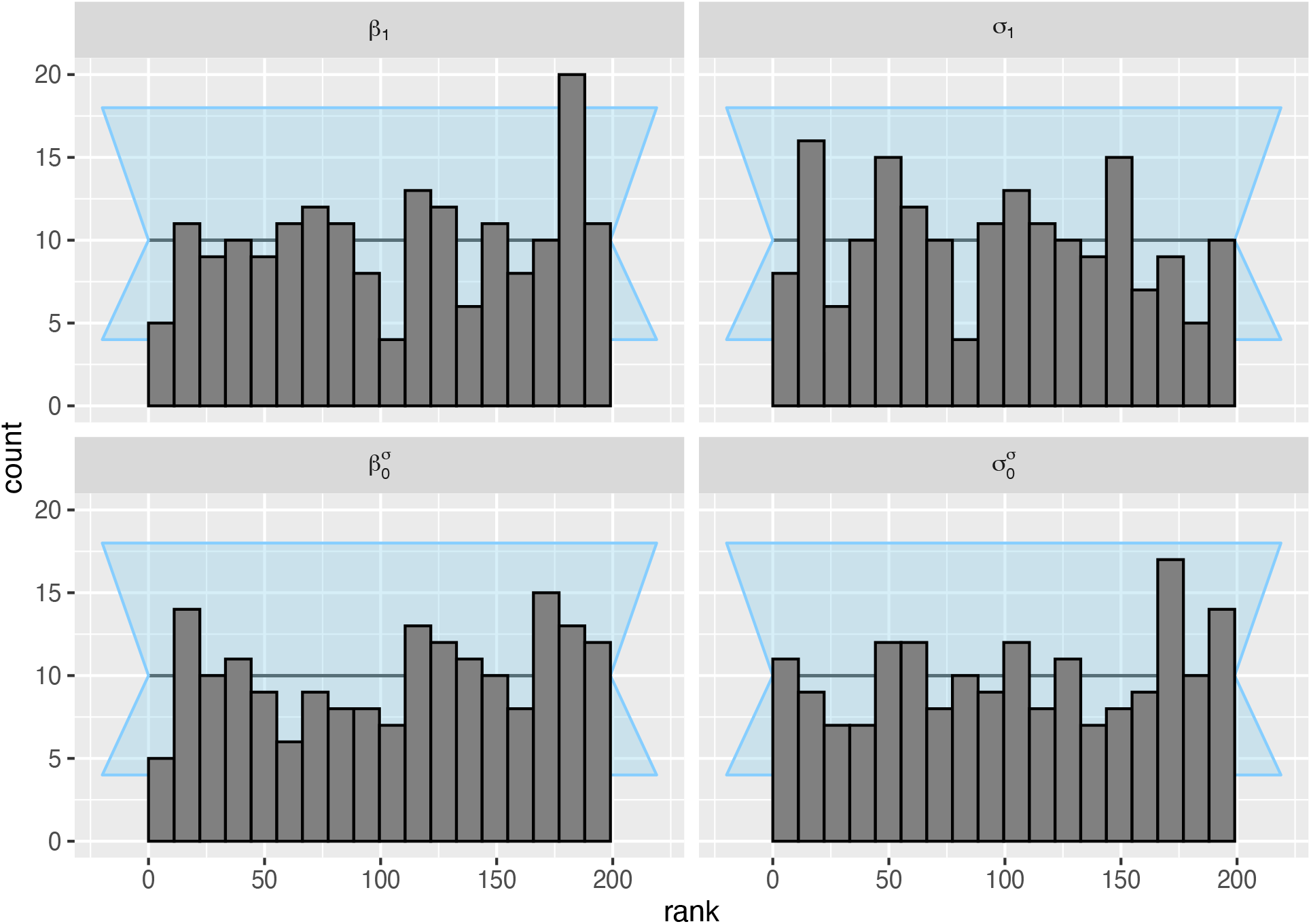
The posterior ranks of the prior draws were approximately normally distributed for the parameters of interest in model S2.4. Results are shown for n = 181 simulated data sets. Background (light blue shading) shows an approximate 95% interval for expected deviations.

**Figure S15:**
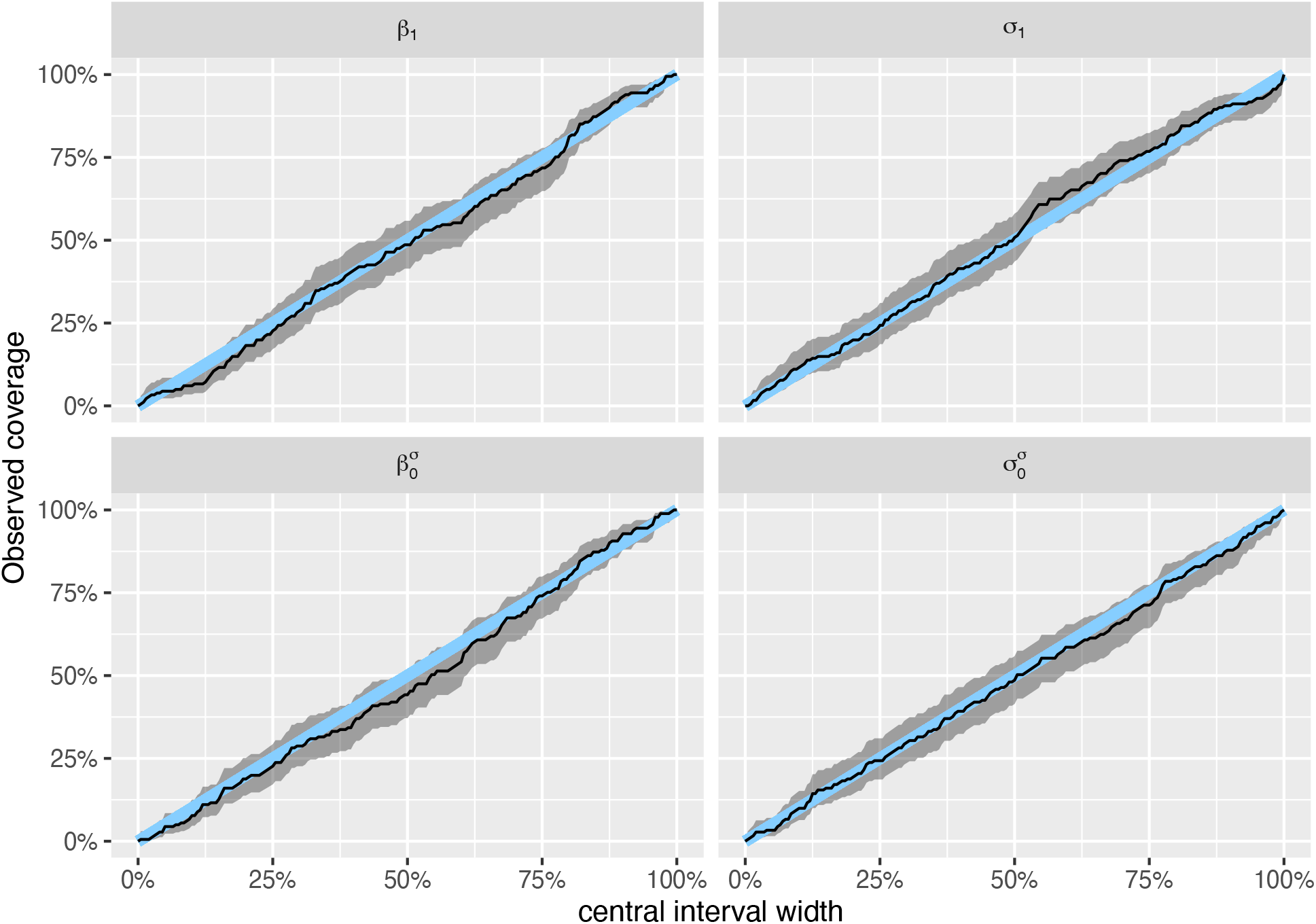
Model S2.4 had good coverage for all parameters of interest. Results are shown for n = 181 simulated data sets. Blue line is 1:1 line, and shading shows 95% uncertainty interval for the coverage.

**Figure S16:**
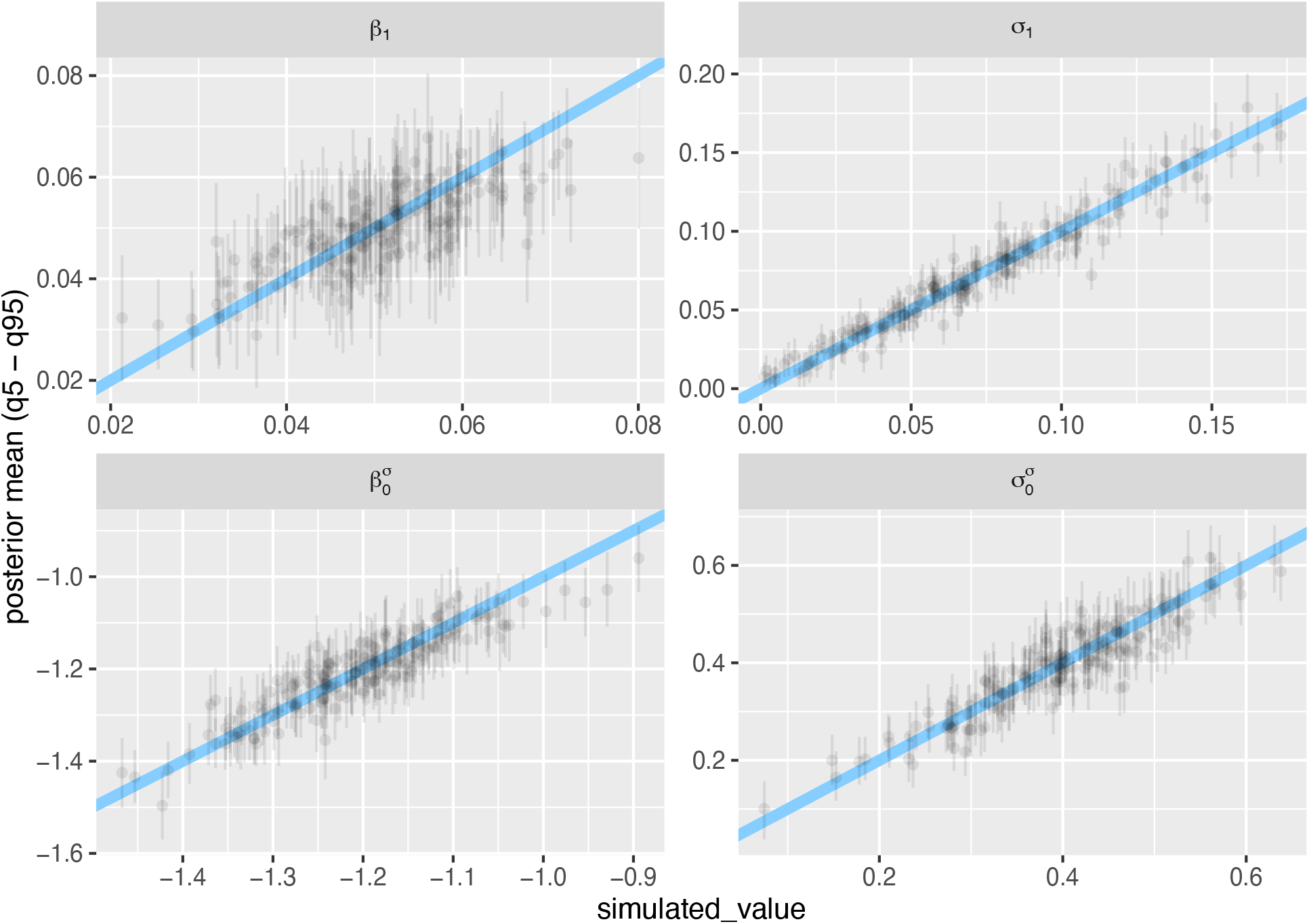
Model S2.4 was able to recover known parameter values with reasonable accuracy. Results are shown for n = 181 simulated data sets; “simulated_value” (x-axis) is the known value of the parameter for a given simulation. Each point shows a parameter estimate, whiskers show 95% credible interval; diagonal line is the 1:1 line.

#### Model 2.5

The final model extended model 2.3 to allow for correlations between all of the varying study-level parameters (i.e., for the location and the scale):

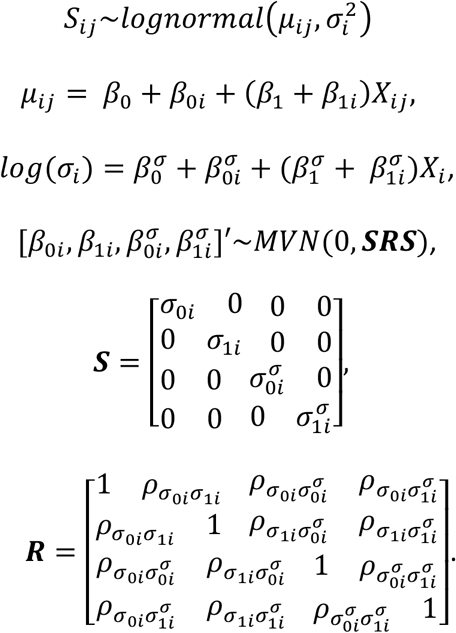

I fit the model with weakly regularizing priors:

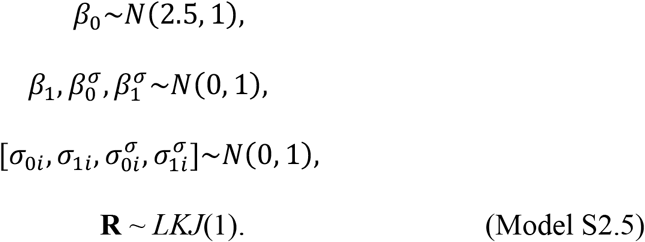

This model had good convergence (all Rhat < 1.01), and showed a reasonable fit to the empirical data (Appendix S3: Fig. S17).

**Figure S17:**
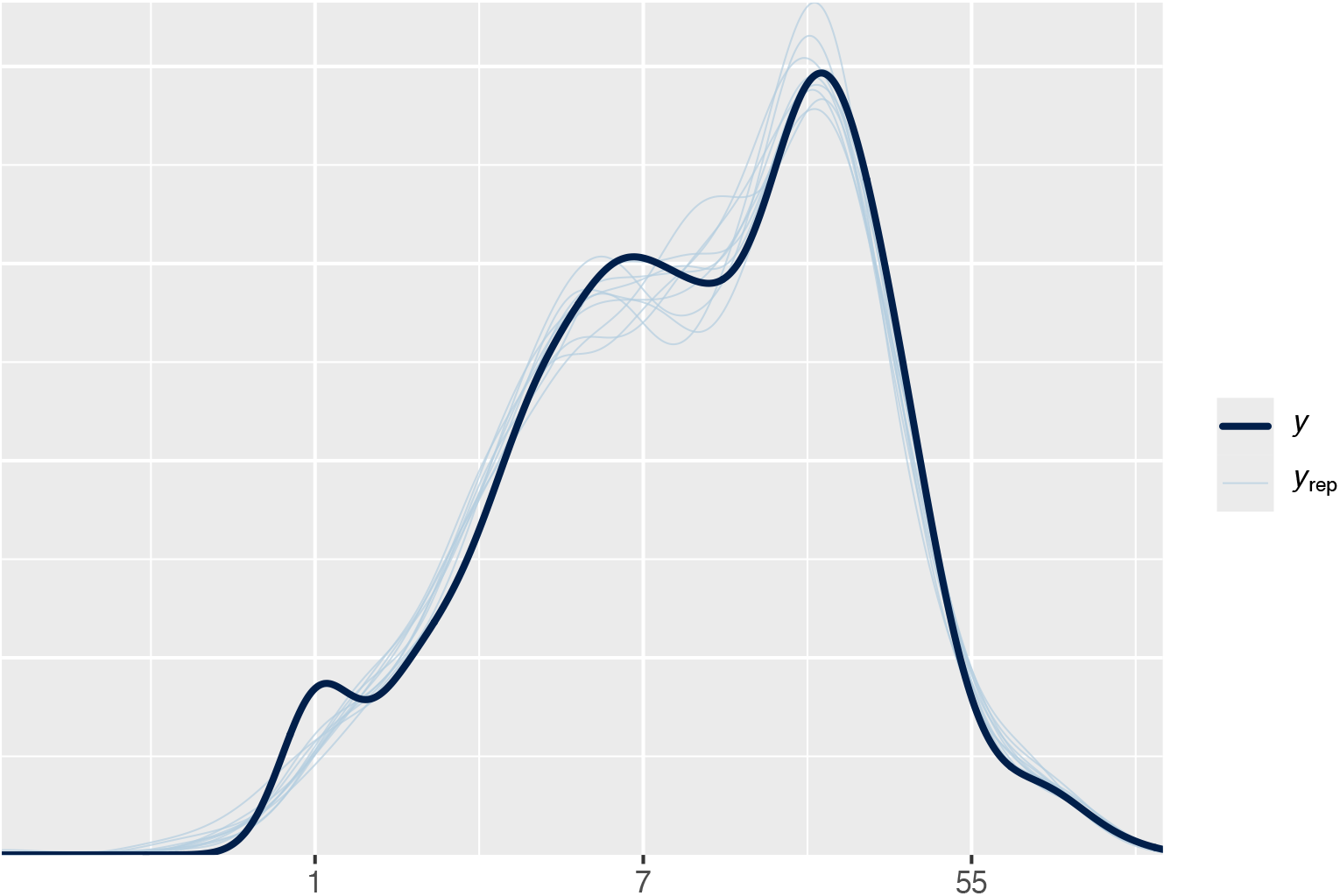
Model 2.5 made predictions that resembled the observed data closely.

The parameter estimates from the fit of model S2.5 to empirical data (β_0_: 2.47 [95% credible interval: 2.3 – 2.66]; β_1_: 0.05 [95% credible interval: 0.03 – 0.06]; *σ*_0i_: 1.01 [95% credible interval: 0.89 – 1.15]; *σ*_1i_: 0.06 [95% credible interval: 0.04 – 0.07]; 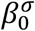 : -1.28 [95% credible interval: -1.39 – -1.18]; 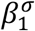 : -0.06 [95% credible interval: -0.09 *–* -0.04]; 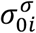 : 0.46 [95% credible interval: 0.39 – 0.56], and 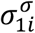 : 0.07 [95% credible interval: 0.03 *–* 0.11]) were used to inform the following priors:

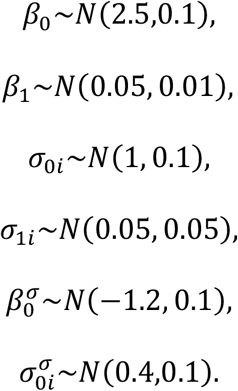

Simulation-based calibration for model 2.5 showed that the rank statistics were approximately uniformly distributed (Appendix S3: Fig. S18), and that the coverage of parameters was reasonable (Appendix S3: Fig. S19); known parameters were approximately recovered (Appendix S3: Fig. S20).

**Figure S18:**
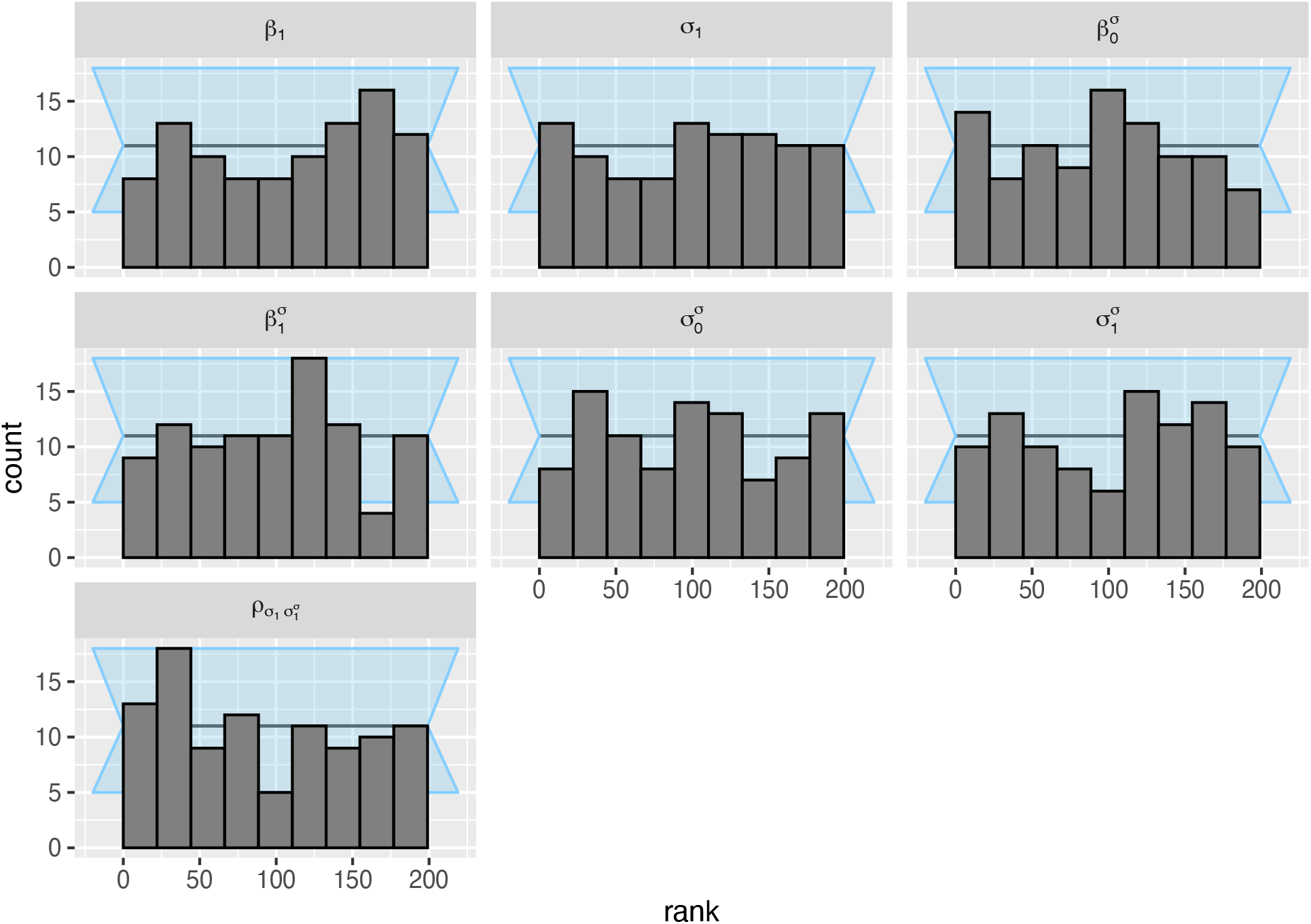
The posterior ranks of the prior draws were approximately normally distributed for the parameters of interest in model S2.5. Results are shown for n = 98 simulated data sets.

**Figure S19:**
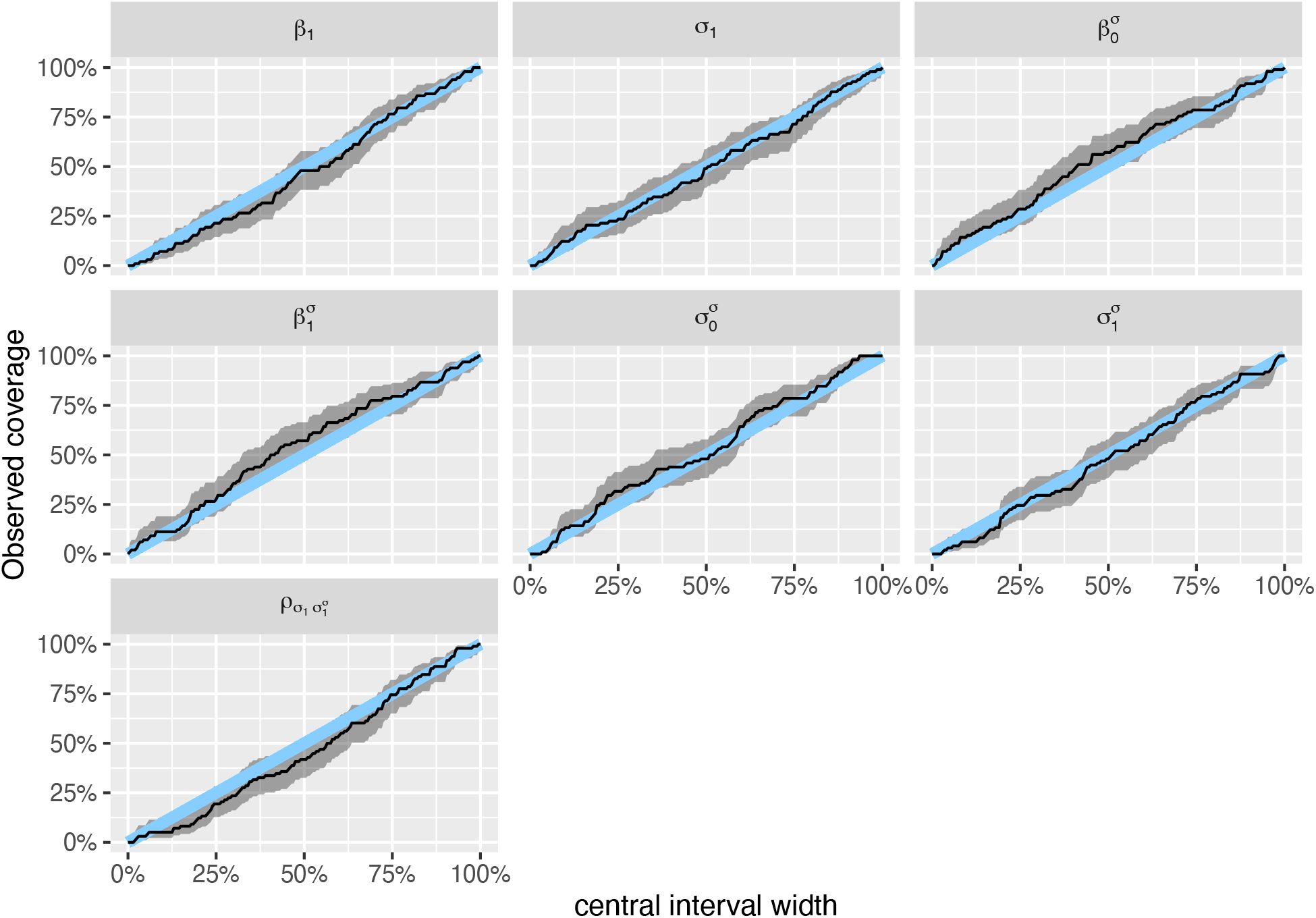
Model S2.5 had good coverage for all parameters of interest. Results are shown for n = 98 simulated data sets. Blue line is 1:1 line, and shading shows 95% uncertainty interval for the coverage.

**Figure S20:**
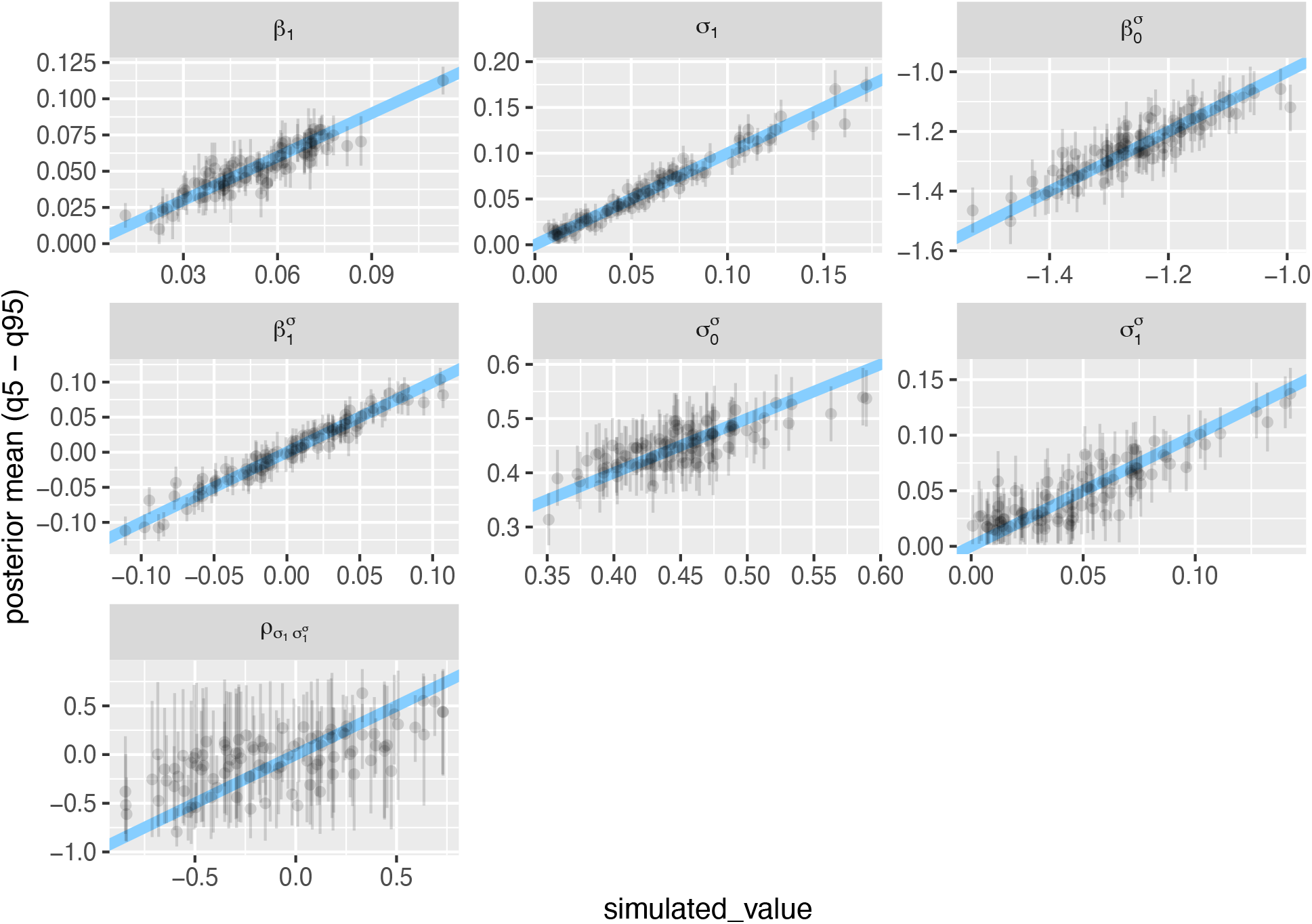
Model S2.5 was able to recover known parameter values with reasonable accuracy, with the exception of the correlation 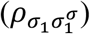, which was estimated with high uncertainty (low precision). Results are shown for n = 98 simulated data sets; “simulated_value” (x-axis) is the known value of the parameter for a given simulation. Each point shows a parameter estimate, whiskers show 95% credible interval; diagonal line is the 1:1 line.

**Figure S21:**
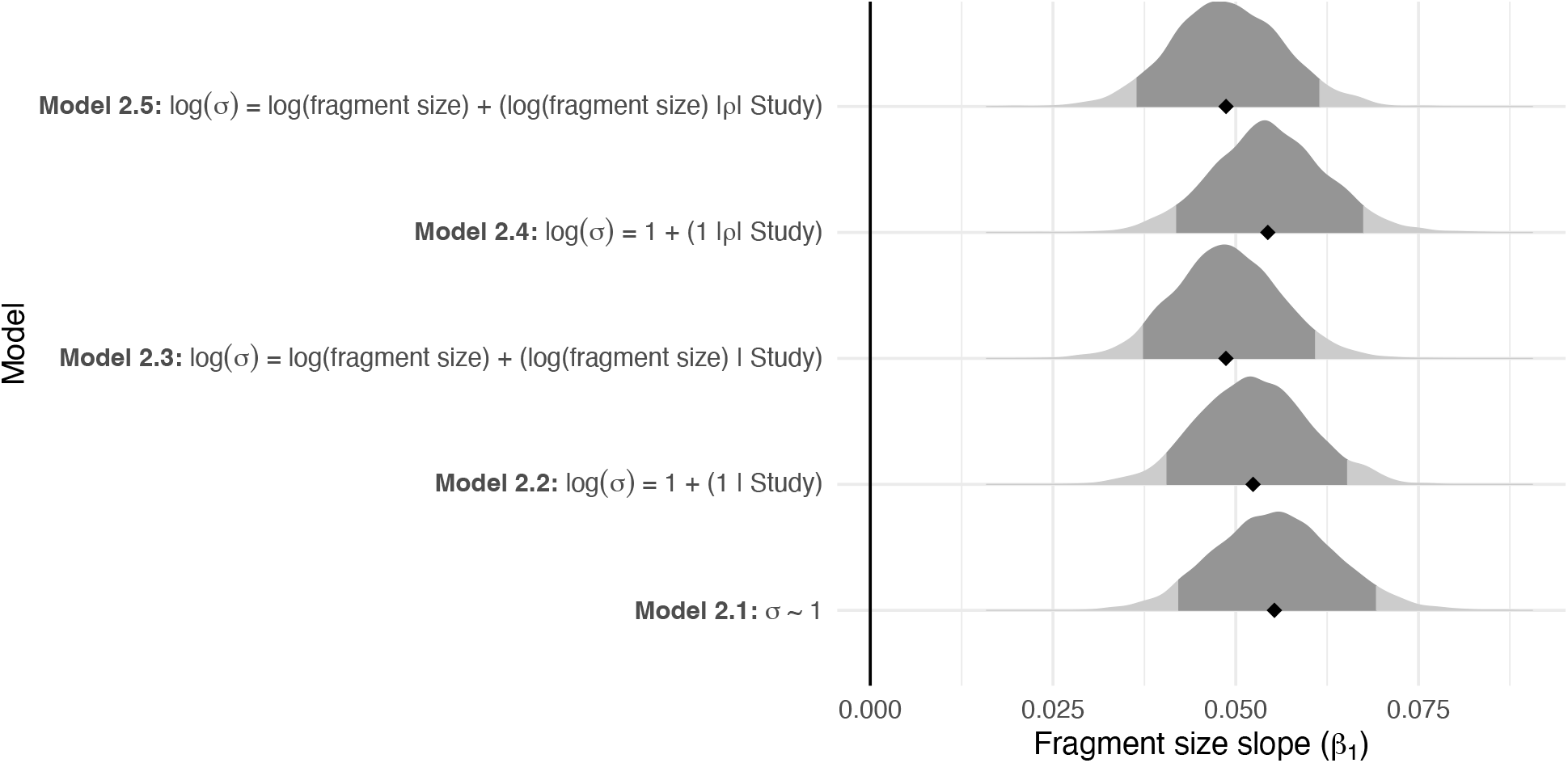
The different models for heteroscedasticity did not qualitatively change the support for the ecosystem decay hypothesis (i.e., posterior distributions of the slope parameter for the location [*β*_1_] were greater than zero).

**Figure S22:**
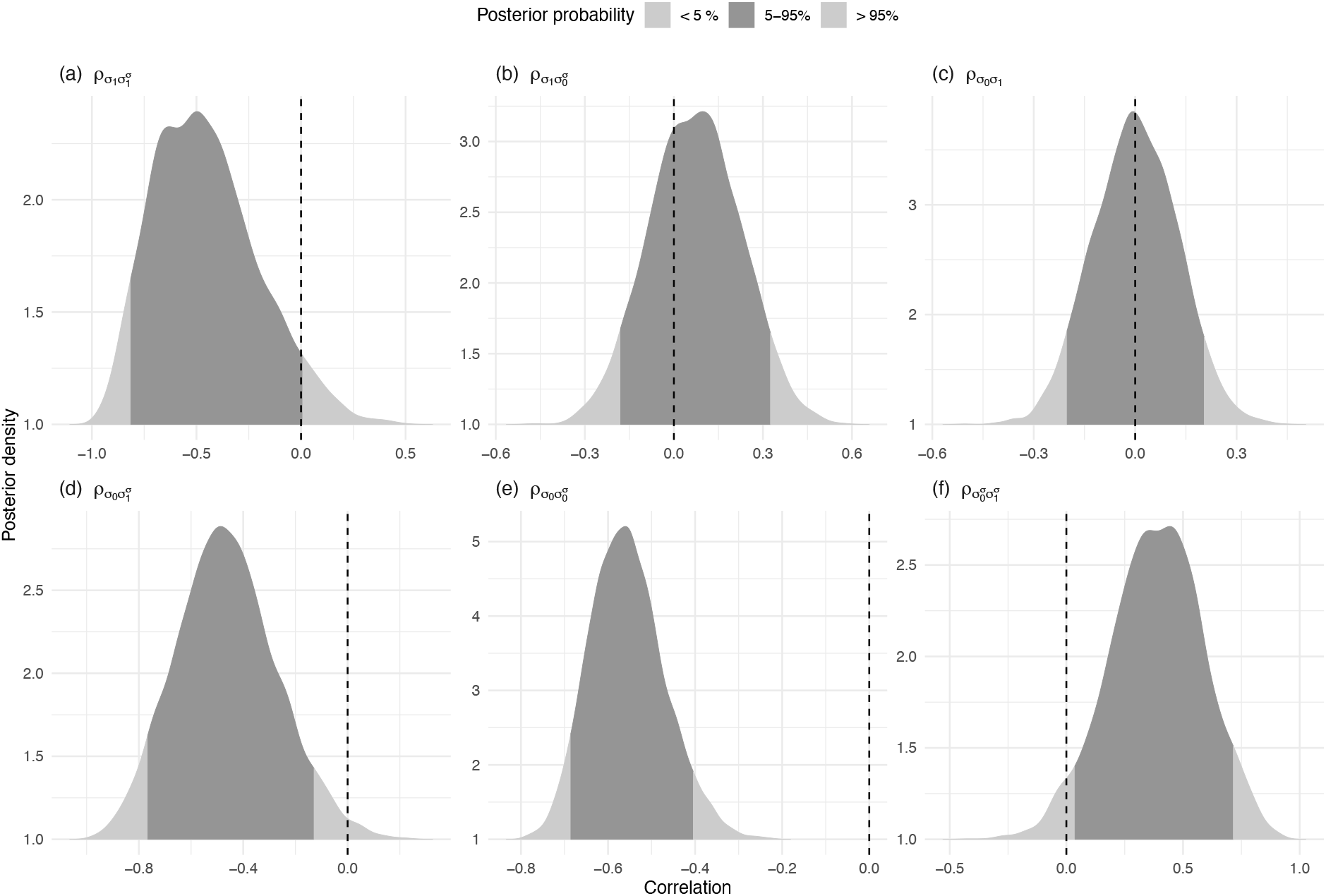
Posterior distributions (density plots for 1000 posterior draws) showing correlations among the varying study-level parameters for model 2.5.

**Figure S22:**
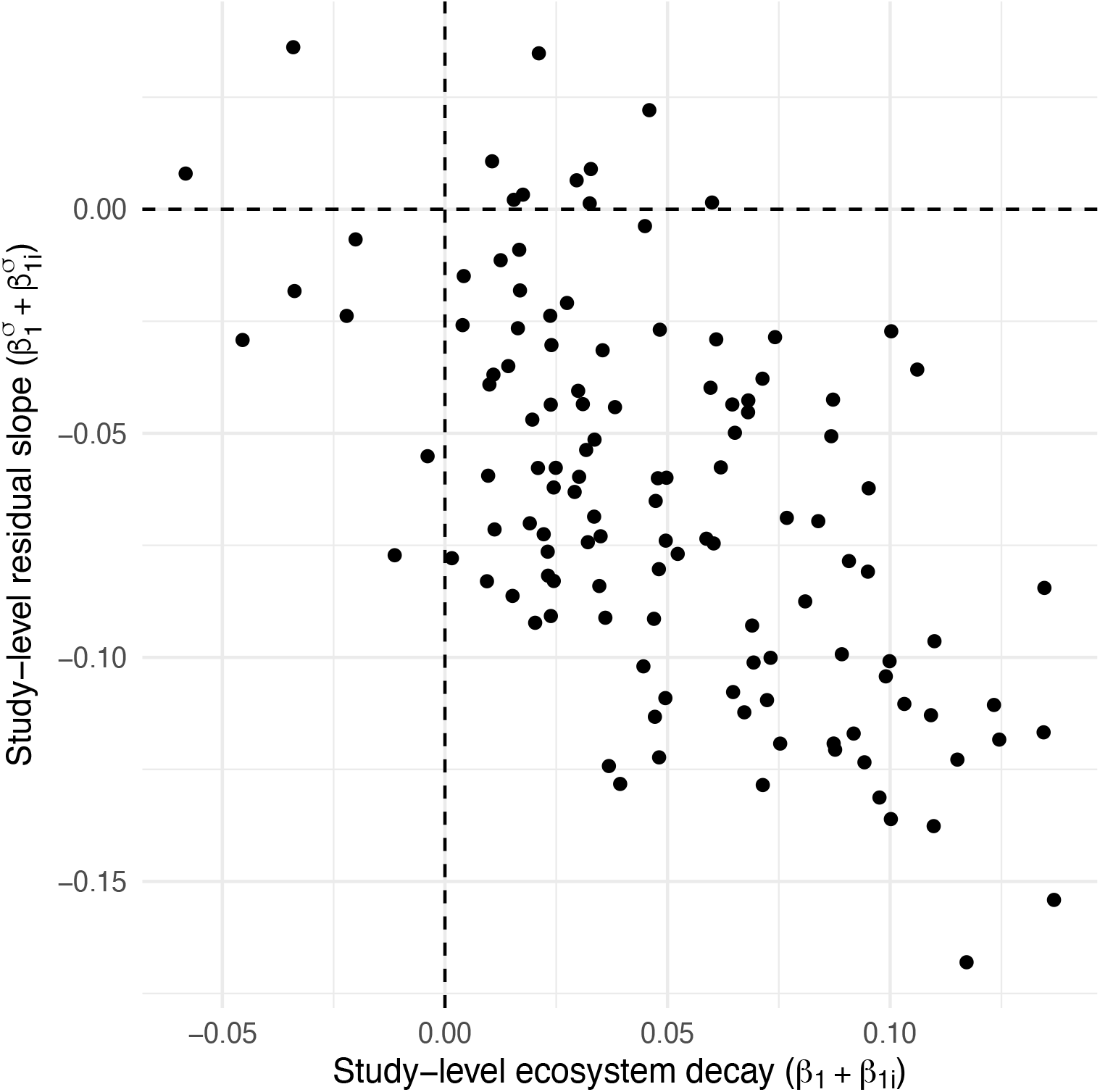
The decline of residual variation (scale) with increasing patch size was greatest for studies with the strongest ecosystem decay. Each point shows the estimate of a study-level slope, with slopes for the location on the x-axis, and the scale on the y-axis.

